# H3K27me3-rich genomic regions can function as silencers to repress gene expression via chromatin interactions

**DOI:** 10.1101/684712

**Authors:** Yichao Cai, Ying Zhang, Yan Ping Loh, Jia Qi Tng, Mei Chee Lim, Zhendong Cao, Anandhkumar Raju, Erez Lieberman-Aiden, Shang Li, Lakshmanan Manikandan, Vinay Tergaonkar, Greg Tucker-Kellogg, Melissa Jane Fullwood

## Abstract

Gene repression and silencers are poorly understood. We reasoned that H3K27me3-rich regions (MRRs) of the genome defined from clusters of H3K27me3 peaks may be used to identify silencers that can regulate gene expression via proximity or looping. MRRs were associated with chromatin interactions and interact preferentially with each other. MRR component removal at interaction anchors by CRISPR led to upregulation of interacting target genes, altered H3K27me3 and H3K27ac levels at interacting regions, and altered chromatin interactions. Chromatin interactions did not change at regions with high H3K27me3, but regions with low H3K27me3 and high H3K27ac levels showed changes in chromatin interactions. The MRR knockout cells also showed changes in phenotype associated with cell identity, and altered xenograft tumor growth. MRR-associated genes and long-range chromatin interactions were susceptible to H3K27me3 depletion. Our results characterized H3K27me3-rich regions and their mechanisms of functioning via looping.

## Introduction

The 3-dimensional organization of our genomes is important for gene regulation^1–3^. The genome is organized into large Topologically-Associated Domains (TADs) and chromatin interactions. Gene transcription is controlled by transcription factors (TFs) that bind to enhancers and promoters to regulate genes^4^. TFs can bind to proximal enhancers in the genome, and enhancers distal to genes can loop to gene promoters via chromatin interactions to activate gene expression^3^. Cancer cells show altered chromatin interactions^2,3^ including altered chromatin loops to key oncogenes such as *TERT*^5^.

By contrast, mechanisms for gene repression are much less well understood. Silencers are regions of the genome that are capable of silencing gene expression. Silencers have been shown to exist in the human genome, but are less well characterized than enhancers. Until now, there are only a few known experimentally validated silencers that have been demonstrated to repress target genes *in vitro*, such as the human synapsin I gene^6^, the human *BDNF* gene^7^ and human *CD4* gene^8,9^ (experimentally validated silencer examples are discussed in Table S1). The reason for the paucity of known silencers in the literature is that methods to identify human silencer elements in a genome-wide manner are only starting to be developed now. Moreover, the mechanism by which silencers can regulate distant genes is still uncharacterized. Distant silencers are thought to loop over to target genes to silence them^10,11^, and this has been demonstrated in studies of polycomb-mediated chromatin loops in *Drosophila*^12^ and in mice^13^, but no such examples have been characterized to date in humans.

Polycomb Group (PcG) proteins including Polycomb Repressive Complexes, PRC1 and PRC2 are widely recognized to mediate gene silencing of developmental genes^14^. During the development process, PRC1 and PRC2 have the ability to orchestrate genome architecture and repress gene expression^15^. There are two different types of genomic domains: active domains and repressive domains, which to regulate gene expression and construct cellular identity. Genes involved in cell self-renewal are contained within the active domains which are governed by super-enhancers, while genes specifying repressed lineage are organized within chromatin structures known as PcG domains^16^. Moreover, intact PcG domains have been shown to be necessary to maintain the chromatin interaction landscape^17,18^. However, the mechanisms of PcG domain formation and PcG proteins recruitment are not fully characterized yet^19^, which makes finding silencers more difficult.

PcG domains are marked by H3K27me3, which is deposited by the catalytic component of PRC2 complex, mainly Enhancer of zeste homolog 2 (EZH2) and sometimes EZH1^20^. H3K27me3 marks are associated with gene repression for cell type-specific genes. Unlike H3K9me3 which remains silenced all the time and prevents multiple TFs from binding^21^, H3K27me3 still allows these genes to be activated through TF binding in a different cell state^22^. H3K27me3 is known to be a characteristic of silencers^18,23^. Although large blocks of H3K27me3-marked loci have been observed in previous studies^24–26^, their regulatory actions and roles in chromatin loops were not explored in these manuscripts.

Recently, several studies have proposed methods to identity silencer elements in a genome-wide manner. Huang *et al* defined silencers using the correlation between H3K27me3-DNase I hypersensitive site (DHS) and gene expression^27^. At the same time, Jayavelu *et al* used a subtractive analysis approach to predict silencers in over 100 human and mouse cell types^28^. Moreover, Pang and Snyder identified silencers through an innovative “ReSE screen” which screened for genomic regions that can repress caspase 9 expression upon apoptosis induction^29^. Interestingly, Ngan *et al* characterized silencers in mouse development through PRC2 Chromatin Interaction Analysis with Paired-End Tag sequencing (ChIA-PET) in mouse embryonic stem cells. They concluded that PRC2-bound looping anchors function as transcriptional silencers suggesting that we can identify silencers through investigating chromatin interactions^13^.

However, there is no consensus yet in terms of how to identify silencers. Notably, each of these methods identify different genomic regions as silencers, raising the question of whether there may be different classes of silencers. Moreover, current methods for identifying silencers are laborious and require complicated bioinformatics analyses and/or genome-wide screening (Table S2, “comparison of different human silencer identification methods”). A simple, easy to perform method to identify silencers in the genome in a high-throughput manner would be ideal. Further investigation is needed to understand whether there are different classes of silencers and to characterize the roles of silencers in the genome.

The term “super-enhancer”^30^ has been used to describe clusters of H3K27ac peaks which show very high levels of H3K27ac or other transcription-associated factors such as mediators as determined from ChIP-Seq data. Super-enhancers have high levels of chromatin interactions to target genes^31^, and are associated with oncogenes in cancer cells^32^ and cell fate-associated genes in embryonic stem cells^33^. While more research needs to be done to determine if super-enhancers are a distinctly different entity from enhancers, super-enhancers are thought as strong enhancers, and the definition has been useful in identifying genes important for cell-type specification^34^.

Here, we reasoned that we can similarly identify “super-silencers” or “H3K27me3-rich regions (MRRs)” from clusters of H3K27me3 peaks in the genome through ChIP-Seq on H3K27me3. We hypothesized that H3K27me3-rich regions may be a useful concept in identifying genomic regions that contain silencers which can repress target genes either in proximity or via long-range chromatin interactions. The target genes may be tumor suppressors in cancer cells, and also cell fate-associated genes that need to be turned off for differentiation to occur.

We found several hundred MRRs in the K562 chronic myelogenous leukemia cell line, which showed dense chromatin interactions to target genes and to other MRRs. Next, we experimentally validated two looping silencers through CRISPR mediated removal, and both showed upregulation of target genes indicating that they are indeed *bona fide* silencers. Through CRISPR excision of one of the *IGF2* looping silencer components and one of the *FGF18* looping silencer components, we found that silencers control cell identity as their removal caused cell identity changes. Using the silencer at *IGF2* as an example, we dissected the consequences of silencer removal through 4C and ChIP-Seq with H3K27me3 and H3K27ac. We found that removal of a component of a silencer by CRISPR excision leads to changes in chromatin loops. Remarkably, regions that originally presented with very high H3K27me3 levels were stable in terms of chromatin loops while chromatin interactions to regions with low H3K27me3 and high H3K27ac levels changed. Moreover, genes in close proximity to, and genes that loop to MRRs by long-range chromatin interactions, were more susceptible to EZH2 inhibition. These genes showed higher levels of upregulation upon EZH2 inhibition, as compared with genes in close proximity or which loop to typical H3K27me3 peaks. EZH2 inhibition led to changes in long-range chromatin interactions at MRRs.

Taken together, our results indicated that clustering of H3K27me3 peaks in a manner similar to the super-enhancer analyses can identify MRRs that contain silencers that can loop over to target gene promoters. Silencer perturbation by H3K27me3 depletion and CRISPR excision led to epigenomic, transcriptomic and phenotypic consequences.

## Results

### Identification and characterization of H3K27me3-rich regions (MRRs) in the human genome

We identified highly H3K27me3-rich regions (MRRs) from cell lines using H3K27me3 ChIP-seq data^35^ in the following manner: we first identified H3K27me3 peaks, then clustered nearby peaks, and ranked the clustered peaks by average H3K27me3 signals levels. The top clusters with the highest H3K27me3 signal were called as “H3K27me3-rich regions” (MRRs) and the rest were called as “typical H3K27me3” regions (Figure 1A, 1B). The peaks that were merged together during this process were called constituent peaks. This method is similar to how super-enhancers were defined^33,36^. Recently, Pang and Snyder identified a list of silencer elements in K562 cells using a lentiviral screening system called ReSE^29^. We overlapped our list of MRRs in K562 with the list of silencers that identified by ReSE and found that 10.66% of ReSE silencer elements overlap with our MRRs (Figure 1C). This overlap percentage of 10.66% between our MRR and the ReSE silencer elements is significantly higher when compared to random expectation (Figure 1C). Although typical H3K27me3 peaks also have more overlap when compared with expectation, the differences in the percentage between actual and expected overlap percentage are larger for MRR (Figure 1C). This indicated that MRRs can be used to identify silencers in the genome. While the overlap percentage between our MRR and ReSE silencer elements is higher than random expectation, it is still relatively low compared with the total number of ReSE elements, which could be because ReSE elements contain other types of silencers such as DNA hypomethylated regions.

**Figure 1.**
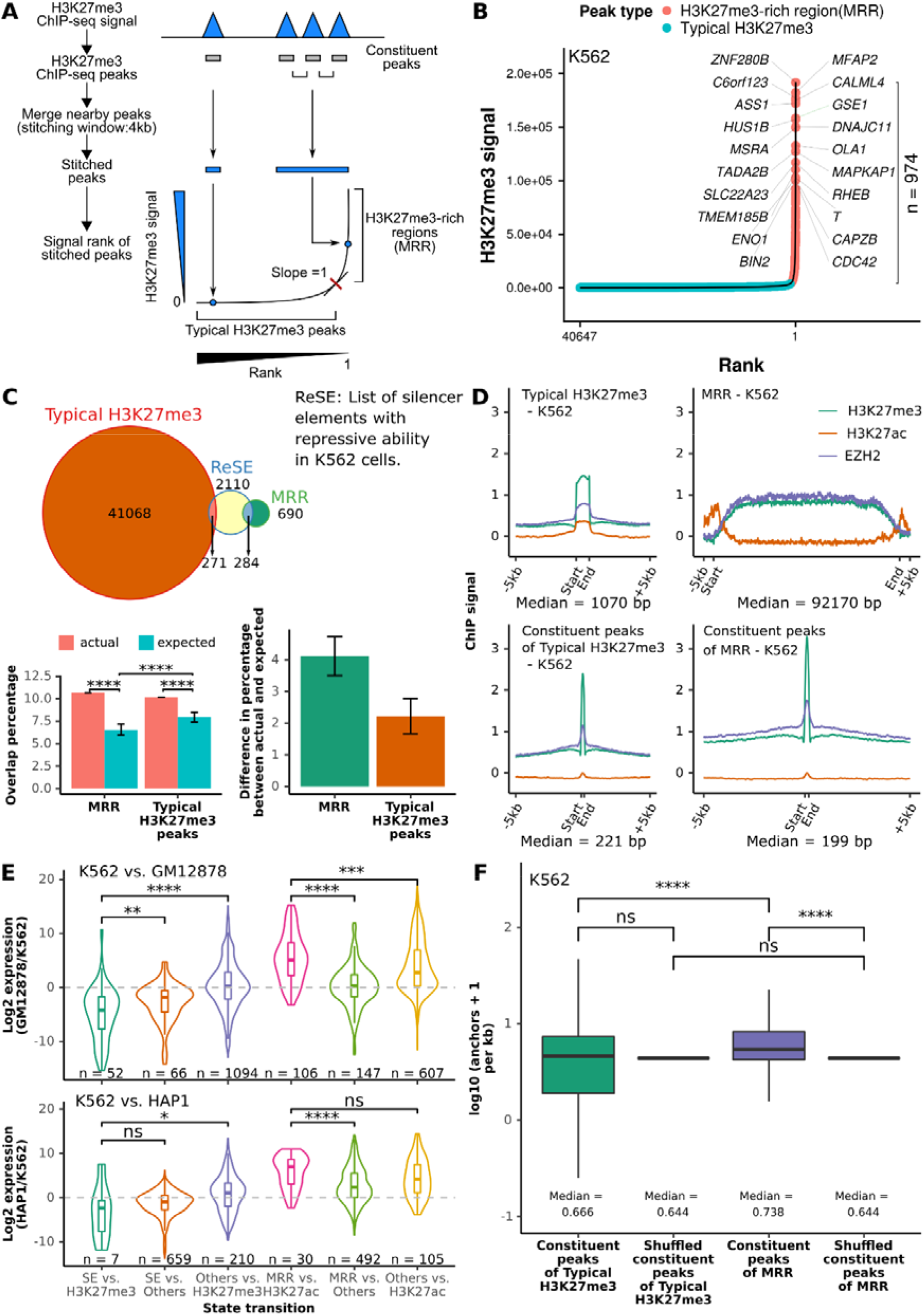
Definition of H3K27me3-rich regions (MRRs) and their characterization. **A.** H3K27me3 ChIP-seq peaks within 4kb are stitched together and the stitched peaks ranked according to their H3K27me3 signal. The rank-ordered signal with a slope of 1 is used as cut-off for defining H3K27me3-rich MRRs. Constituent peaks, peaks that are stitched during the process of merging peak. **B.** H3K27me3-rich regions (MRRs) and typical H3K27me3 peaks in K562 and their associated genes. A representative overlapping gene from each of the top 10 MRRs was shown. **C.** Overlap of MRR and typical H3K27me3 with ReSE list^29^. The Venn diagram shows the observed overlap between our MRR (H3K27me3-rich region)/typical H3K27me3 peaks and the ReSE list. Left barplot: The barplots show the percentage of the overlap number relative to total number in ReSE list. Actual, observed overlap percentage; expected, expected overlap percentage generated by: 1) first randomly shuffled 1000 times on MRR/typical H3K27me3 peaks on the same chromosome; 2) calculated overlap percentage using the 1000 time randomly shuffled regions. Wilcoxon test p values are indicated, ns: p > 0.05, *: p <= 0.05, **: p <= 0.01, ***: p <= 0.001, ****: p <= 0.0001. Right barplot: The difference between actual and expected percentage. Difference in percentage is calculated by actual percentage minus expected percentage. **D.** ChIP-seq signal on typical H3K27me3, MRR, constituent peaks of typical H3K27me3 peaks, and constituent peaks of MRR regions in K562. Peaks are scaled to the same median length of peaks in typical H3K27me3 (1070 bp), MRR (92170 bp), constituent peaks of typical H3K27me3 (221 bp), or constituent peaks of MRRs (199 bp), and the plot expanded by 5kb on both sides of the peak. **E.** Expression changes associated with different peaks between different cells. K562 vs. GM12878/K562 vs. HAP1 cell lines used in the comparison. Genes are classified based on the states of their overlapping peaks in different cell lines: [state in the first cell line] vs. [state in the second cell line], where the state can be super-enhancer (SE), H3K27me3-rich region (MRR), typical enhancer (H3K27ac), typical H3K27me3 peak (H3K27me3), or no overlapping peaks (Others). The expression data is from Epigenetic RoadMap^68^ and in-house HAP1 RNA-seq. Wilcoxon test p values are as indicated. **F.** Constituent peaks of MRRs have more Hi-C interactions compared to the constituent peaks of typical H3K27me3. Constituent peaks are peaks that form MRRs as described in A. The shuffled peaks are generated by expanding the midpoint of each constituent peaks to the median length of all the constituent peaks, and then followed by random genomic region shuffling. Wilcoxon test p values are indicated, ns: p > 0.05, *: p <= 0.05, **: p <= 0.01, ***: p <= 0.001, ****: p <= 0.0001.

The number of constituent peaks and overlapping genes at MRRs is larger than typical H3K27me3 peaks (Figure S1A, S1B). Considering the differences in the lengths of MRRs and typical H3K27me3 peaks, we used constituent peaks of MRRs and typical H3K27me3 peaks to study CpG methylation and gene features. The results showed that the constituent peaks of MRRs and typical H3K27me3 peaks mostly overlap with inter CpG island methylation (Figure S1C) and the intronic regions of genes (Figure S1D).

Many MRR-overlapping genes in different cell lines are known or predicted tumor suppressor genes^37^ (Figure S1E). For example, *NPM1*, the most commonly mutated gene in leukemia^38–41^, overlaps with an MRR in the leukemic cell line K562. *FAT1*, which is frequently mutated in chronic lymphocytic leukemia (CLL) and can act as a tumor suppressor through inhibiting Wnt signaling^42,43^, also overlaps with an MRR in K562. Gene ontology analysis showed that MRR-related genes are enriched in developmental and differentiation processes, while genes associated with typical H3K27me3 peaks are enriched in cell metabolism and transportation processes (Figure S1F, S1G). These results suggested that MRR may regulate important genes related to development and tumorigenesis.

ChIP-seq signals of EZH2 showed high correlation with H3K27me3 signal at typical H3K27me3, MRRs, constituent peaks of typical H3K27me3 and constituent peaks of MRRs, which is consistent with EZH2’s role in H3K27me3 mark deposition (Figure 1C; Figure S1H, S1I). Notably, the constituent peaks of MRRs had higher H3K27me3 and EZH2 signals than the constituent peaks of typical H3K27me3 peaks. This suggests that there are genomic regions with higher level of H3K27me3 and EZH2 compared with others, and they can be found in MRRs. In addition, the ChIP-seq profiles of SUZ12 and BMI1 are also higher in the constituent peaks of MRRs, suggesting that these regions may be targeted by PRC1 and PRC2 complex (Figure S1J, S1K).

MRRs were different in different cell lines, where a same gene can overlap with different types of peaks (Figure S1L – S1N). For example, the cadherin-like coding gene *CPED1* is covered by a broad MRR in GM12878, but overlaps with a super-enhancer in K562 (Figure S1L). Conversely, the gene for *DENND2D* is associated with an MRR but overlaps with a super-enhancer in GM12878 (Figure S1L). In addition, most MRRs were unique to individual cell lines (Figure S1O).

Analysis of cell line expression data showed that genes which are MRR-associated in one cell line, but H3K27ac peak-associated in a second cell line were upregulated in the second cell line, while genes that are super enhancer-associated in one cell line but are H3K27me3 peak-associated in a second cell line were down-regulated in the second cell line (Figure 1E). This observation is consistent with previously identified elements with dual function in both enhancing and silencing in mouse, human^43^, and *Drosophila*^44^. The expression fold changes between repressive and active state are higher than those genes that merely lost MRR or SE (Figure 1E; MRR vs. others and SE vs. others) or gained H3K27ac or H3K27me3 (Figure 1E; others vs. H3K27ac and others vs. H3K27me3), respectively. Further, genes whose expression were more cell line-specific were associated with more MRRs than those genes with lower expression specificity (Figure S1P). The uniqueness and specificity of MRRs suggested they might be primed for specific regulation in different contexts.

We overlapped MRRs with high-resolution *in situ* Hi-C data^45^, and found that constituent peaks of MRRs had a higher density of chromatin interactions than the constituent peaks of typical H3K27me3 peaks in both K562 and GM12878 (Figure 1F; Figure S1Q, S1R). The involvement of chromatin interactions in MRRs was similar to super-enhancers compared with typical enhancers^46^, which suggested that chromatin interactions might be important within regions rich in histone modification.

In summary, we defined MRRs using H3K27me3 ChIP-seq peaks, and showed that MRRs might be involved with specific gene repression related to development, differentiation and cancer via chromatin interactions.

### H3K27me3-rich regions (MRRs) preferentially associate with MRRs in the human genome via chromatin interactions

We assigned chromatin states at Hi-C interaction anchors using H3K27me3 and H3K27ac peaks: active (A) anchors overlap with H2K27ac peaks, repressive (R) anchors overlap with H3K27me3 peaks, bivalent (B) anchors overlap with both H3K27me3 and H3K27ac peaks, and quiescent (Q) anchors overlap with neither peak (Figure 2A). We further defined the chromatin state pair of an interaction as the chromatin states of its anchors and calculated the proportion of different chromatin interaction in the Hi-C data (Figure 2B, “Obs”). Next, we calculated the expected proportion of interactions for each state pair under a homogeneous model (Figure 2B, Exp), and compared those expectations to the actual number of observations (Figure 2B, log_2_(Obs/Exp) on the x-axis). If the observed proportion of a certain category of interactions were more frequently seen, the log_2_(Obs/Exp) value would be positive; conversely, if a certain category was depleted, the log_2_(Obs/Exp) value would be negative.

**Figure 2.**
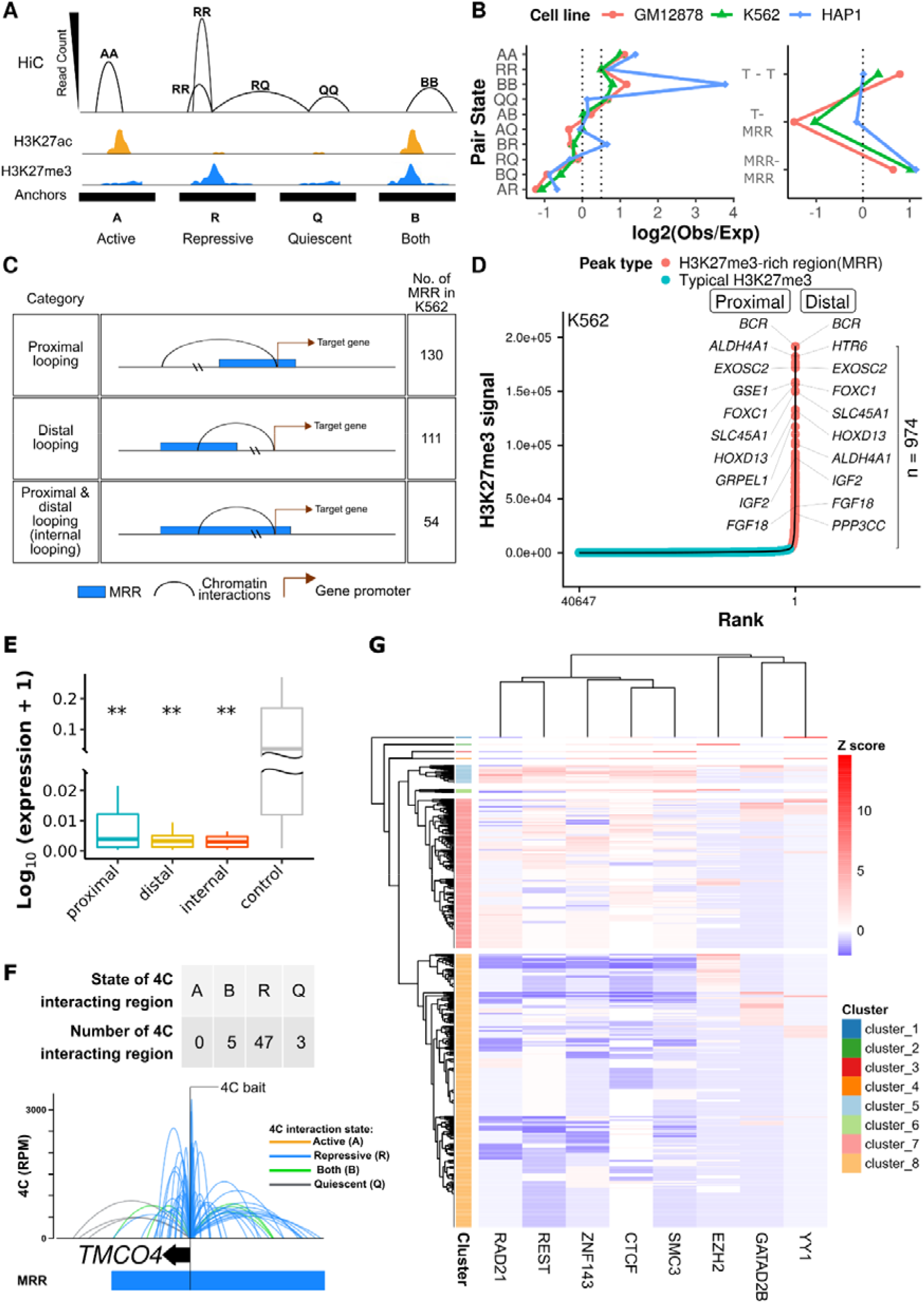
H3K27me3-rich regions (MRRs) preferentially associate with MRRs in the human genome via chromatin interactions. **A.** Schematic plot of how different categories of Hi-C interactions are defined. Hi-C anchors are classified by whether they overlap with H3K27me3 or H3K27ac peaks. A (active), overlap with only H3K27ac peaks; R (repressive), overlap with only H3K27me3 peaks; Q (quiescent), overlap with neither H3K27ac nor H3K27me3 peaks; B (both), overlap with both H3K27ac and H3K27me3 peaks. The height of Hi-C interactions (arcs) represents the highest read counts in the interacting regions. **B.** Observed/expected ratio of Hi-C interactions in different categories. Left: categories of chromatin pair states. Right: T (typical H3K27me3) or H (MRR) peaks. The expected interactions are calculated from the marginal distributions of different anchors. **C.** Different categories of MRR associated with genes. **D.** H3K27me3-rich regions (MRRs) and typical H3K27me3 peaks in K562 and their associated genes through chromatin interactions. Peaks overlapping with Hi-C interactions are labeled with associated genes: for peaks labeled “proximal”, the gene TSS and peak occupy the same Hi-C anchor; “distal” peaks are connected to the gene via Hi-C interactions. **E.** Expression of genes that are associated with MRR in proximal, distal, and internal category in K562 cells. The three categories are described in **C**. The control category is generated by: 1) first filtering out genes that are overlapped with ENCODE blacklist regions and also H3K9me3 peaks as H3K9me3 is associated with constitutive heterochromatin and such regions are likely to be highly silenced; 2) only retaining genes that overlapped with Hi-C interactions; 3) randomly sampling the same amount of genes as the average gene number in proximal/distal/internal category. Wilcoxon test p values are indicated, ns: p > 0.05, *: p <= 0.05, **: p <= 0.01, ***: p <= 0.001, ****: p <= 0.0001. **F.** Example of 4C at the *TMCO4* gene promoter bait showing extensive internal looping within an MRR in K562. The colors of 4C interactions are based on the distal interacting regions to the 4C bait. Blue: repressive; orange: active; green: both; grey: quiescent. The state of the 4C bait is labeled by text. Each ChIP-seq tracks contains ChIP signal and peaks. TE, typical enhancer; SE, super-enhancer; T, typical H3K27me3; MRR, H3K27me3-rich region. **G.** Heatmap of transcription factors binding enrichment at interacting regions of MRRs. Each row representing an interacting region of MRRs. The number overlapping transcription factor peaks at interacting regions are normalized to Z score per transcription factor. Red colors indicate more binding events.

Interactions between anchors of the same state (AA, RR, and BB) were more likely to interact with each other, while interactions with vastly different chromatin state pairs (e.g., AR, BQ) less likely (Figure 2B, left), regardless of cell line. When grouped into typical H3K27me3 peaks (T) versus high H3K27me3 regions or MRRs (MRR), the high H3K27me3 regions showed a preference for interactions with other MRRs (Figure 2B, right). In keeping with A/B chromatin compartments of the nucleus, this ‘like-like’ preference indicated that loci of similar chromatin state were more prone to interact with each other.

To further explore the potential regulatory role of MRRs in chromatin interactions, we identified the subset of MRR-anchored interactions where at least one anchor peak overlapped a gene transcription start site, and grouped them according to whether the MRR anchor was proximal or distal to the TSS anchor (Figure 2C, 2D; Figure S2A-S2F, S2G; examples of genes can be found in Figure S2I-S2L). Both proximal and distal gene looping occur for MRR-anchored interactions, but some MRRs are large enough that both anchors occur in the same MRR. While proximal looping genes are a subset of the genes within MRRs, distal looping genes are only identified by chromatin interactions (Figure 2D, right panel). The expression of genes that are proximally, distally, or internally associated with MRR are lower than randomly sampled genes that are involved in chromatin interactions (Figure 2E; Figure S2H). The difference in gene expression levels between proximal, distal, and internal categories is not significant, suggesting that distal looping by MRRs is associated with reduced gene expression to a similar extent as proximal regulation by MRRs (Figure 2E). There is no significant difference between proximal, distal and internal categories, thus showing that genes regulated by distal looping may be silenced to the same extent as genes proximal to MRRs. This indicated the importance of long-range looping in mediating silencing between distal regulatory elements and gene promoters. The top-ranking MRRs are often involved in extensive internal looping (Figure S2K-S2L). Gene ontology analysis showed that MRR-associated genes in the context of chromatin interactions are involved in developmental and differentiation processes (Figure S2M).

In order to validate the ‘like-like’ preference of chromatin interactions, we performed Circular Chromosome Conformation Capture (4C) experiments on selected loci at MRR to investigate the associated chromatin interactions in a comprehensive and high-resolution manner. We annotated the interactions based on the chromatin state of the anchor distal from the bait in K562 (Figure 2F and Figure S2N-P), and across multiple cell lines (Figure S2Q-R). The interaction profiles of 4C baits of different states were largely dominated by interacting regions of the same state as the baits. In addition, the *TMCO4* 4C data showed that most 4C interactions fell within the same MRR as the bait and only a handful of them were outside of the MRR. This suggested that MRR can have extensive internal looping.

We also carried out 4C experiments on the same bait across different cell lines. The interactions and the chromatin state at the bait locus varied in different cell lines, but the interaction profile maintained a preference for the same chromatin state as the bait (Figure S2Q, S2R). As a further test of this concept, the extensive BB long-range interactions (green arcs) connecting *PSMD5* and *TOR1A* in K562 were validated using reciprocal 4C bait design. When the *PSMD5* bait region was A (active) in either GM12878 or HAP1 cells, the BB interactions were largely reduced and other types of interactions started to appear (Figure S2Q).

Next, we analyzed the transcription factors binding to the regions of MRRs that are connected by chromatin interactions. ChIP-seq peaks of chromatin architectural proteins (CTCF, YY1, ZNF143), cohesin subunits (RAD21, SMC3), and transcription repression-associated proteins (EZH2, REST, GATAD2B) were downloaded from ENCODE and overlapped with the interacting regions of MRRs, which were then normalized to Z-score and clustered by hierarchical clustering. Specific enrichments of one specific transcription factor can be found in several small clusters (Figure 2G; YY1 in cluster_1, EZH2 in cluster_2, and SMC3 in cluster_3). Another cluster was identified with very high binding affinity of RAD21, REST, ZNF143, CTCF, and SMC3 (Figure 2G cluster_5). Our results demonstrated that different chromatin architectural proteins are involved in the regulation of different silencer-associated chromatin interactions.

### CRISPR excision of a looping anchor within an MRR (MRR1-A1) leads to upregulation of multiple genes like *FGF18*, cell differentiation and tumor growth inhibition

Next, we asked if MRRs function as silencers to regulate gene expression. We selected 2 MRRs for functional testing based on the H3K27me3 signal, the presence of Hi-C anchors and the number of Hi-C anchors they associated with whether the genes were involved in cell identity (Supplementary Text). Briefly, there are 974 MRRs in K562 (Figure S3A) and of those MRRs, 237 MRRs are associated with genes. Among these, 130 MRRs show proximal looping to genes (MRRs overlap with target gene promoters), 111 MRRs show distal looping to genes (MRR loops over to the promoter of target gene by long-range chromatin interactions) and 51 MRRs show internal looping to genes (part of the MRR overlaps with the target gene promoter and the other part of the MRR loops over to the promoter of the target gene by long-range chromatin interactions). From this list, we selected MRR1, an internal looping example which showed 2 Hi-C loops to *FGF18*, a fibroblast growth factor involved in cell differentiation and cell-to-cell adhesion^47,48^ (Figure 3A) and MRR2, an internal looping example which showed 3 Hi-C loops to *IGF2*, an imprinted gene known to be associated with genomic silencers^50^ and involved in growth, development and cancer^49^ (Figure 5A).

**Figure 3.**
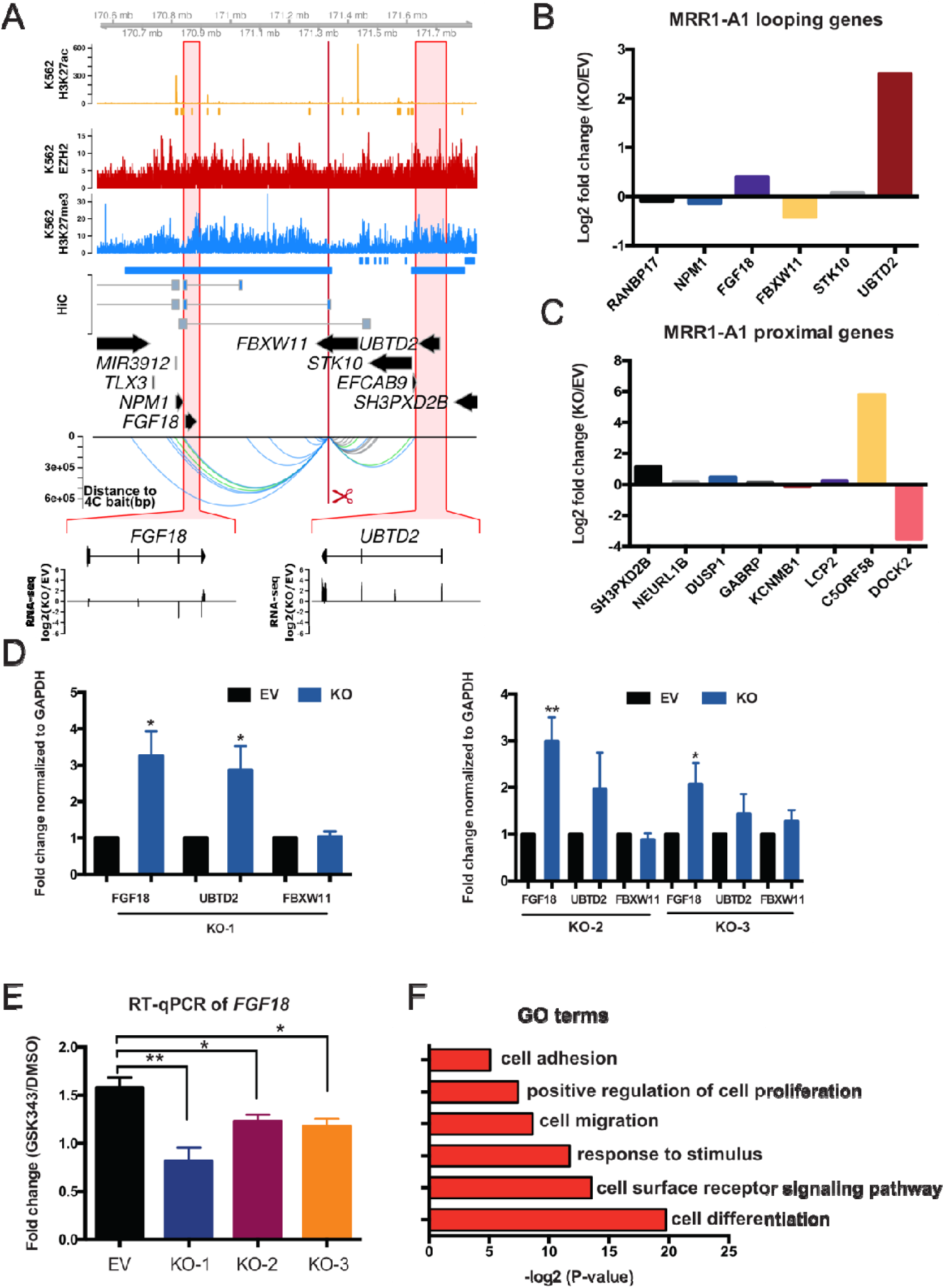
CRISPR excision of MRR1-A1 leads to gene upregulation of multiple proximal and looping genes including *FGF18*. **A.** Screenshot showing EZH2 ChIP-seq, H3K27me3 ChIP-seq, H3K27ac ChIP-seq and chromatin interactions as identified from previously published Hi-C data^22^, gene information, and 4C performed on the CRISPR-excised region in wild-type cells confirming chromatin interactions to *FGF18*, as well as showing chromatin interactions to *UBTD2* and other genes. The regions highlighted in the red boxes are shown in more detail, with RNA-seq was shown as one CRISPR knockout clone over wild-type at *FGF18* and *UBTD2*. The blue bar shows the predicted whole MRR. The red box with the red scissors indicates the region which was excised. **B.** RNA-seq fold changes calculated from two replicates of RNA-Seq data of MRR1-A1 looping genes in one MRR1-A1 knockout clone (KO) as compared with one vector control clone (“Empty Vector”; “EV”)**. C.** RNA-seq fold changes of MRR1-A1 proximal genes in KO as compared with EV. **D.** RT-qPCR of *FGF18*, *UBTD2* and *FBXW11* in three different CRISPR-excised clones (“KO-1”, “KO-2”, “KO-3”) as compared with EV. **E.** RT-qPCR of *FGF18* expression upon GSK343 treatment in EV and three KO clones. Fold change was plotted compared to *GAPDH* for EV and KO cells in DMSO and GSK343 condition. **F.** Gene Ontology (GO) was performed using significant differentially expressed (DE) genes in the RNA-seq data which was shown as – log_2_(p value). All data shown here indicates average + standard error. P value less than 0.05 is shown as *. P value less than 0.01 is shown as **.

We designed the CRISPR deletion site at a 1 kb region in MRR1 (termed “MRR1-A1”) located in the *FBXW11* intronic region that was associated with one of two Hi-C anchors that loop over to *FGF18* (Figure 3A). This region has high H3K27me3 as validated by ChIP-qPCR (Figure S3B). MRR1-A1 is part of cluster_8 (associated with low levels of cohesin proteins, high binding to GATAD2B; Table S8) from Figure 2G. We performed 4C using MRR1-A1 as the bait to detect all the genomic locations that have chromatin interactions with this region in wild-type K562. The 4C-seq results showed that this region indeed had chromatin interactions with *FGF18* and several other genes such as *NPM1* and *UBTD2* (Figure 3A).

Next, we performed CRISPR deletion and generated three knock out (KO) clones (Figure S3C). To scan for target genes, we prepared RNA-seq from one KO clone and aligned this data with the 4C-seq data using MRR1-A1 as the bait (Figure 3A). From RNA-seq fold changes of MRR1-A1 looping genes, we found upregulation of *FGF18* and *UBTD2* (Figure 3B, Figure SD). For proximal genes, we found upregulation of genes including *SH3PXD2B* and *C5ORF58* (Figure 3C, Figure S3E). Among those genes, upregulation of the *FGF18* was further confirmed by RT-qPCR consistently in three different KO clones (Figure 3D) while *UBTD2* was upregulated significantly in KO1 but not in other clones. Therefore, we focused on *FGF18* gene for further analysis.

Next, we treated the K562 cells with GSK343 (EZH2 methyltransferase inhibitor). Upon GSK343 treatment, *FGF18* gene was upregulated compared with DMSO control. This indicates that *FGF18* gene was upregulated upon H3K27me3 depletion. By contrast, in MRR1-A1 KO clones treated with GSK343, *FGF18* was upregulated to a much smaller extent as compared with wild-type cells (Figure 3E). This indicates that *FGF18* gene upregulation upon H3K27me3 depletion is partially dependent on intact MRR1-A1, which further suggested that MRR1-A1 is a silencer.

To explore if the MRR1 is cell type specific, we identified MRRs in seven cell lines and found that MRR1 is specific to two of the seven cell lines, K562 and GM12878 (Figure S4B) which suggested that silencers are specific to different cell types and might control the cell identity related genes.

Since *FGF18* has been reported to be involved in cell-to-cell adhesion^47,48^, next we asked if KO clones showed changes in adhesion. To address this, we performed gene ontology (GO) analysis which showed that KO clones may undergo cell adhesion and cell differentiation (Figure 3F). First, we observed that the KO clones show increased adhesion to the cell culture plate surface and formed aggregates while wild type cells remained as suspension cells (Figure 4A). The adhesion ability was further quantified by cell adhesion assays (Figure 4B).

**Figure 4.**
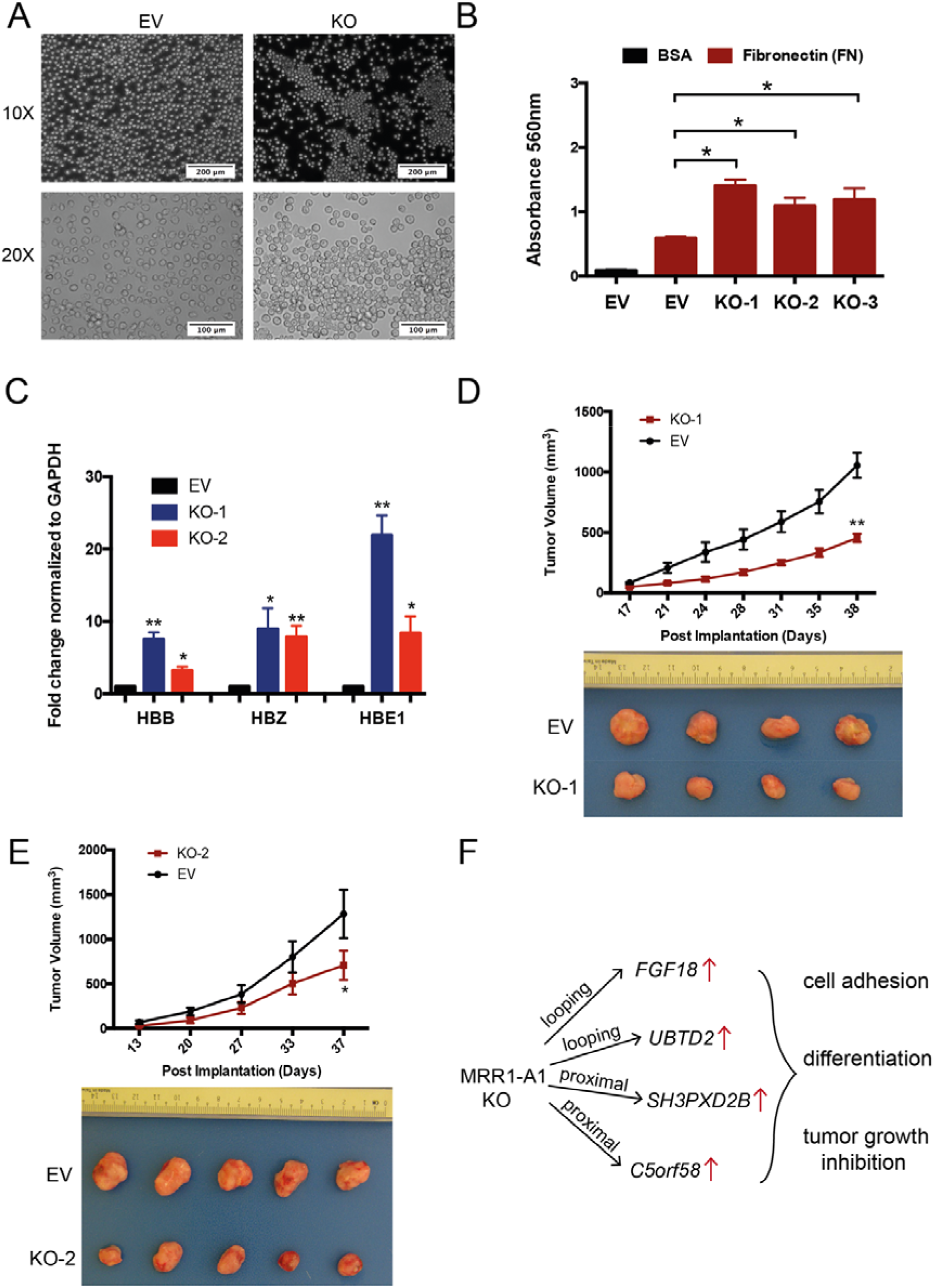
CRISPR excision of MRR1-A1 leads to altered adhesion, erythroid differentiation and tumor growth inhibition. **A.** Light microscopy photos of empty vector (EV) and CRISPR knockout clones (KO) showing increased cell adhesion and aggregates in the KO clones. 10X and 20X magnification were shown. **B.** A fibronectin adhesion assay showed increased adhesion of the three CRISPR knockout clones (KO) as compared with empty vector (EV). Bovine Serum Albumin (BSA) was used as a negative control. **C.** RT-qPCR of haemoglobin genes (*HBB*, *HBZ* and *HBE1*) in EV and two KO clones. **D&E**. Tumor growth in SCID (Severe Combined Immunodeficiency) mice injected with MRR1-A1 knock out clones and empty vector cells (EV). The upper panel shows the tumor growth curve, and data shown as tumor volume with different post implantation days. The panel below was the representative tumor picture at the final day. **F.** Model of MRR1-A1 excision leads to multiple genes change which further leads to cell adhesion, differentiation and tumor growth inhibition. All data shown here indicates average + standard error. P value less than 0.05 is shown as *. P value less than 0.01 is shown as **.

**Figure 5.**
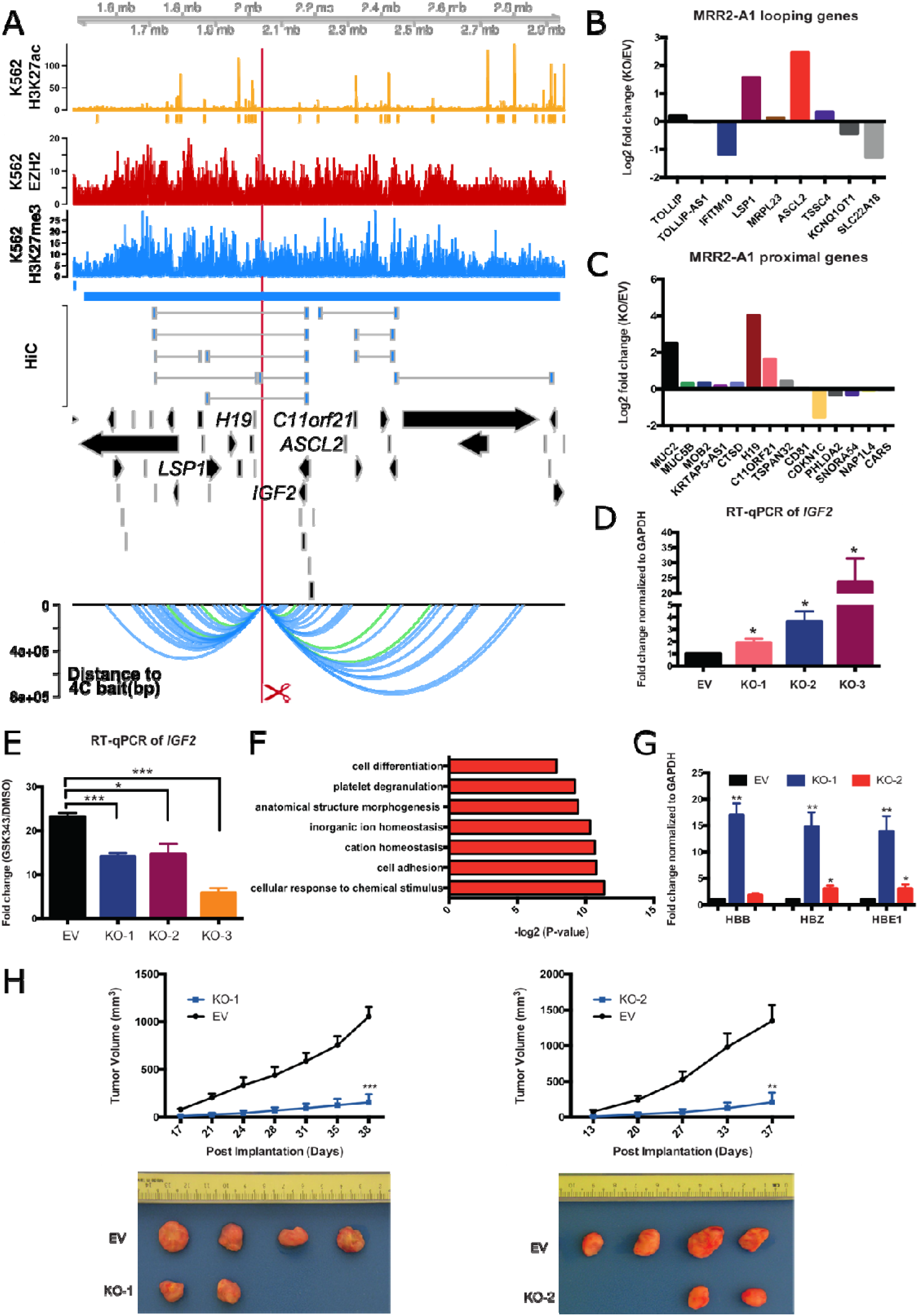
CRISPR excision of MRR2-A1 leads to multiple gene upregulation including *IGF2* gene, erythroid differentiation and tumor growth inhibition. **A.** Screenshot showing EZH2 ChIP-seq, H3K27me3 ChIP-seq, H3K27ac ChIP-seq and chromatin interactions as identified from previously published Hi-C data^22^, gene information, and 4C performed on the CRISPR-excised region in wild-type cells confirming chromatin interactions to *IGF2* as well as other genes. The blue bar shows the predicted MRR. The red box with the red scissors indicates the region which was excised. **B.** RNA-seq fold changes of MRR2-A1 looping genes in KO as compared with EV. **C.** RNA-seq fold changes of MRR2-A1 proximal genes in KO as compared with EV. **D.** RT-qPCR of *IGF2* in three different CRISPR-excised clones (KO-1, KO-2, KO-3) as compared with vector control cells (“EV”). **E.** RT-qPCR of *IGF2* expression upon GSK343 treatment in EV and three KO clones. Fold change was plotted compared to *GAPDH* for EV and KO cells in DMSO and GSK343 condition. **F.** Gene Ontology (GO) was performed using significant DE genes in the RNA-seq data shown as −log2(p value). **G.** RT-qPCR of haemoglobin genes (*HBB*, *HBZ* and *HBE1*) in EV and two KO clones. **H**. Tumor growth in SCID (Severe Combined Immunodeficiency) mice injected with MRR2-A1 knock out cells and empty vector cells (EV). The upper panel shows the tumor growth curve, and data shown as tumor volume with different post implantation days. The panel below was representative tumor picture at the final day. All data shown here indicates average + standard error. P value less than 0.05 is shown as *. P value less than 0.01 is shown as **. P value less than 0.001 is shown as ***.

Next, because *FGF18* is associated with differentiation^47,48^, we investigated whether KO clones showed an erythroid differentiation phenotype. Cellular aggregates were reported by several publications^51,52^ to be associated with cell differentiation such as erythroid and megakaryocyte lineage of K562 cells. Therefore, we checked the expression of haemoglobin genes which can be the indicator of erythroid lineage differentiation^53^ in the RNA-seq data and further confirmed some of their upregulation (*HBB*, *HBZ* and *HBE1*) by RT-qPCR (Figure 4C).

To investigate whether the differentiation phenotype might be partially caused by upregulation of *FGF18*, we performed siRNA knock down targeting *FGF18* gene in the KO clones, which led to 60%-80% reduction in *FGF18* gene expression levels. Haemoglobin genes can be partially rescued by *FGF18* knocking down (Figure S4A) which suggested that erythroid differentiation partially caused by *FGF18* upregulation (Figure 4F). We speculate that *FGF18* siRNA knockdown did not lead to a complete rescue because MRR1-A1 knockout also upregulates other genes in addition to *FGF18*. For example, *SH3PXD2B* may also play roles in controlling erythroid differentiation^54^.

Leukemic cell differentiation induction is associated with cell growth inhibition and small molecule inhibitors such as All-*trans* Retinoic Acid (ATRA) that can induce differentiation have been useful in treatment of Acute Promyelocytic Leukemia, suggesting that methods to induce differentiation could lead to potential leukemia treatments^53,55^. Therefore, we asked if silencer removal is associated with growth inhibition *in vivo*, given that silencer removal leads to cell differentiation. To test this, we performed xenograft experiments for two different KO clones and both two KO clones showed inhibition of tumor growth in the mice (Figure 4D and 4E). This tumor growth inhibition suggested that MRR1-A1 might play tumor suppressor roles in leukemia and suggests the possibility that silencers can control cell identity through repression of tumor suppressor gene expression. In summary, our analyses suggested that MRR1-A1 can function as a looping silencer of *FGF18* as well as other genes and MRR1-A1 removal leads to cell identity changes such as cell adhesion, cell differentiation and tumor growth inhibition (Figure 4F).

### CRISPR excision of a looping anchor within an MRR (MRR2-A1) leads to multiple gene upregulation including *IGF2*, cell differentiation and tumor growth inhibition

MRR2 was characterized in the same manner as MRR1. Specifically, we designed another 1 kb deletion in MRR2 (termed “MRR2-A1”) located in an intergenic region 10 kb away from the long non-coding RNA *H19* that was associated with one of three Hi-C anchors looping over to *IGF2* (Figure 5A). High H3K27me3 signal of MRR2-A1 was confirmed by ChIP-qPCR (Figure S5A) and chromatin interactions to *IGF2* and other genes were confirmed by 4C-seq (Figure 5A). The MRR2-A1 anchor was in cluster_5 in Figure 2G, and it has high binding affinity of CTCF, RAD21, SMC3 and REST (Table S8).

RNA-seq of one MRR2-A1 KO clone (Figure S5B) showed upregulation of multiple genes which loop to MRR2-A1 (looping genes) including *LSP1*, *ASCL2* and *TSSC4* (Figure 5B, Figure S5C). For proximal genes, *MUC2*, *H19* and *C11ORF21* were upregulated in KO (Figure 5C, Figure S5D). H3K27me3 and H3K27ac ChIP-seq of this KO also showed changes in H3K27me3 and H3K27ac levels around MRR2 (Figure S5F). *IGF2* is expressed at a very low level in differentiated cells of the haematopoietic lineage^56^ and detected at very low levels by RNA-seq and therefore not shown in the fold change calculation. As *IGF2* has been previously shown to be regulated by silencers via chromatin interactions in mice^13^, we asked whether RT-qPCR could detect *IGF2* in our clones. Using RT-qPCR, we could detect *IGF2* and we found that *IGF2* was upregulated in all three KO clones (Figure 5D). By contrast, *H19* was upregulated in one of the three KO clones as measured by RT-qPCR (Figure S5E). This indicated MRR2-A1 can function as a looping silencer to repress *IGF2* in human K562 cells. Again, *IGF2* was upregulated upon GSK343 treatment and the level of upregulation was reduced by MRR2-A1 removal, which showed that MRR2-A1 is a silencer (Figure 5E). Similar to MRR1, MRR2 was also cell type specific (Figure S5I).

Through gene ontology (GO) analysis of the RNA-Seq on the MRR2-A1 KO clone, we found the term “cell differentiation” (Figure 5F). Thus, we asked if these KO clones also undergo erythroid differentiation. RT-qPCR showed the haemoglobin genes (*HBB*, *HBZ* and *HE1*) were upregulated in the KO clones (Figure 5G) and *IGF2* siRNA knock down can partially reduce this upregulation (Figure S5G) which suggests the differentiation was partially caused by *IGF2* upregulation in MRR2-A1 KO clones (Figure S5H). Similar to *FGF18* siRNA knockdown, we did not see a complete rescue of the differentiation phenotype by *IGF2* siRNA, which we also speculate might be because MRR2-A1 also upregulates other genes besides *IGF2*.

Finally, we tested to see whether the CRISPR KO clones showed tumor growth inhibition *in vivo*, similar to MRR1. Xenograft experiments showed severe tumor growth inhibition of two different clones (Figure 5H) which further suggests that silencers can control cancer growth. Therefore, this MRR2-A1 example together with MRR1-A1 example confirmed the existence of two looping silencers and showed that looping silencers are involved in the control of cell identity and tumor growth.

### *IGF2* looping silencer (MRR2-A1) removal caused changes of distant chromatin interactions

Through the previous two examples, we confirmed the existence of looping silencers and demonstrated they can control cell identity. Next, we investigated the epigenomic consequences of a looping silencer removal using the *IGF2* looping silencer (MRR2-A1) example. First, we asked whether chromatin interaction landscape will be changed upon looping silencer removal. We performed 4C-seq in the KO and control clones. Using *IGF2* as the bait, we detected there are 33 chromatin interactions lost and 12 chromatin interactions gained after MRR2-A1 knocking out while a control bait remains highly unchanged (Figure 6A, Figure S6A). Several lost loops were confirmed by 3C-PCR (Figure S6B) which indicates that looping silencer removal could lead to alterations in the chromatin interaction landscape.

**Figure 6.**
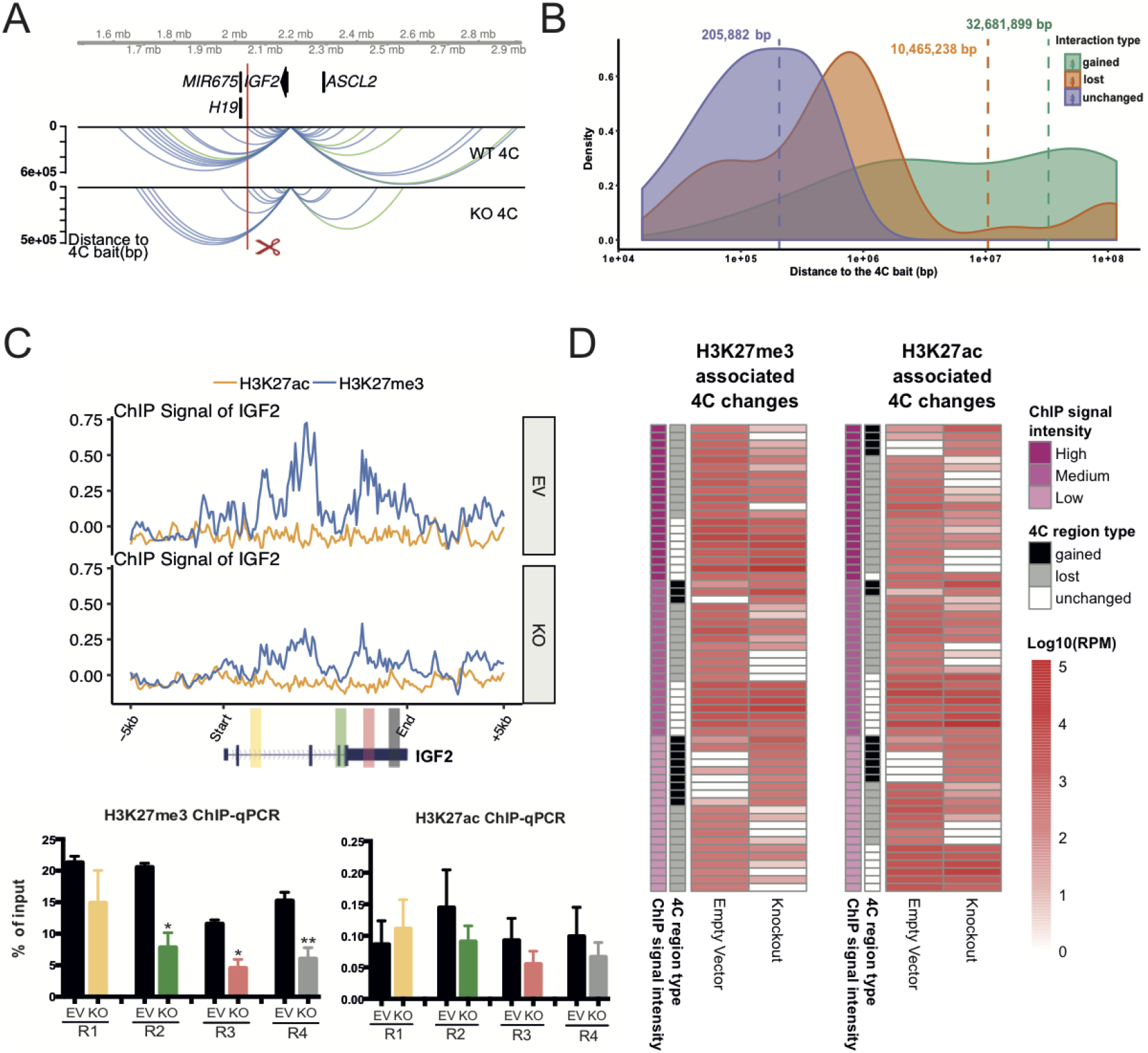
Initial histone states predict the changed loops upon MRR2-A1 removal. **A**. Representative chromatin interactions at *IGF2* bait in KO and control clones which shown as loops. **B**. The average distance of changed loops (gained loops and lost loops) is greater than unchanged loops upon MRR2-A1 KO when using *IGF2* promoter as the bait. **C**. ChIP-seq and ChIP-qPCR of H3K27me3 and H3K27ac for four regions (R1-R4) at *IGF2* gene in EV and KO clones. Data shown here are average + standard error. P value less than 0.05 is shown as *. P value less than 0.01 is shown as **. **D**. Heatmap about Integrative analysis of 4C, H3K27me3 and H3K27ac ChIP-seq in EV. Left panel: different 4C regions are classified according to their H3K27me3 signal intensity in EV. H3K27me3 signal level at these 4C regions are tertiled in three cohorts: high, medium, and low. 4C region type indicates different categories of 4C regions (Gained, lost and unchanged). The 4C interaction intensities are shown in log10 (RPM). Right panel: different 4C regions are classified according to their H3K27ac signal intensity in EV. Similar to the left panel, the H3K27ac signal level at these 4C regions are tertiled in three cohorts.

Next, we classified chromatin interactions into three types: unchanged loops, gained loops and lost loops to explore their features. Through mapping their distance and density, we found that the average distance of changed loops are greater than unchanged loops which indicates that the long-range chromatin interactions which are further away to the bait tend to change (Figure 6B). Moreover, the long-range chromatin interactions have a greater propensity to be lost than to be gained. Given that long-range chromatin interactions require more energy to be held together^57^, we speculated that when an anchor is lost, the amount of energy present in the system to hold together the long-range chromatin interactions may not be sufficient.

### Integrative analysis of histone modification states and chromatin interactions before and after *IGF2* looping silencer (MRR2-A1) removal

MRR2 has high H3K27me3 signals and histone modifications may play a key role in *IGF2* upregulation. Therefore, we performed H3K27me3 and H3K27ac ChIP-seq in the KO and control clones (Figure S6C). We found that H3K27me3 decreased along *IGF2* gene region upon knockout (Figure 6C) while a control region remained similar (Figure S6D). This suggested that silencer removal will cause H3K27me3 loss at the target gene region.

Next, we performed integrative analysis of 4C-seq and ChIP-seq. Surprisingly, we found that the initial histone states of the cells before knockout were associated with whether the chromatin interactions would be gained, lost or unchanged upon knockout of MRR2-A1 (Figure 6D). Specifically, very repressed loops with high H3K27me3 in control cells were unchanged or lost after KO. Loops with high H3K27ac and loops with low H3K27me3 in control cells tend to be easily changed either gained or lost after KO (Figure 6D).

Moreover, when we compared the integrative analysis in EV and KO, we observed significant decrease in H3K27me3 for unchanged loops while levels H3K27ac increased slightly (Figure 7A-B) which suggested that the repressive ability of the chromatin interaction became weaker and all the chromatin interactions looping to *IGF2* became more active in terms of histone state after MRR2-A1 KO. An example of the unchanged loops is shown in Figure 7C, which displays unchanged loops to *IGF2* promoter along with decreased H3K27me3 levels in KO. When examining the gained loops to *IGF2* gene, we observed increase in H3K27ac and no change in H3K27me3 (Figure 7A-B) indicating that *IGF2* promoter could also gain more active loops to activate gene expression. An example of the gained loops is shown in Figure 7D.

**Figure 7.**
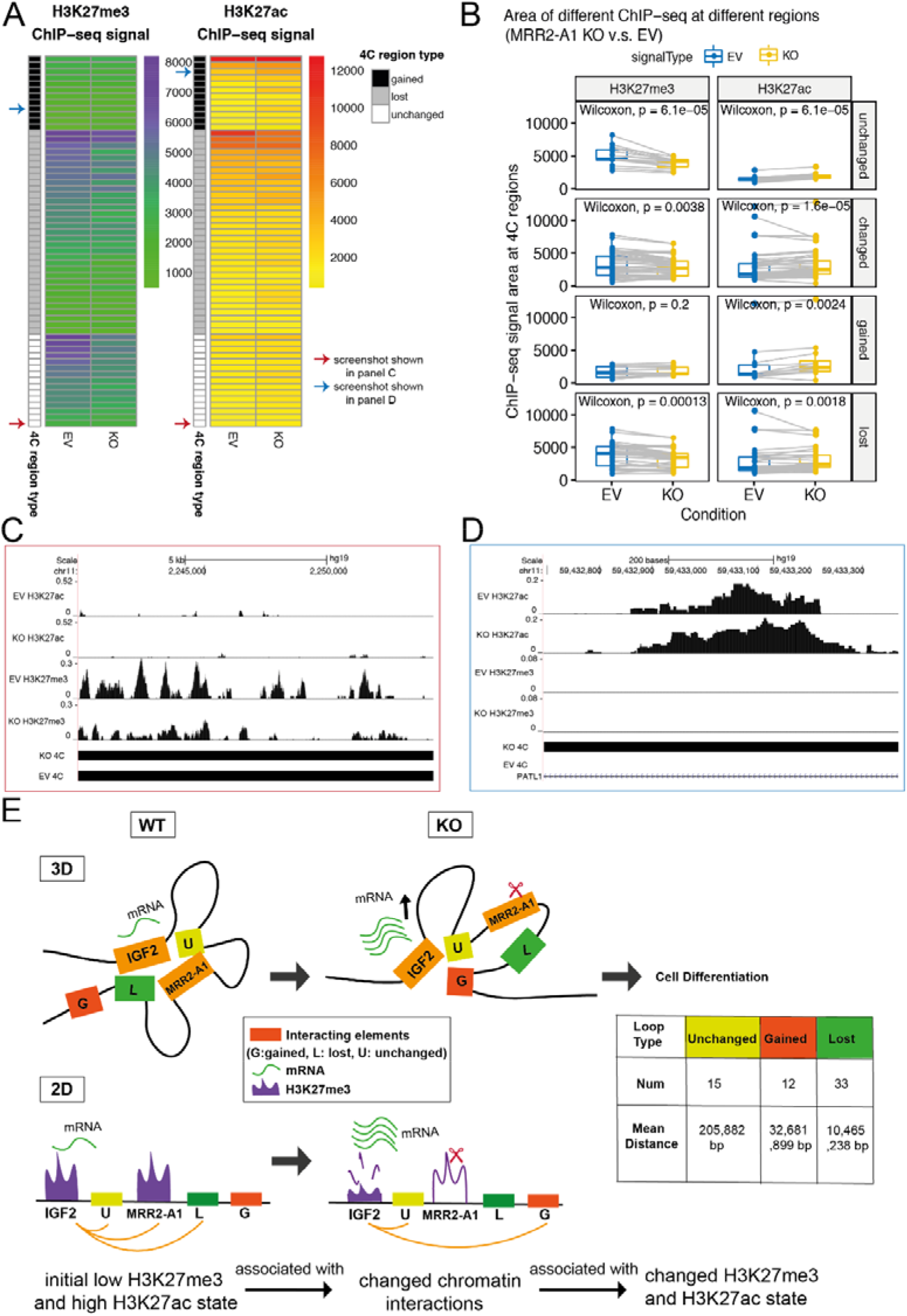
Unchanged loops and gained loops to *IGF2* become increased H3K27ac and decreased H3K27me3 levels upon MRR2-A1 removal. **A**. Heatmap of ChIP-seq signal changes of H3K27me3 and H3K27ac at different types of 4C regions (gained, lost and unchanged) in empty vector (EV) and MRR2-A1 KO clones. Blue arrow: this region is shown as a screenshot in panel C. Red arrow: this region is shown as a screenshot in panel D. **B**. Boxplots of ChIP-seq signal changes of H3K27me3 and H3K27ac at different types of 4C regions in EV and MRR2-A1 KO clones. The same 4C regions are connected by grey lines. Wilcoxon paired test p value are indicated. **C**. Zoomed screenshot about one of the unchanged 4C regions indicated in **A** which showed decrease of H3K27me3. **D**. Zoomed screenshot about one of the gained 4C regions in **A** which showed increase of H3K27ac. **E**. 3-dimensional and 2-dimensional cartoon schematics of our proposed model that initial histone states are associated with changed loops and MRR2-A1 removal leads to increase of H3K27ac levels on unchanged loops and gain of chromatin loops in regions with high H3K27ac levels.

Taken together, the regions that loop to *IGF2* in the KO clones are now more active with higher H3K27ac and lower H3K27me3 levels. These findings demonstrate two mechanisms by which I*GF2* might be upregulated in KO clones. First, *IGF2* showed gain of chromatin loops to more active anchors and losses of loops to several repressive anchors. Second, the retained loops which had strong H3K27me3 levels at the control cells became weaker after KO (Figure 7E). A combination of these mechanisms may operate in different cellular and physiological contexts.

### MRR-associated gene expression and long-range chromatin interactions are susceptible to EZH2 perturbation

In order to investigate the effects of H3K27me3 on MRR-associated chromatin interactions and associated gene expression, we eliminated or reduced H3K27me3 by EZH2 inhibitor treatment (GSK343) in K562 cells and CRISPR mediated knockout of EZH2 in HAP1 cells (a near haploid cell line derived from chronic myeloid leukemia).

After treatment with GSK343 in K562 cells, the levels of H3K27me3 decreased globally, leading to the loss of nearly half of the H3K27me3 ChIP-seq peaks (Figure 8A). However, there were still residual H3K27me3 peaks after GSK343 treatment, and these were the regions that had higher H3K27me3 signal before the treatment as compared with the susceptible peaks. Western blot confirmed that 1μM of GSK343 treatment in K562 cells and EZH2 knockout in HAP1 cells were sufficient to lead to global loss of H3K27me3 (Figure S7A-B).

**Figure 8.**
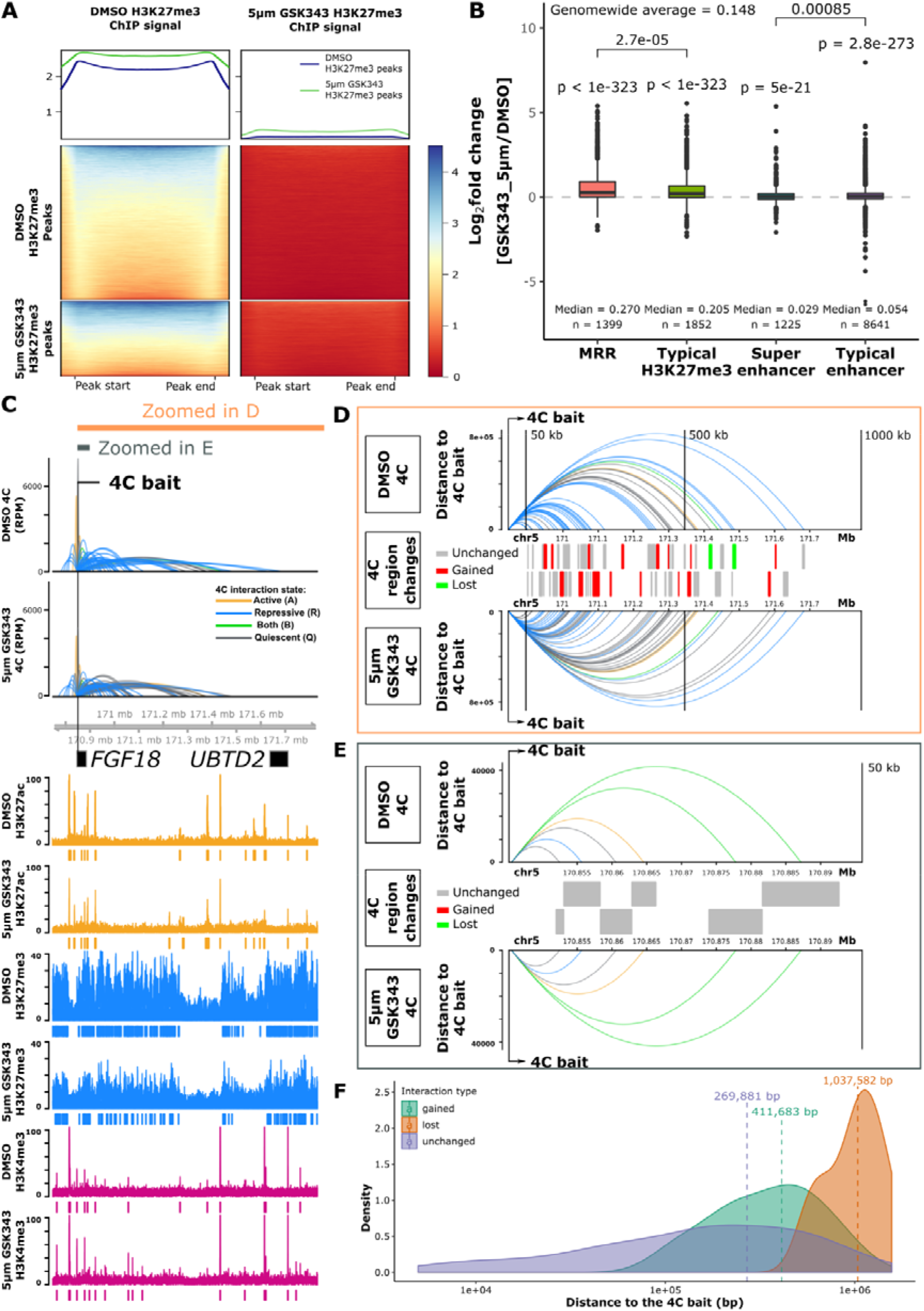
MRR-associated gene expression and chromatin interactions changes after EZH2 perturbation. **A.** H3K27me3 ChIP-seq signal at peaks from DMSO-treated and 5μM GSK343-treated K562 cells. The H3K27me3 ChIP-seq peaks are called using MACS2 in DMSO and GSK343 condition respectively, and then ChIP-seq signal are calculated on these two sets of peaks. The top panel shows the average H3K27me3 signal of H3K27me3 peaks in DMSO and GSK343 condition. The middle panel shows the changes of H3K27me3 signal at the DMSO H3K27me3 peaks. The bottom panel shows the changes of H3K27me3 signal at the GSK343 H3K27me3 peaks. The remaining H3K27me3 peaks in GSK343 condition have higher H3K27me3 levels in DMSO condition. **B.** Expression changes of genes associated with different types of peaks in 5μM GSK343-treated K562 cells. Genes included: 1) Genes with transcription start sites (TSS) overlapped with different peaks; 2) Genes associated with different peaks through Hi-C interaction. One-tail wald test was used for testing significantly up-regulation. All the P values of genes in each category are aggregated. Wilcoxon test p values are indicated, ns: p > 0.05, *: p <= 0.05, **: p <= 0.01, ***: p <= 0.001, ****: p <= 0.0001. **C.** 4C results of *FGF18* in DMSO and GSK343-treated K562 cells. The colors of 4C interactions are based on the distal interacting regions to the 4C bait. Blue: repressive; orange: active; green: both; grey: quiescent. The height of the 4C is shown in Reads Per Million (RPM). The ChIP-seq signal and peaks of H3K27ac, H3K27me3, and H3K4me3 are shown. **D.** Zoomed-in view of 1000kb region downstream of the 4C bait indicated in **C**. Top and bottom panel, 4C interactions in DMSO and 5μM GSK343 conditions. Noted that the y-axis is in distance to the 4C bait. The colors of the 4C interactions are the same as in **C**. Middle panel, detail types of the 4C HindIII fragment. Grey, unchanged 4C regions, which are the 4C interactions that are present in both DMSO and 5μM GSK343 conditions; Red, gained 4C regions, which are the 4C interactions that are only present in 5μM GSK343 condition; Green, lost 4C regions, which are the 4C interactions that are only present in DMSO condition. All the 4C regions are shown in two alternate rows to have a better visual separation. **E.** Zoomed-in view of 50kb region downstream of the 4C bait indicated in **C**. The details of each panel are the same as in **D**. **F.** Density plot of different categories of 4C interactions on the same chromosome as the bait. All the 4C interactions that have p-value < 0.05 on the same chromosome as the 4C bait are included. Gained, 4C interactions present in GSK343-treated 4C but not DMSO-treated 4C; lost, 4C interaction present in DMSO-treated 4C but not GSK343-treated 4C; unchanged, 4C interactions present in both DMSO-treated and GSK343-treated 4C. Mean distances of each category are indicated by the vertical dashed line.

To interrogate the gene expression changes of MRR-related genes, we performed RNA-seq in DMSO-treated and 5μM GSK343-treated K562 cells. The RNA-seq results indicated strong upregulation of H3K27me3-associated genes, while genes associated with H3K27ac peaks (super enhancers or typical enhancers) underwent minimal net change (Figure 8B). Notably, MRR-associated genes were the most strongly upregulated as compared with other categories (typical H3K27me3, super-enhancer and typical enhancers) (Figure 8B). Similarly, a lower dose of 1μM GSK343 treatment in K562 and EZH2 knockout in HAP1 also induced H3K27me3 depletion and significant upregulation of MRR-associated genes as compared with other categories (Figure S7C-E). In addition, cell adhesion related genes in RNA-seq of HAP1 and K562 cells were significantly upregulated (Figure S7F-S3I). This is in concordance with the increased aggregation HAP1 *EZH2* KO cells (Figure S7J). HAP1 *EZH2* KO cells also expressed slower growth rate compared with EZH2 WT cells (Figure S7K), possibly due to contact inhibition of the cells. Taken together, our results showed that MRR-associated genes were highly susceptible to EZH2 inhibition and cell adhesion pathways were upregulated.

To further understand the chromatin interactions changes after EZH2 inhibition treatment, we also performed 4C and ChIP-seq experiments and investigated our candidate genes used in the CRISPR KO experiments in more detail. ChIP-seq data at *FGF18* gene showed that H3K27me3 level was decreased and there were accompanied lost peaks, while the H3K27ac and H3K4me3 signal were mostly unaltered (Figure 8C). By comparing the 4C interactions at *FGF18* promoter in DMSO and GSK343 condition, we found that long-range 4C interactions were altered (Figure 8D), while short-range 4C interactions were almost unchanged (Figure 8E). Density plot showed that the unchanged 4C interactions have a closer distance relative to the 4C bait compared with gained or lost categories (Figure 8F). We also performed 4C experiments in 5μM treated GSK343 K562 cells using MRR1-A1, *IGF2*, and MRR2-A1 as baits, and their interaction profiles showed that short-range interactions are mostly unchanged (Figure S7L-M). To compare the effects of different drug concentrations, we performed all the 4C experiment using same baits (*FGF18*, MRR1-A1, *IGF2*, and MRR2-A1) in 1μM treated GSK343 K562 cells. The 4C interaction profiles in 5μM and 1μM treated GSK343 K562 cells were very similar (Figure S7N).

In addition, we performed 4C experiments using other baits in 1μM treated GSK343 K526 cells and EZH2 KO HAP1 cells which show the same conclusion that the short-range chromatin interactions in the vicinity of the 4C baits were largely unchanged (Figure S7O-R). By contrast, the long-range chromatin interactions tend to change. One question is whether the changing long-range chromatin interactions in H3K27me3 perturbed cells is due to changing numbers of cells displaying chromatin interactions or if it is due to changing chromatin interaction intensities. As the HAP1 cell line is near-haploid, there will just be one copy of a particular gene locus, and a change in the level of chromatin interactions would not be due to changes in the levels chromatin interactions at different alleles. Therefore, every occurrence of a particular chromatin interaction would indicate the presence of one cell, and the number of loops would be equivalent to the number of cells containing the loop in HAP1 cells. As EZH2 KO HAP1 cells showed changing chromatin interactions, we can infer that this is likely to be due to changing numbers of cells that contain such chromatin interactions.

Taken together, these results demonstrated that H3K27me3 perturbation by EZH2 inhibition, either genetically or pharmacologically, can lead to alteration of long-range chromatin interactions.

### Integrative analysis of H3K27me3, H3K27ac and chromatin interactions upon EZH2 inhibition

Since several examples including 4C-seq using *FGF18* promoter as the bait showed long-range chromatin interaction changes upon GSK343 treatment which is consistent with previous MRR2-A1 KO results, we wondered if all the 4C libraries showed the same trend. We classified the chromatin interactions into three categories (short distance, intermediate distance and long distance) based on the distance to bait and found four libraries show the same trend upon 5μM GSK343 treatment that short distance category has higher proportion of unchanged loops (Figure 9A). A similar trend was also observed in 1μM GSK343-treated K562 cells and HAP1 *EZH2* KO cells (Figure S8A). The results of all these libraries strengthen the conclusion that long-range chromatin interactions are susceptible to EZH2 inhibition. It is interesting to note that although EZH2 inhibition and chromatin interaction anchor knockout are two very different types of perturbation experiments, both show that long-range chromatin interactions have a higher tendency to change upon perturbation as compared with short-range chromatin interactions.

**Figure 9.**
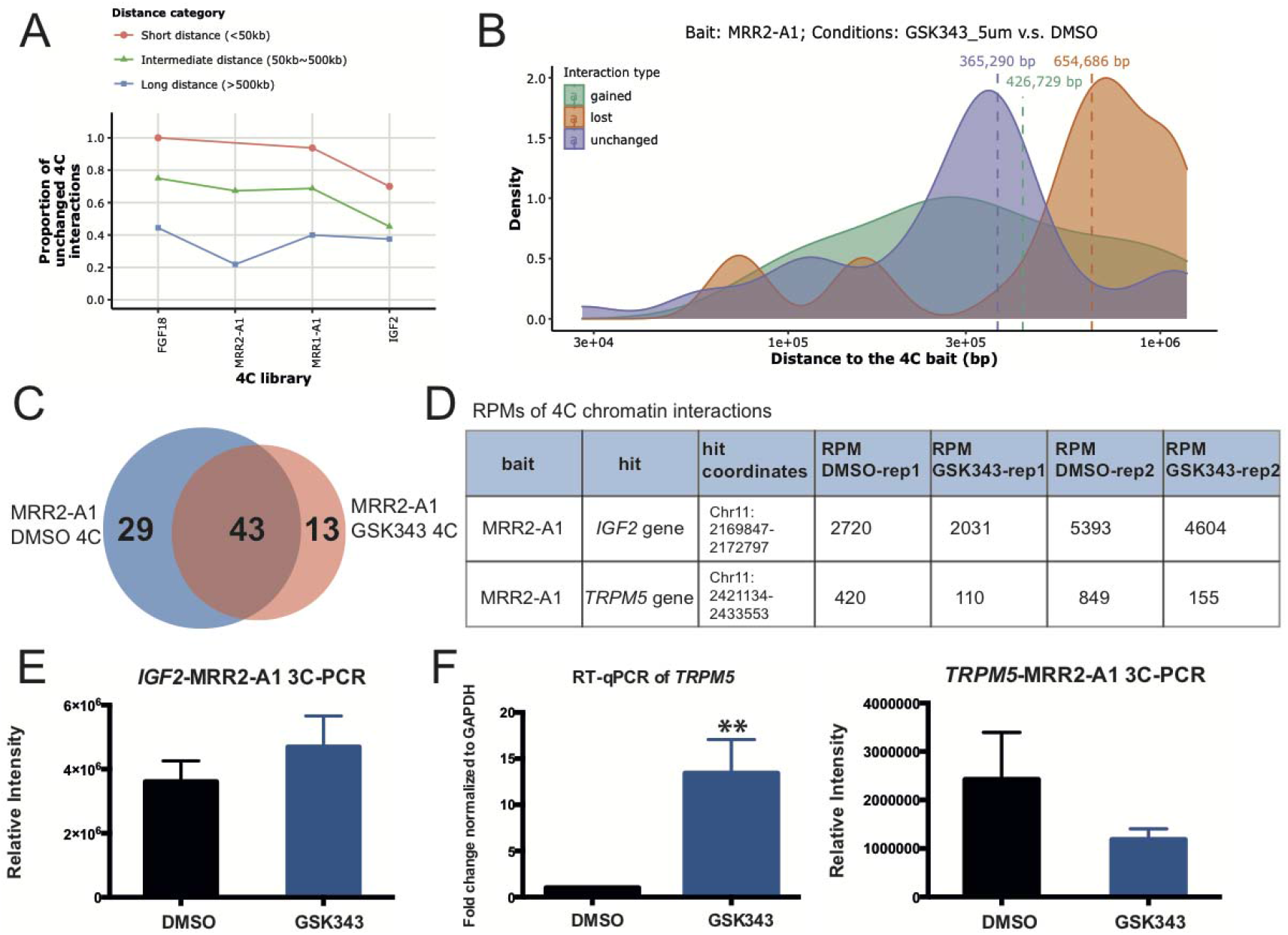
Analysis of stable and changing chromatin interactions upon EZH2 inhibition. **A**. Proportion of unchanged 4C interactions in different distance categories (short, intermediate and long) in 5μM GSK343-treated K562 cells. The bait name is used as the name of the 4C libraries. As the distance of 4C interactions increases, the proportion of unchanged 4C interactions drops, suggesting that long-range interactions are perturbed. **B.** The average distance of changed loops (gained loops and lost loops) is greater than unchanged loops upon GSK343 treatment when using MRR2-A1 as the bait. **C.** Venn diagram of 4C chromatin interactions using MRR2-A1 as the bait in DMSO and GSK343 condition. **D.** Table of Reads Per Million (RPMs) of 4C chromatin interactions in two individual replicates. **E.** 3C-PCR of *IGF2*-MRR2-A1 loop in DMSO and GSK343 condition. The data is shown as relative intensity. **F.** RT-qPCR of *TRPM5* gene and 3C-PCR of *TRPM5*-MRR2-A1 in DMSO and GSK343 condition. All data shown here are average + standard error. P value less than 0.01 is shown as **.

Next, we examined the constant and dynamic chromatin interactions in relation to gene upregulation. We chose the MRR2-A1 region for our EZH2 inhibition analyses in order to compare our results with the MRR2-A1 KO results. 4C-seq data with MRR2-A1 as the bait showed 29 lost loops and 13 gained loops upon GSK343 treatment (Figure 9C). We found the loop to *IGF2* remained unchanged (Figure 9D-E, Figure S8C) while *IGF2* expression was increased (Figure 5E). This phenomenon also observed in MRR1-A1-*FGF18* loop (Figure S8B). We speculated that loss of H3K27me3 at a silencer engaged in stable looping to a target gene promoter will lead to loss of gene silencing at the gene promoter. Next, to investigate changing chromatin interactions, we selected *TRPM5* gene as an example from 29 lost loops (Figure 9C). *TRPM5* gene was significantly upregulated upon GSK343 treatment (Figure 9F). This upregulation was accompanied by disrupted looping to MRR2-A1 which was confirmed by 3C-PCR (Figure 9D-F, Figure S8D). Notably, *TRPM5* gene promoter is more distal than *IGF2* gene promoter in terms of the distance to MRR2-A1 bait, which again supports the conclusion that long-range chromatin interactions tend to change.

As the MRR2-A1 KO example demonstrated that initial histone state is associated with chromatin interactions and silencer KO leads to altered chromatin interactions and histone state which demonstrates interplay between histone modifications and chromatin interactions (Figure 7E), we asked whether EZH2 inhibition by GSK343 will also lead to histone modifications alterations at changing and unchanging chromatin interactions. We performed integrative analysis using MRR2-A1 4C-seq and ChIP-seq as we did for the KO clones (Figure 10A-B). Unlike integrative analysis of MRR2-A1 KO which showed that the initial histone state could predict which chromatin interactions would change (Figure 6D), in the DMSO condition, high H3K27ac levels and low H3K27me3 levels were not associated with unchanged chromatin interactions (Figure S9A).

**Figure 10.**
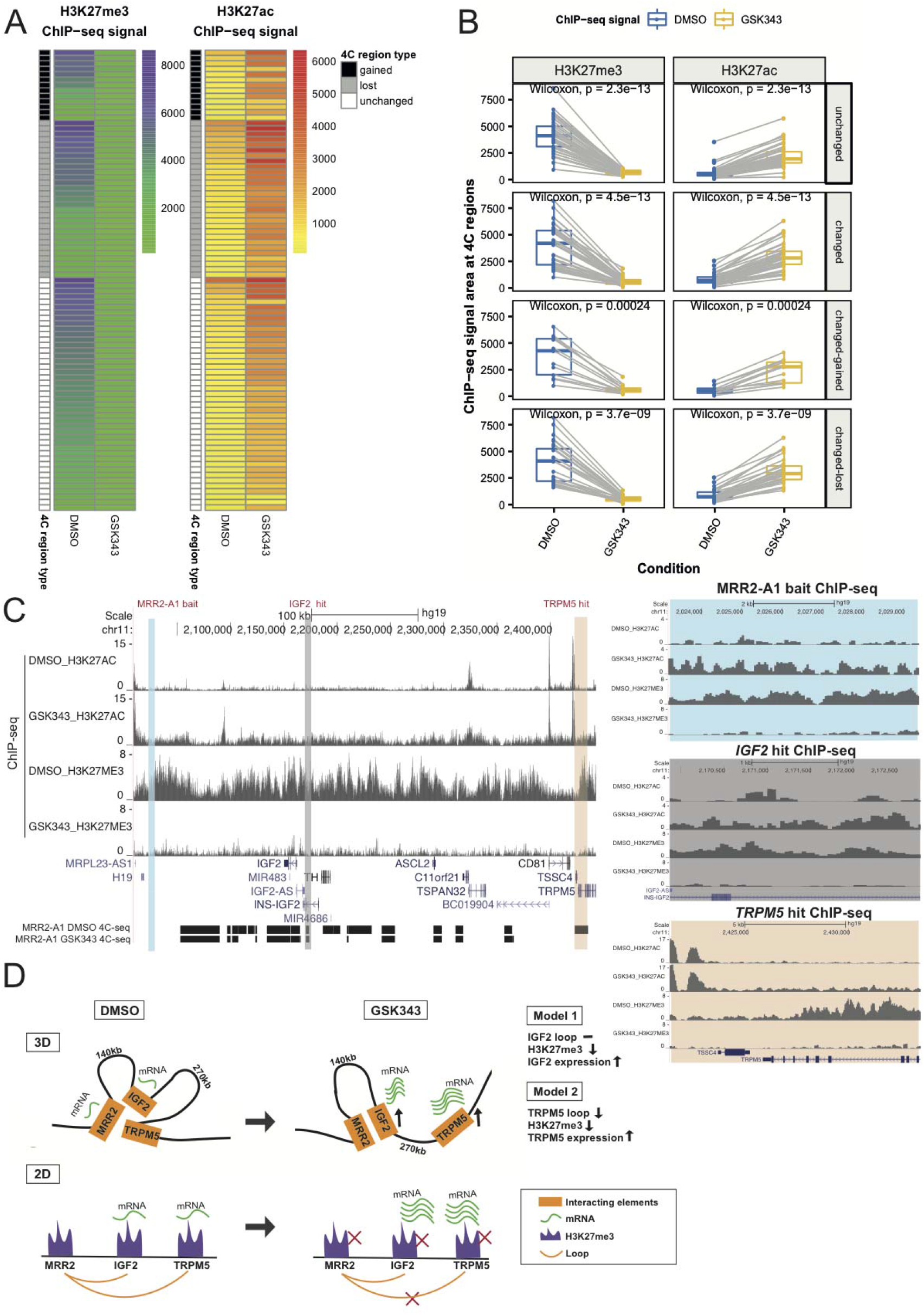
Integrative analysis of H3K27me3, H3K27ac and chromatin interactions upon EZH2 inhibition. **A**. Heatmap of ChIP-seq signal changes of H3K27me3 and H3K27ac at different types of 4C regions (gained, lost and unchanged) in DMSO and GSK343 treated K562 cells. **B.** Boxplots of ChIP-seq signal changes of H3K27me3 and H3K27ac at different types of 4C regions in DMSO and GSK343 treated K562 cells. The same 4C regions are connected by grey lines. Wilcoxon paired test p value are indicated. **C.** Screenshot of H3K27me3 and H3K27ac ChIP-seq at MRR2-A1, *IGF2* gene and *TRPM5* gene regions in DMSO and GSK343 as well as 4C-seq using MRR2-A1 as the bait. MRR2-A1 bait, *IGF2* bait and *TRPM5* bait were highlighted and zoomed in for ChIP-seq. **D.** 3-dimensional and 2-dimensional cartoon schematics of our proposed model involving two mechanisms of how GSK343 leads to *IGF2* gene and *TRPM5* gene upregulation at stable and changing chromatin interactions respectively.

Upon GSK343 treatment, we observed there are global histone modification changes which was shown as decreased H3K27me3 levels and increased H3K27ac levels for all three categories (unchanged, gained and lost loops) (Figure 10A-B), which is consistent with the western blot results (Figure S7A-B). In terms of the two upregulated genes (*IGF2* and *TRPM5*) that we explored before, they both demonstrated loss of H3K27me3 and gain of H3K27ac at the 4C interacting anchors (Figure 10C) although the loop to *IGF2* remained unchanged while the loop to *TRPM5* was reduced. This trend of decreased H3K27me3 and increased H3K27ac was also observed in *FGF18* which showed stable looping with increased gene expression upon GSK343 treatment (Figure S9B).

Taken together, the integrative analysis combined with the 3C-PCR showed two models regarding how EZH2 inhibition leads to target gene upregulation (Figure 10D). Model 1 showed decreased H3K27me3 levels with stable loop upon GSK343 treatment which applies to both *IGF2* gene and *FGF18* gene. However, we noticed differences between 4C-seq using the *IGF2* promoter as the bait and 4C-seq using the *FGF18* promoter as the bait (Figure S9C-D). Specifically, there are many repressive loops lost at the *IGF2* promoter while there are only a few repressive loops lost at the *FGF18* promoter in GSK343 condition which may explain the differences in gene upregulation upon GSK343 treatment (Figure 3E, Figure 5E). Model 2 showed decreased H3K27me3 levels with disrupted loop upon GSK343 treatment as observed with the *TRPM5* gene (Figure 10D). Therefore, in terms of the relationship between H3K27me3, chromatin interactions and gene upregulation, we think that there are two different models. The first model is that long-range chromatin interactions facilitate the deposition of H3K27me3 modification onto the target gene promoter by MRRs to repress target genes (e.g. *IGF2* and *FGF18*). The second model is that depletion of H3K27me3 abrogates long-range chromatin interactions, which in turn cause upregulation of target genes (e.g. *TRPM5*).

## Discussion

Silencers are important regulatory elements for gene regulation, and several studies have suggested that they loop to target genes, in a manner analogous to enhancers. Although there are several examples of proposed silencers that have been experimentally validated (Table S1) and several methods have been proposed to identify silencer elements (Table S2), however, there is however no consensus on their foolproof identity yet. Additionally, except several PRC2-bound silencers in mouse^13^ no silencers that work via a looping mechanism have been characterized yet. Here, we propose a new method to identify H3K27me3-rich regions (MRRs) or putative “super-silencers” through clustering and ranking H3K27me3 signals.

We found that MRRs are highly associated with chromatin interactions and can be perturbed by EZH2 inhibition. Through H3K27me3 clustering, ranking and associate them with chromatin interactions, we validated two looping silencer examples (MRR1-A1 and MRR2-A1). We showed that silencer removal cause cell identity changes and further related to tumor growth inhibition. Moreover, MRR2-A1 example demonstrated that silencer removal will cause changes of long-range chromatin interactions and high H3K27ac loops were gained to activate *IGF2* gene expression.

The mechanism of how silencers function to repress genes will be an interesting topic to explore. Through the *IGF2* silencer example, we showed that that looping silencer removal causes distant loops to change and histone states in the initial conditions can predict loop changes. Importantly, we found that loops with high H3K27ac and low H3K27me3 tend to change, which provides evidence that histone modifications can affect overall genome architecture. Secondly, we found that short-range loops tend to remain unchanged while long range loops are disturbed either upon silencer KO or GSK343 treatment which is in line with the finding that showed PRC1 and PRC2 are necessary to maintain the chromatin interactions landscape^17,61^. Thirdly, there are multiple regions inside an MRR that are involved in chromatin interactions and may also function as silencers. It would be interesting to see whether the putative silencers in an MRR function similarly or differently and to dissect different functional mechanisms of silencers. Fourthly, transcription factors can contribute to the chromatin interaction landscape and cell type-specific transcription factors may result in different chromatin interaction landscape^62^. Therefore, elucidating the transcription factors involved in silencer functioning would be an important future direction for research.

Another interesting question would be the importance of H3K27me3 at MRR and H3K27me3 on target gene promoters. In our CRISPR knockout results, we observed that excision of a distally interacting MRR region (MRR2-A1) to *IGF2* can lead to *IGF2* upregulation, which indicated the repressive ability of H3K27me3 at distal MRR. Notably, the residual H3K27me3 on the *IGF2* gene promoter (Figure 5C) still showed a repressive effect even after excision of MRR2-A1. However, we still cannot infer the relative importance of H3K27me3 at MRR and H3K27me3 on target gene promoters. We also cannot exclude the possibility that there are other distal silencers controlling gene expression. This question could be further addressed by more targeted perturbation of H3K27me3 at both or either end of the chromatin interactions.

Our pharmacological and genetic inhibition of EZH2 showed that MRR-associated genes as well as long-range chromatin interactions were susceptible to the depletion of H3K27me3 histone marks. MRR-associated genes were more susceptible to the depletion of H3K27me3 marks than genes associated with typical H3K27me3 peaks. This result suggested that the response of different genes to H3K27me3 loss may correlate with their chromatin state. Although differences in chromatin interactions have been observed in cells at different developmental stages^58,59^, whether chromatin interactions can be affected by histone modifications and such perturbations is still an open question. A study in human fibroblast cells showed that the contacts between enhancers and promoters were present in the cells before the transient treatment of TNF-α^60^, suggesting a pre-existing and stable chromatin architecture. However, we observed that long-range chromatin interactions are susceptible to EZH2 inhibition or knockout, which is consistent with previous studies showing that PRC1^18,63^ and PRC2 complexes^63^ are necessary for long-range chromatin interactions formation. Our results are different from these previous studies in that we dived into more details by classifying the loops into different categories and performing plenty of different 4C libraries in both GSK343 condition and KO condition. Furthermore, through 4C and ChIP-seq integrative analysis, we revealed the interplay between histone modifications and changing chromatin interactions. Taken together, our results suggest that regulatory elements at a great distance can be brought into proximity to genes and form a permissive or repressive microenvironment around genes to help regulate their expression. To address the relationship of histone modifications and chromatin interaction formation, further investigation of the structural differences in response to different cell treatments should be done using high-resolution and whole-genome chromatin interactions sequencing methods. This can help us to understand the mechanisms of gene activation or repression in cellular pathways.

We and other people found that silencers are cell-type specific and highly context-dependent^27–29^ (Table S2). Specifically, the same genomic region was a silencer in one cell line but a super-enhancer in another cell line. Not surprisingly, this change is associated with different gene expression in the different cell lines. Moreover, it has been shown that silencers can transit into active enhancers during differentiation^13^. Thus, the study of relationships between cell types and silencers can shed light on cell type specific regulation of gene expression. The mechanism by which important oncogenes such as *TERT* are silenced in normal cells is unclear. It would be interesting to investigate whether *TERT* is regulated by MRRs that transit into active enhancers in cancer cells.

Interestingly, we found that silencer removal leads to cell differentiation and tumor growth inhibition, which is in line with previous observed studies that showed that more H3K27me3 can render Topologically-Associated Domains (TADs) inactive and repress tumor suppressor genes^61^. It will be interesting to study the detailed mechanism of how silencers regulate tumor suppressor genes. In this way, it may be possible for us to activate the tumor suppressor genes expression by perturbing silencers, just as super-enhancer perturbation can result in loss of oncogene expression^32^.

Notably, the question of whether super-enhancers are indeed different from enhancers is not settled yet^64^. Our research raises similar questions: are MRRs, which are potentially “super-silencers”, different from typical silencers? The regions of the long MRR that are critical for silencer function are not fully elucidated yet. Here we showed that the components of the MRRs that are involved in looping interactions are important in repressing long-range chromatin interactions, while the roles of other components of the MRRs are not yet known. Moreover, we found that different anchors within the same MRR can be associated with different proteins, suggesting that these different anchors may play different roles within the MRR. Detailed dissection of the different anchors and other components of MRRs will be required to answer these questions in future work.

Moreover, it will be interesting to explore how looping is mediated at MRRs. Given that super-enhancers have been shown to be involved in phase condensation^65^, and HP1 which is a component of constitutive heterochromatin associated with H3K9me3 has also been shown to be able to form phase condensates^66^, it would be interesting to explore whether PRC2 complex and H3K27me3 can give rise to phase condensates.

In conclusion, maintenance of cellular identity requires that the right genes are expressed and other genes are silenced. Distal looping silencers have been well explored in *Drosophila* and mice^67^ but there are no known examples in human. Our results add to the understanding of silencers by identifying silencer elements in human and demonstrating the existence of looping silencers in human. Just as the concept of “super-enhancers” has been useful in identifying oncogenes and therapeutic vulnerabilities in cancer cells, the concept of silencers calling by clustering of H3K27me3 may be useful in identifying genes involved in controlling cellular identity and cancer progression.

## Methods

We performed Hi-C interaction analysis, ChIP-seq, RNA-seq, gene expression analyses, cell culture, RT-qPCR, CRISPR excision, 4C, 3C, xenograft models, western blot, adhesion assays, and growth curves as described in the **Supplementary Methods**. A list of all libraries used and generated is provided in **Supplementary Table S3**. A list of all the primers used is provided in **Supplementary Table S4**.

## Acknowledgements

We would like to thank all members of the Fullwood Lab and Ah Jung Jeon for helpful comments. This research is supported by the National Research Foundation (NRF) Singapore through an NRF Fellowship awarded to M.J.F (NRF-NRFF2012-054) and NTU start-up funds awarded to M.J.F. This research is supported by the RNA Biology Center at the Cancer Science Institute of Singapore, NUS, as part of funding under the Singapore Ministry of Education Academic Research Fund Tier 3 awarded to Daniel Tenen as lead PI with M.J.F as co-investigator (MOE2014-T3-1-006). This research is supported by a Singapore MOE Academic Research Research Fund (T1) grant to G.T-K. This research is supported by an National Research Foundation Competitive Research Programme grant awarded to V.T. as lead PI and M.J.F. as co-PI (NRF-CRP17-2017-02). This research is supported by the National Research Foundation Singapore and the Singapore Ministry of Education under its Research Centres of Excellence initiative.

## Author contributions

Y.C.C., Y.Z., M.J.F. and G.T-K. conceived of the research. Y.C.C., Y.Z., M.J.F. and G.T-K. contributed to the study design. Y.C.C. performed bioinformatics analysis. Y.Z. and S.L. designed CRISPR knock out experiments. Y.Z. performed CRISPR knock out, 4C, 3C-PCR, RNA-seq, ChIP-seq, ChIP-qPCR and other functional experiments for KO clones. Y.Z. and Y.P.L. performed EZH2 inhibitor and HAP1 *EZH2* knockout experiments and 4C experiments. E.L-A. advised on the interpretation of EZH2 inhibitor results. J.Q.T performed ChIP-seq and ChIP-qPCR experiments for HAP1 *EZH2* KO cells. Z.C. and M.Q.L. performed 4C experiments. A.R., L.M. and V.T. designed xenograft experiments. A.R. performed xenograft experiments. Y.C.C., Y.Z., M.J.F. and G.T-K. reviewed the data and wrote the manuscript. All authors reviewed and approved of the manuscript.

## Data deposition

The list of libraries used in the study is provided in Table S3. All datasets have been deposited into GEO.

## Author information

The authors declare that they have no competing interests.

Correspondence and requests for materials should be addressed to.

