## Supplementary Materials for "H3K27me3-rich genomic regions can function as silencers to repress gene expression via chromatin interactions"

**Contents:**

1. **Supplementary Methods**
2. **Supplementary Text**
3. **Supplementary Figure Legends**

**4. Supplementary Tables**

**5. Supplementary Figures**

**6. Supplementary References**

**Supplementary Methods**

**Definition of H3K27me3-rich regions (MRRs)**

H3K27me3 ChIP-Seq signal and peaks were obtained from ENCODE and used as inputs of an in-house customized script that mimicked the signal calculation of the ROSE package. Missing H3K27me3 peaks from ENCODE were called using MACS2 (2.1.0.20150731)^1^ with pooled replicates using option "--broad -q 0.05". First, ChIP-Seq peaks of H3K27me3 were stitched using a window size of 4 kb. After stitching, the treatment and control signal of the stitched peaks were calculated and used in the ranking in MRR calling. Super-enhancers were called in a similar manner except that a stitching window of 12.5 kb was used.

**Cell culture**

GM12878 normal lymphoblastoid cell line and Chronic Myelogenous Leukemia cell line K562 were cultured in RPMI-1640 supplemented with 10% Fetal Bovine Serum (FBS) and 1% penicillin-streptomycin. HAP1 cell line is a haploid human cell line that was derived from KBM7 cells^2^. HAP1 wildtype and EZH2 knockout cells (purchased from Horizon) were cultured in IMDM supplemented with 10% FBS and 1% penicillin-streptomycin. All cultures were maintained at 37 °C, 5% CO2 in a humidified incubator.

**Circular chromosome conformation capture (4C)**

4C-seq assays were performed as previously described^3^ with slight modifications. Briefly, 4 × 10^7^ cells were cross-linked with 1% formaldehyde. The nuclei pellets were isolated by cell lysis with cold lysis buffer (10mM Tris-HCl, 10mM NaCl, 5mM EDTA, 0.5% NP 40) supplemented with protease inhibitors (Roche). First step digestion was performed overnight at 37°C with HindIII enzyme (NEB). Digestion efficiency was measured by RT-qPCR with HindIII site-specific primers. After confirmation of good digestion efficiency, DNA was ligated overnight at 16°C by T4 DNA ligase (Thermo Scientific) and de-crosslinked. Following de-crosslinking, DNA was extracted by phenol-chloroform and this is the 3C library. The DNA was then processed for second digestion with DpnII enzyme (NEB) overnight at 37°C. After final ligation, 4C template DNA was obtained, and the concentration was determined using Qubit assays (Thermo Scientific). The 4C template DNA was then amplified using specific primers with Illumina Nextera adapters and sent for sequencing on the MiSeq system. All the 4C genome coordinates are listed in Table S5.

**3C-PCR**

The 3C libraries were generated as described in the 4C section. Digestion efficiency was checked by gel electrophoresis and concentration was determined by Qubit assay. The Taq PCR core kit (Qiagen) was used for PCR reactions with 600ng 3C library template using the following protocol: 98°C 3 min, 33 cycles [94°C 1 min, 60°C 1 min, 72°C 20 sec], 72°C 10 min. PCR products were run on 1.5% agarose gels. After gel electrophoresis, bands corresponding to the expected products were gel excised (Qiagen) and purified for Sanger sequencing. The intensities of the bands were measured by Image Lab. Primers were designed for 3C-PCR following the unidirectional strategy^4^. Primers used are listed in Table S4. At least two replicates were performed for 3C analyses.

**ChIP-Seq and ChIP-qPCR**

ChIP-seq was performed as described previously^5^ with slight modifications. Briefly, cells were crosslinked with 1% methanol-free formaldehyde (Thermo Scientific) at room temperature for 10 minutes, followed by quenching with glycine for 5 minutes at room temperature. The fixed cell pellet was lysed in 1% SDS lysis buffer supplemented with protease inhibitor cocktail tablet (Roche), and sonicated using Bioruptor (Diagenode).

The cell lysate was precleared through centrifugation in dilution buffer and incubation with Protein G Dynabeads (Invitrogen) overnight at 4°C. The precleared lysate was added into the prepared antibody-conjugated beads, and incubated overnight at 4°C, with rotation. The beads were then washed thrice in 0.1% SDS lysis buffer, once in high salt wash buffer, once in lithium chloride wash and once in TE buffer. The beads were eluted in elution buffer treated with RNase A (Qiagen) followed by decrosslinking with Proteinase K (Ambion) at 37°C overnight. ChIP DNA was cleaned up with QIAquick PCR purification kit (Qiagen), and quantitated using Qubit High Sensitivity dsDNA Assay (Invitrogen).

ChIP DNA was used for the construction of DNA library for Illumina HiSeq 4000 NGS sequencing using ThruPLEX DNA-seq 48D Kit (Rubicon) according to the instruction. Antibodies used include H3K4me3 (#ab8580, Abcam), H3K27me3 (C36B11, Cell Signaling Technologies), H3K27ac (#ab4729, Abcam) and mouse IgG (#sc-2025, Santa Cruz).

ChIP-qPCR reactions were performed in triplicates on a Quantstudio 5 quantitative PCR machine (Life Technologies) using GoTaq qPCR Master Mix (Promega). Primers used are listed in Table S4.

**EZH2 inhibitor treatment**

Small molecular inhibitor GSK343 (Sigma-Aldrich) targeting EZH2 were solubilized in DMSO (Sigma-Aldrich) according to manufacturer’s instructions and used at a final concentration of 5μM. K562 cells with GSK343 or DMSO vehicle control were incubated at 37 °C, 5% CO2 humidified incubator for 48 hours before harvesting for various experiments.

**RNA extraction and RT-qPCR**

Total RNA were isolated from the cells using RNeasy Mini Kit (Qiagen) with on-column DNase digestion (Qiagen). 1ug of total RNA was then reverse transcribed to cDNA using the SuperScript III first-strand synthesis system using oligodT (Invitrogen). The expression levels of various genes were analysed by real-time PCR. Quantitative real-time PCR (qPCR) was performed using the Applied Biosystems QuantStudio 3 Real-Time PCR system using SYBR Green PCR Master Mix and appropriate primers. Primers used are listed in Table S4 and tested by plotting the standard curve for various dilution ratios. We only accepted the primers whose efficiency are between 80%-120%. The transcript levels of genes were analysed by 2^−ΔΔCt^ method^6^.

**RNA-Seq**

Total RNA was extracted from cells using RNeasy Mini Kit (Qiagen), with on-column DNase I treatment (Qiagen). The quality of the RNA extracted was checked using Agilent RNA 6000 Nano Kit (Agilent), and quantitated using Nanodrop ND1000 (Thermo Scientific). 850ng of RNA was used for the construction of cDNA library for Illumina HiSeq 2500 High Output v4 NGS sequencing using TruSeq Stranded Total RNA LT (w/ Ribo- Zero Gold) Set A (Illumina) as per protocol.

**Protein extraction and western blot**

Proteins were extracted from the cells using RIPA buffer (Sigma-Aldrich) with protease inhibitor cocktail (Life Technologies). Protein concentrations were determined using BCA assay (Thermo Scientific). 20μg of proteins were separated in 4-20% Mini-PROTEAN® TGX™ Precast Gels (Bio-Rad) and transferred to PVDF membrane. After blocking with TBST containing 5% nonfat dried milk for 1h at room temperature, the membrane was washed twice with TBST and incubated with primary antibodies: EZH2 (Cell Signaling Technology AC22), TBP (abcam ab51841), beta-Actin (abcam ab6276), total H3 (abcam ab1791), H3K27me3 (Cell Signaling Technology C36B11), H3K27ac (abcam ab4729) and H3K9me3 (abcam ab8898) overnight at 4°C. The membrane was washed three times with TBST for 10min and then incubated for 1h at room temperature with HRP-conjugated secondary antibodies (Cell Signaling Technology). After extensive washing, bands were detected by enhanced chemiluminescence reagent (Bio-Rad) and imaged using the ChemiDoc™ imaging system (Bio-Rad).

**CRISPR excision**

CRISPR excision was performed as described previously^7^ using all-in-one CRISPR/Cas9 vector system. Briefly, gRNAs were designed using Zhang Feng’s website (http:// CRISPR.mit.edu)^8^ and two gRNAs were designed for each region. Single gRNA was cloned into either pX330A/pX330S vector (gift from Li Shang, pX330A modified to include GFP reporter marker) followed by Sanger sequencing to confirm insertion of single gRNA. Golden gate assembly of two gRNAs was performed as described previously. Positive two gRNAs insertion plasmids were confirmed by Sanger sequencing using CRISPR-step2-F and CRISPR-step2-R primers (sequences were shown in Table S4). The plasmid was then electroporated into the K562 cell line using the Neon transfection system (Thermo Fisher). After 48 hours, transfected cells were FACS sorted into 96-well plates as single-cell colonies based on GFP signal.

Cells were harvested from each clone, pelleted and lysed in lysis buffer. Genotyping was carried out using an internal and a flanking primer pair. Final PCR products were imaged by agarose gel electrophoresis, and successful clones were confirmed through Sanger sequencing (First Base).

**Adhesion assay**

Cell adhesion assay was performed using CytoSelect 48-Well Adhesion Assay (Cell Biolabs, San Diego, CA) according to the manufacturer’s protocol. Briefly, a cell suspension (5X10^5^ cells/200 ml FBS-free medium) was added to fibronectin coated wells and BSA coated wells (negative control) respectively. After 3 hours of incubation, cells were washed with PBS, stained with crystal violet and then eluted with extraction solution. The levels of adhesion were quantified by optical absorbance at 560 nm using the Tecan plate reader.

**Growth curve assay**

1000 cells/well were seeded in 96 well plates and cell growth was measured at day0, day1, day2, day4 and day5 using the CellTiterGlo assay kit (Promega, G7571). Luminescence was measured on a Tecan plate reader.

**SiRNA knock down experiment**

SiRNAs (Thermo Fisher) were introduced into cells using Neo transfection system (Thermo Fisher) according to the manufacturer’s instructions. *FGF18*-siRNA-1, 5’-GAGACGGAAUUCUACCUGUtt-3’, *FGF18*-siRNA-2, 5’-AGACACCUUCGGUAGUCAAtt-3’, *IGF2*-siRNA-1, 5’-CCAUGCAAAUGAAAUGUAAtt-3’ *IGF2*-siRNA-2, 5’-GGAAGCACAGCAGCAUCUUtt-3’ were used for RNA interference. After 48h incubation, cells were harvested for RT-qPCR. Primers used were listed in Table S4.

**Xenograft experiment**

All the animal studies were carried out in accordance with animal care and use guidelines approved by Biological Resource Centre, Singapore.

Female CB17 SCID mice (6-8 weeks old) were used for the present study. The mice (n=5) were injected subcutaneously. All the mice were monitored for tumor growth at the site of inoculation and the tumor volume was measured twice a week using Vernier caliper for 40 days or till the tumor volume reaches 1000 mm^3^ whichever is earlier. The tumor volume was calculated using the following formula V = a × b^2^ × 0.52, where a is the largest and b the smallest diameter of the tumor. At the end of the experimental period, the tumors were resected out and each tumor piece was then divided into 2 pieces. One piece of the tumor was fixed in 10% NBF for 24 hours at room temperature and then paraffin embedded for H&E & IHC analysis. The other piece was snap frozen for RNA and protein analysis.

KO-1 and EV were injected into separate mice in Experiment 1. In view of the concern that the differences in tumor growth seen in Experiment 1 might be because the KO-1 and EV were grown in different mice, in experiment 2, KO-2 and EV were injected in the same mice (left side and right side).

**Data sources**

Processed Hi-C interactions in K562, GM12878, and HAP1 were obtained from GEO (GSE63525). H3K27me3 and H3K27ac ChIP-Seq peaks in K562 and GM12878 obtained from ENCODE at UCSC (wgEncodeEH000031, wgEncodeEH000044, wgEncodeEH000030, wgEncodeEH000043). Other H3K27me3 ChIP-Seq data in H1hESC, HeLaS3, HepG2, and MCF7 are obtained ENCODE (ENCSR000ALU, ENCSR000APB, ENCSR000AOL, and ENCSR768LHG). H3K27me3 ChIP-Seq data in KARPAS-422, Pfeiffer, and WSU-DLCL2 were obtained from GEO (GSE40970). EZH2 ChIP-Seq data in K562, GM12878, H1hESC, HepG2, and HeLaS3 were obtained from ENCODE (ENCFF083IDB, ENCSR000ARD, ENCSR000ASY, ENCSR000ARI, and ENCSR000ATC).

**Visualization of chromatin interactions, and ChIP-Seq data**

Hi-C and 4C interactions were drawn in arc style using Sushi (1.16.0)^9^ from Bioconductor. The heights of Hi-C and 4C were highest read counts at Hi-C interacting regions and RPM at 4C interacting regions, respectively. The colors of Hi-C and 4C interactions were decided by the state on the distal interacting regions relative to gene TSS or 4C baits, respectively. Blue: repressive; orange: active; green: both; grey: quiescent. Tracks of ChIP-Seq signal and peaks were generated by Gviz (1.22.3)^10^ from Bioconductor.

**Feature enrichment, gene ontology and pathway enrichment analysis**

Genomic feature enrichment analysis was performed using R package annotatr^11^. Gene ontology, pathway enrichment analysis (REACTOME & KEGG) and map-view representation of enriched pathways were performed using R package clusterProfiler^12^ and ReactomeRA^13^.

**Gene expression specificity**

Cell line expression data were obtained from Epigenetic RoadMap (K562, GM12878, H1hESC, HeLaS3 and HepG2)^14^, CCLE (KARPAS, Pfeiffer, and WSUDLCL2)^15^. Gene expressions in each cell line were compared with 69 facets of annotated CAGE clusters with normalized expression data from FANTOM5^16^. The grouping facets were obtained from Andersson *et al*^17^*.* The average expression of all samples in a facet was assigned to that facet. For each cell line, gene expression was considered as 1 facet and combined with the other 69 facets from FANTOM5 to form 70 facets in total. Gene expression specificity was calculated on these 70 facets for each cell line independently. The specificity of each gene X is calculated by: Specificity(X)=1−(entropy(X)/log2(N)) where X is the vector of expression values of the cluster in across all facets, and N = |X|. The definition of specificity is identical to that used by Andersson *et al*^17^*.* Quartile Q1 and Q3 are used as cut-off of 'Low specificity' and 'High specificity', respectively.

**RNA-Seq, ChIP-Seq & 4C data analysis**

RNA-Seq reads of HAP1 EZH2 KO/WT and K562 GSK343/DMSO were analyzed with kallisto (0.44.0)^18^ with option '-b 100'. Differentially expressed genes were called using sleuth (0.29.0)^19^ with gene-level aggregation and wald test. ChIP-Seq reads of H3K2me3, H3K27ac and H3K4me3 from HAP1 WT and EZH2 KO clones were mapped by BOWTIE2 (v2.2.5)^20^ using default parameters in pair-end mode and filter out alignment with a mapq score smaller than 30. The two replicates were combined and peaks and bigWig files were generated by MACS2 (2.1.0.20150731)^1^ using option '-q 0.01' for H3K27ac and H3K4me3 and '--broad --broad-cutoff 0.1 -q 0.05' for H3K27me3. 4C reads were trimmed off HindIII digestion site using tagdust (2.33)^21^ and only those remained paired were mapped by BOWTIE2 (v2.2.5)^20^ with option '--end-to-end' in single-end mode. R3Cseq (1.24.0)^22^ was used to call significant interactions against Hind III digested genome background with a cut-off p value of 0.05. Two replicates were performed for each 4C analysis, and significant interactions of two replicates were pooled.

**Transcription factor binding enrichment analysis**

ChIP-seq peaks of CTCF (ENCFF738TKN), RAD21 (ENCFF002CXU), SMC3 (ENCFF041YQC), REST (ENCFF895QLA), ZNF143 (ENCFF114IWY), EZH2 (ENCFF083IDB), GATAD2B (ENCFF549KOD), and YY1 (ENCFF557DSM) from K562 were downloaded from ENCODE. Interacting regions of MRRs were generated by overlapping MRRs with Hi-C loops. The number of overlapping ChIP-seq peaks was calculated by overlapping interacting regions with different TF. These numbers were then standardized into Z-score across all interacting regions and subjected to heatmap clustering (<https://CRAN.R-project.org/package=pheatmap>)^44^.

**Analysis of changing as compared with unchanging chromatin interactions**

All the 4C interactions that passed the threshold of p-value < 0.05 in each 4C library were used. The 4C interactions in different conditions were compared to the control condition (either HAP1 WT or DMSO-treated K562). After that, 4C interactions were classified into gained, lost, or unchanged. Gained, 4C interactions present in the experimental condition but not in the control condition; lost, 4C interaction present in the control condition but not in the experimental condition; unchanged, 4C interactions present in both control and experimental conditions. The proportions of unchanged 4C interactions in different distance categories were calculated as unchanged 4C interactions divided by the total number of 4C interactions in that distance category. Categories with less than 3 4C interactions were excluded in this analysis.

**Heatmap and boxplot of RPM signal and ChIP-seq signal at different 4C regions**

4C regions are classified as gained, lost, and unchanged according to their presence in experiment versus control condition. For GSK343-treated K562 4C data, the experiment condition is GSK343-treated K562, while the control condition is DMSO-treated K562. Similarly, for CRISPR KO K562 4C data, the experiment condition is either *FGF18*-MRR1-A1 KO or *IGF2*-MRR2-A1 KO clones, while the control condition is the EV clones. Gained, 4C interactions only present in the experiment condition but not in the control condition; Lost, 4C interactions only present in the control condition but not in experiment condition; unchanged, 4C interactions present in both control and experiment conditions. In this comparison, only those 4C interactions with p-value < 0.05 are considered.

For RPM signal heatmap, different types of 4C regions (gained/lost/unchanged) are first classified according to their H3K27ac/H3K27me3 ChIP-seq signal levels in the control condition. Tertiles are used to classify these 4C regions into high, medium, or low category. The 4C intensity in RPM of each 4C regions are shown in a color-scaled manner.

For ChIP-seq signal heatmap, ChIP-seq signal different types of 4C regions (gained/lost/unchanged) are calculated as the area of signal at these regions. Deeptools computeMatrix^45^ is used to calculated ChIP-seq signal at these 4C regions, and then the total signal area are calculated as: Total Signal Area = sum (Sig *BS), where BS is the size of bins used when summarizing RPKM, and Sig is the ChIP-seq signal in RPKM at individual bin. For ChIP-seq signal boxplot, the same 4C region (gained/lost/unchanged) in different conditions are connected using grey line. Wilcoxon paired test are used, and p value are indicated accordingly: ns: p > 0.05, *: p <= 0.05, **: p <= 0.01, ***: p <= 0.001, ****: p <= 0.0001.

**Supplementary Text**

**Selection of regions to CRISPR**

We found that there are 974 MRRs in K562 cell line and their median size is 92,170bp. We aligned these MRRs with Hi-C anchors and found that there are 560 MRRs associated with interactions. Next, through aligning with gene annotation, we found that there are 237 MRRs which are associated with genes.

These MRRs can be further classified into three different categories: (1) proximal looping (MRR overlap with target gene promoter), (2) distal looping (MRR loops over to the promoter of target gene while not overlap with its promoter) and (3) internal looping (MRR loops over to the promoter of target gene and at the same time overlap with gene promoter). There are 54 internal looping MRRs and since we want to investigate the relationship between H3K27me3, chromatin interactions and gene expression, we chose CRISPR examples from this category.

Firstly, we filtered out some known translocated regions and rearrangements such as *BCR-ABL* and the *IGHV* cluster. Next, we selected 13 MRRs whose H3K27me3 signal rank is lower than 80 and loop number is equal or larger than 10. Among those MRRs, we further chose those contain genes which loop to more than one anchor. Finally, we selected two MRRs associated with cell identity including MRR1 whose target gene is *FGF18* and MRR2 whose target gene is *IGF2* (Figure S3A).

**Supplementary Figure Legends**

**Figure S1**. Identification of H3K27me3-rich regions (MRRs).

**A.** Number of constituent peaks in typical H3K27me3 peaks, MRRs, typical enhancers, and super-enhancers in K562, GM12878, and HAP1 cells, respectively. **B.** Number of overlapping genes at typical H3K27me3 peaks, MRRs, typical enhancers, and super-enhancers in K562, GM12878, and HAP1 cells, respectively. **C.** Proportion of constituent peaks of typical H3K27me3 and constituent peaks of MRR that were overlapped with CpG island. All, all the constituent peaks including typical H3K27me3 peaks and MRRs; random regions, randomly shuffled regions of all the constituent peaks. **D.** Proportion of constituent peaks of typical H3K27me3 and constituent peaks of MRR that were overlapped with different gene features. All and random regions were generated as described in **Figure S1C**. **E.** H3K27me3-rich regions (MRR) and typical H3K27me3 peaks and their associated genes in K562, GM12878, HAP1, and H1hESC. One of the genes that had TSS overlapped with top 10 MRR were shown here. TSG, predicted tumor suppressor genes by TUSON. **F & G.** Gene ontology analysis of gene associated with MRR (F) or typical H3K27me3 (G) in K562, GM12878, HAP1 and H1hESC. **H & I.** H3K27me3, H3K27ac, and EZH2 ChIP-seq signal on typical H3K27me3, MRR, constituent peaks of typical H3K27me3 peaks, and constituent peaks of MRRs in GM12878 and HAP1. Peaks were scaled to the same median length of peaks in typical H3K27me3, MRR or constituent peaks, and the ranges were expanded by 5kb on both sides of the peaks. **J & K.** EZH2, SUZ12, and BMI1 ChIP-seq signal on typical H3K27me3, MRR, constituent peaks of typical H3K27me3 peaks, and constituent peaks of MRRs in K562 and GM12878. **L.** Example of CPED1 and DENND2D and their associated MRR/SE in K562 and GM12878 cell lines. MRR and SE could be interchangeable in different cell lines. SE, super enhancers; MRR, H3K27me3-rich regions. Expression level of CPED1 is 107.826 and 0.029 in K562 and GM12878, respectively; expression level of DENND2D is 0.002 and 78.004 (expression in RPKM). **M & N.** Examples of CPED1 and DENND2D and their associated MRR/SE in K562 and HAP1 cells. SE, super enhancers; MRR, H3K27me3-rich region. Expression level of CPED1 is 107.826 and 0.67 in K562 and HAP1, respectively; expression level of DENND2D is 0.002 and 0.14 (expression in RPKM). **O.** UpSet plot of how MRRs overlap with each other in different cells. HOMER mergePeak command is used to overlap MRRs in different cells. **P.** Number of genes with different expression specificities that are associated with MRR. Gene expression in each cell line are compared with 69 facets of tissue/cell expression data from FANTOM. Specificity of each gene X is calculated by: Specificity(X)=1−(entropy(X)/log2(N)). Quartile Q1 and Q3 are used as cut-off of ‘Low specificity’ and ‘High specificity’, respectively. **Q & R.** Density of Hi-C anchors overlapped with constituent peaks of typical H3K27me3 peaks and constituent peaks of MRRs in GM12878 and HAP1 cells. The shuffled peaks were generated by expanding the midpoint of each constituent peaks to the median length of all the constituent peaks, and then followed by random genomic region shuffling. Wilcoxon test p values are as indicated.

**Figure S2**. Analysis of chromatin interactions associated with MRRs.

**A, B & C.** Number of MRRs that involved in chromatin interactions and are associated with genes in K562, GM12878, and HAP1 cells. Venn diagrams showed number of MRR associated with genes through chromatin interactions in different scenarios described in **Figure 2C**. **D, E & F.** Number of genes that are associated with MRRs in different positional relationships. These numbers were not necessarily the same as the numbers in Figure S1A-C, because the numbers were genes and MRRs, respectively. **G.** H3K27me3-rich regions (MRR) and typical H3K27me3 peaks in GM12878 HAP1, and their associated genes through chromatin interactions. Proximal, gene and peak occupy the same Hi-C anchor, or gene with promoter overlapping peak; Distal, the peak is connected to the gene via Hi-C interactions (they occupy different anchors of a Hi-C interaction). **H.** Expression of genes that are associated with MRR in proximal, distal, and internal category in GM12878 and HAP1 cells. The three categories are described in **Figure 2C**. The control category is generated by: 1) first filter out genes that are overlapped with ENCODE blacklist regions and also H3K9me3 peaks; 2) only retain genes that are overlapped with Hi-C interactions; 3) randomly sample the same amount of genes as the average gene number in proximal/distal/internal category. Wilcoxon test p values are indicated, ns: p > 0.05, *: p <= 0.05, **: p <= 0.01, ***: p <= 0.001, ****: p <= 0.0001. **I, J, K & L.** Examples of genes that were associated with MRRs in different categories as described in **Figure 2C**. **M.** Gene ontology analysis of MRR and typical H3K27me3 associated genes through chromatin interactions. **N, O & P**. Examples of 4C interactions of different candidate regions of different states in K562 cells. The colors of 4C interactions are based on the distal interacting regions to the 4C bait. Blue: repressive; orange: active; green: both; grey: quiescent. The state of the 4C bait is labeled by text. The numbers of different states of 4C interactions were given in table. **Q & R.** Validation of Hi-C interactions using *PSMD5* and *TOR1A* as 4C bait, respectively. Extensive ‘B-B’ interactions (green arcs) connecting *PSMD5* and *TOR1A* gene are shown in Hi-C and validated by 4C in K562 cells. The colors of Hi-C and 4C interactions are based on the distal interacting regions to the gene’s promoter and the 4C bait, respectively. Blue: repressive; orange: active; green: both; grey: quiescent. The numbers of different states of 4C interactions were given in table.

**Figure S3**. CRISPR candidate selection process and MRR1-A1 knock out leads to *FGF18* upregulation.

**A**. Selection process of H3K27me3-rich regions (MRRs) candidates to perform CRISPR. **B**. ChIP-qPCR of the MRR1 in wild-type K562 cells using H3K27me3 antibody, H3K27ac antibody and IgG antibody. Two regions (MRR1-R1 and MRR1-R2) within MRR1 were tested. The y axis indicates as percentage of total input. The data shows average + standard error. **C**. Genotyping results of three MRR1-A1 knockout clones (KO-1, KO-2 and KO-3). Genotyping is performed using flanking primers and internal primers along the deletion region and results are shown by agarose gel electrophoresis and Sanger sequencing. **D & E.** Table of MRR1-A1 looping genes and proximal genes. The fold changes in RNA-seq of KO clones of the looping and proximal genes are shown.

**Figure S4.** MRR1-A1 KO leads to erythroid differentiation.

**A**. Hemoglobin genes (*HBB*, *HBZ* and *HBE1*) expression decreased upon siRNA knocking down *FGF18* gene. **B.** MRR calling at *FGF18* region in seven cell lines (K562, GM12878, HAP1, H1hESC, HeLaS3, HepG2 and KARPAS). MRRs are indicated by blue bars and the *FGF18* gene region is highlighted by the red box. All data shown here are average + standard error. P value less than 0.05 is shown as *. P value less than 0.01 is shown as **. P value less than 0.001 is shown as ***.

**Figure S5**. MRR2-A1 knock out leads to upregulation of multiple genes including *IGF2* gene.

**A**. ChIP-qPCR of the MRR2 in wild-type K562 cells using H3K27me3 antibody, H3K27ac antibody and IgG antibody. Two regions (MRR2-R1 and MRR2-R2) within MRR2 were tested. The y axis indicates as percentage of total input. The data shows average + standard error. **B**. Genotyping results of three MRR2-A1 knockout clones (KO-1, KO-2 and KO-3). Genotyping is performed using flanking primers and internal primers along the deletion region and results are shown by gel electrophoresis and Sanger sequencing. **C & D.** Table of MRR2-A1 looping genes and proximal genes, and the fold changes in the RNA-seq of KO clones. **E.** RT-qPCR of *H19* gene shown in empty vector cells (EV) and three MRR2-C1 knock out cells (KO-C1, KO-C2 and KO-C3). **F.** Screenshot of MRR2 nearby regions with ChIP-seq of H3K27ac and H3K27me3 in EV and KO clones. MRR2 was indicated by yellow bar. **G.** Hemoglobin genes (*HBB*, *HBZ* and *HBE1*) expression decreased upon siRNA knockdown of *IGF2* gene. **H.** Model of MRR2-A1 knockout which leads to gene upregulation of multiple looping and proximal genes. Collectively, these changing gene expression levels lead to cell differentiation and tumor growth inhibition. **I.** MRR calling at *IGF2* region in seven cell lines (K562, GM12878, HAP1, H1hESC, HeLaS3, HepG2 and KARPAS). MRRs are indicated by as blue bars and *IGF2* gene region is highlighted by the red box. All data shown here are average + standard error. P value less than 0.05 is shown as *. P value less than 0.01 is shown as **. P value less than 0.001 is shown as ***.

**Figure S6**. Chromatin interactions changes upon MRR2-A1 KO were confirmed by 3C-PCR.

**A.** Chromatin interactions changed at *IGF2* bait compared to *IRF2BP2* (control) bait after silencer KO. The Y-axis shows the percentage of changing loops in KO clones**. B.** The losses of two different loops (F1 and F2) were validated by 3C-PCR followed by Sanger sequencing. Two different 3C libraries were used here. **C**. ChIP-qPCR at *IGF2* CRISPR excised region showed this region was completely deleted in the total input. **D**. *IRF2BP2* gene (control used in 4C-seq) had similar expression in EV and KO. ChIP-qPCR of H3K27me3 and H3K27ac for the *IRF2BP2* gene did not change after KO. Data shown here are average + standard error.

**Figure S7**. Histone modifications, gene expression, and chromatin interactions changes upon EZH2 inhibition.

**A & B.** Western blot of different histone marks in 1μM GSK343-treated K562 cells and HAP1 EZH2 KO clones, respectively. Bottom panel is the quantification of the western blot. **C & D.** TPM changes of genes associated with different types of peaks in 1μM GSK343-treated K562 cells and HAP1 EZH2 KO clones, respectively. Genes included: 1) Genes overlapped with different peaks as it is normally considered; 2) Genes associated with different peaks through HiC interaction. Wilcoxon test p value, ns: p > 0.05, *: p <= 0.05, **: p <= 0.01, ***: p <= 0.001, ****: p <= 0.0001. **E.** RT-qPCR of selected genes associated with different peaks in HAP1 WT and EZH2 KO clones. Proximal: gene and peak occupy the same Hi-C anchor; distal: the peak is connected to the gene via Hi-C interactions; both: both anchors of gene-associated Hi-C interactions overlap with the peak. **F.** Gene ontology analysis of differentially expressed genes in 1μM GSK343-treated K562, 5μM GSK343-treated K562, and HAP1 EZH2 KO clones. Up and down stand for the direction of expression changes. **G, H & I.** Volcano plot of differentially expressed genes in 1μM GSK343-treated K562, 5μM GSK343-treated K562, and HAP1 EZH2 KO clones, respectively. In each subfigure, top panels are: Top 10 most significant DE genes. Significance threshold of q value < 0.05 is used. Beta value is an estimator of fold change calculated by sleuth in natural log; bottom panels are: Cell-adhesion-related genes were labeled. These genes are under REACTOME pathways of “Extracellular matrix organization”, “Non-integrin membrane-ECM interactions”, “Non-integrin membrane-ECM interactions”, “L1CAM interactions”, and “ECM proteoglycans”. **J.** Cell morphology of HAP1 WT and EZH2 KO clones under microscope. **K.** Growth curve of HAP1 WT and EZH2 KO clones. **L & M.** 4C results using the bait of *FGF18*, *FGF18* KO region (MRR1-A1), *IGF2*, and *IGF2* KO region (MRR2-A1) in DMSO and 5μM GSK343-treated cells. The baits of 4C baits were indicated by vertical lines. The baits of each 4C panel were indicated by textboxes. For the baits at the gene promoters, the 4C baits were designed at around gene promoter. For the bait at zknockouts (KOs), the 4C baits were designed at the CRISPR KO regions (in Figure 3 & Figure 5). The colors of 4C interactions are based on the distal interacting regions to the 4C bait. Blue: repressive; orange: active; green: both; grey: quiescent. Height of 4C is in RPM. ChIP-seq signal and ChIP-seq peaks of H3K27ac, H3K27me3, and H3K4me3 in DMSO and 5μM GSK343-treated K562 cells were given in the lower panels. **N.** Comparison of 4C results in 5μM GSK343 and 1μM GSK343 treatment using baits of *FGF18*, *FGF18* KO region (MRR1-A1), *IGF2*, and *IGF2* KO region (MRR2-A1). ChIP-seq signal and ChIP-seq peaks of H3K27ac, H3K27me3, and H3K4me3 in WT K562 cells were given in the lower panels. Details of the representation of 4C and ChIP-seq are the same as in **Figure S7L-M**. **O.** 4C results of *IGF2*, *FGF18* and *HOXD13* in K562 DMSO and 1μM GSK343 treated cells. ChIP-seq signal and ChIP-seq peaks of K562 cells were given in the lower panels. ChIP-seq signal and ChIP-seq peaks of H3K27ac, H3K27me3, and H3K4me3 in WT K562 cells were given in the lower panels. Details of the representation of 4C and ChIP-seq are the same as in **Figure S7L-M**. **P.** 4C results using the bait of *IGF2*, *FGF18*, and *HOXD13* in WT and EZH2 KO HAP1 cells. ChIP-seq signal and ChIP-seq peaks of WT and EZH2 KO HAP1 cells were given in the lower panels. Details of the representation of 4C and ChIP-seq are the same as in **Figure S7L-M**. **Q.** 4C results of *MYC*, *PSMD5*, *LINC00910* and *ZDHHC11* in HAP1 WT and EZH2 KO clones. ChIP-seq signal and ChIP-seq peaks of WT and EZH2 KO clones were given in the lower panels. Details of the representation of 4C and ChIP-seq are the same as in **Figure S7L-M**. **R.** 4C results of *MYC*, *PSMD5*, *LINC00910* and *ZDHHC11* genes in DMSO and 1μM GSK343 treated K562cells. ChIP-seq signal and ChIP-seq peaks of H3K27ac, H3K27me3, and H3K4me3 in WT K562 cells were given in the lower panels. Details of the representation of 4C and ChIP-seq are the same as in **Figure S7L-M**.

**Figure S8**. Chromatin interactions changes upon GSK343 treatment were confirmed by 3C-PCR.

**A.** Proportion of unchanged 4C interactions in different distance categories in 1uM GSK343-treated K562 cells and HAP1 EZH2 KO clones. As the distance of 4C interactions increases, the proportion of unchanged 4C interactions drops, indicating that long-range interactions are perturbed. **B.** Table of RPMs of 4C chromatin interactions in two different replicates in DMSO and GSK343 condition. **C.** 3C-PCR gel image and Sanger sequencing of the *IGF2*-MRR2-A1 loop in DMSO and GSK343-treated cells (two replicates). **D.** 3C-PCR gel image and sanger sequencing of *TRPM5*-MRR2-A1 loop in DMSO and GSK343-treated cells (two replicates).

**Figure S9**. Integrative analysis of H3K27me3, H3K27ac and chromatin interactions upon EZH2 inhibition.

**A.** 4C interaction intensity changes at 4C regions with different levels of H3K27me3/H3K27ac signal in DMSO-treated K562 cells. Left panel, different 4C regions are classified according to their H3K27me3 signal intensity in DMSO-treated K562 cells. H3K27me3 signal level at these 4C regions are tertiled in three cohorts: high, medium, and low. 4C regions type indicates different categories of 4C regions. Gained, 4C interactions present in GSK343-treated 4C but not DMSO-treated 4C; lost, 4C interaction present in GSK343-treated 4C but not DMSO-treated 4C; unchanged, 4C interactions present in both GSK343-treated and DMSO-treated 4C. The 4C interaction intensities are shown in log_10_ transformed RPM. Right panel, different 4C regions are classified according to their H3K27ac signal intensity in DMSO-treated K562 cells. Similar to the left panel, H3K27ac signal level at these 4C regions are tertiled in three cohorts. **B.** Screenshot of H3K27me3 and H3K27ac ChIP-seq in DMSO and GSK343 condition as well as zoomed-in view at *FGF18* and MRR1-A1 site. **C.** 4C chromatin interactions using *IGF2* promoter as the bait in DMSO and GSK343. **D.** 4C chromatin interactions using *FGF18* promoter as the bait in DMSO and GSK343. The red error indicates the differences between *IGF2* and *FGF18*.

**Supplementary Tables**

**Table S1**. Putative human silencer examples.

**Table S2**. Comparison between different identification methods of human silencers.

**Table S3.** Libraries Used (Excel Spreadsheet). This is a list of all the libraries used in this manuscript. More details can be found in the “README” tab.

**Table S4.** Primers used for ChIP-qPCR, RT-qPCR, Sanger Sequencing, 4C-Seq and guide RNAs used for CRISPR.

**Table S5.** Genomic coordinates of 4C viewpoints.

**Table S6.** MRRs, typical H3K27me3 peaks, super-enhancers and typical H3K27ac peaks in K562 GM12878 and HAP1 (Excel Spreadsheet). This table lists all the coordinates and associated genes of the different categories of peaks. More details can be found in the “README” tab.

**Table S7.** MRRs in other cell lines (Excel Spreadsheet). This table lists all the MRRs in other cell lines (H1hESC, HEPG2, HeLaS3, KARPAS-422, Pfeiffer, WSU-DLCL2) in addition to K562, GM12878 and HAP1. More details can be found in the “README” tab.

**Table S8.** Transcription factor binding enrichment at interacting regions of MRRs (Excel Spreadsheet). This table lists the binding enrichment of different TFs (CTCF, EZH2, GATAD2B, RAD21, REST, SMC3, YY1, and ZNF143) at the interacting regions of MRRs. More details can be found in the “README” tab.

**Table S9.** Differentially expressed genes in RNA-Seq of HAP1 and K562 cells after EZH2 knockout or inhibition (Excel Spreadsheet). This table lists all the differentially expressed genes from the RNA-Seq that we performed for this paper (related to Figures 3). More details can be found in the “README” tab.

**Table S10.** ChIP-seq peaks in HAP1 and K562 cells after EZH2 knockout or inhibition (Excel Spreadsheet). This table lists all the ChIP-Seq peaks of: 1) 5μm GSK343-treated K562 cells and 2) HAP1 EZH2 knockout cells that we performed for this paper (related to Figure 3). More details can be found in the “README” tab.

**Table S1**. Putative human silencer examples.

| Element and locus | Gene |
| --- | --- |
| An AT-rich octamer –binding domain located -140bp from TSS | Human thyrotropin-β gene (*hTSHβT*)^24^ |
| This silencer (5’ CTCTCTAGAGAG 3’) is located -1800 bp from the transcription initiation site | plasminogen-activator-inhibitor type-2 gene (*PAI-2*)^25^ |
| PRE2-S5 | *CCND1* gene^26^ |
| 5’ untranslated region of the first exon | human α1-chimaerin gene^27^ |
| 8bp silencer elements (-97 to -90bp) in the 5’ promoter | human Pi Class  glutathione S-transferase (*GSTP1*) gene^28^ |
| 5’-GGCATAATGGGTCTGTCTCATCGTC-3’ (-211bp to -186bp) | Human interferon-γ gene^29^ |
| MECP2-F3 (in the 210kb region encompassing *MECP2* gene) | *MECP2* gene^30^ |
| 5’- TTCAGCACCNCGGACAGNGCC-3’ (-235bp to -199 bp) | Human synapsin I^31^ |
| Not shown | human INK4b-ARF-INK4a locus^32^ |
| A GU-rich region located  downstream of splicing site 2 | human papillomavirus late mRNA^33^ |
| 5’-GGGGAGGGGG-3’ (-1418bp to -1388bp) | platelet-derived growth factor A-chain gene^34^ |
| A 24bp element within the third intron of the human collagen type IV gene | human collagen type IV gene^35^ |
| First intron of the human apolipoprotein A-II gene | human apolipoprotein A-II gene^36^ |
| A 190bp fragment from the first intron of the human *CD4* gene | *CD4* gene^37^ |
| A 1.8-Mb region of human chromosome 7^38^ | Not shown |
| T39, located on the short arm of the X chromosome (p22.2) in the first intron of the gene *ARHGAP6*^39^ | Not shown |
| Not shown | *BDNF* gene in Huntington’s disease^40^ |

**Table S2**. Comparison between different identification methods of human silencers.

|  | | Huang *et al*^41^ | Jayavelu *et al*^42^ | Pang and Snyder^43^ | Ours |
| --- | --- | --- | --- | --- | --- |
| Methods | | Authors focused on the DHSs overlapping within H3K27me3 peaks (H3K27me3-DHSs), and then explore their correlation with expression of nearby genes across tissues. In this way, H3K27me3-DHS were categorized into three groups: positively correlated (posCOR), negatively correlated (negCOR) and uncorrelated (unCOR). NegCORs were identified as silencers | Authors considered DHS from 82 human cell lines (ENCODE+Roadmap) or from 22 mouse cell line (ENCODE) and the exclude all that overlapped with chromatin features of enhancers, promoters or insulators. The remaining DHS were filtered for cell-type specificity and considered “putative silencer elements” | Authors used the high-throughput ReSE lentiviral screen system. If silencers (fixed size=200bp) are inserted, cells wont undergo apoptosis | We first identified H3K27me3 peaks through H3K27me3 ChIP-seq data in 7 cell lines, then clustered nearby peaks, and ranked the clustered peaks by average H3K27me3 signals levels. The top clusters with the highest H3K27me3 signal were called H3K27me3-rich regions (MRRs) |
|  | Numbers of silencers | 1334 per cell line | Each cell type contains tenths of thousand elements | 2664 potential silencers in k562 cells | Unique to different cell line, eg:974 MRR in K562 |
|  | Specificity | Tissue specific | Cell type specific.  Silencers can act as enhancers in other cell types | Tissue specific | Cell type specific |
|  | TF binding enrichment | Common Repressors: MECOM, SMAD family, FOXG1, CTCF, TCF and SOX21.  Tissue specific TFs: STAT family, EHF, TFAP2A, NKX6-1 and E2F4 | Enriched for binding of REST, YY1, ZBTB33, SUZ12 and EZH2 (based on ChIP-seq) and also for motifs of other repressor TFs | AP2 binding domains were found in both K562 and HepG2. CDH4 and NCoR enriched in K562 while EZH2 and REST enriched in K562 | Different silencer clusters have different TFs |
|  | Sequence signatures | Not explored | Many silencers are hypermethylated and poor conservation | Not explored | No difference in terms of CpG islands |
|  | Chromatin interactions | Used Hi-C to link distal genes but only focused on proximal genes | Majority promoter interactions of silencers correspond to non-expressed or lowly expressed genes | ReSE silencers can directly interact with many gene promoters | MRRs are highly associated with chromatin interactions than typical ones and based on the Hi-C, can be classified into distal, internal looping and proximal silencers |
|  | Disease traits (SNPs) | 6.7% of silencers harbor at least one GWAS SNP | 25% disease traits present are enriched at silencers, similar to other cis-regulatory elements | Not explored | Not explored |
|  | Validation | Five out of ten predicted silencers have decreased luciferase reporter gene activity.  Sharpr-MPRA results showed predicted silencers showed repressive impact | 51% of 7500 selected silencers in K562 was validated through MPRA | Three silencers were validated using CRISPR | Validate two candidate (*FGF18* and *IGF2*) from internal looping category in K562 by CRISPR excision and RT-qPCR |
|  | Mechanism | Not explored | Not explored | Not explored | Based on the IGF2 silencer example, chromatin interactions and histone state control the *IGF2* expression.MRR removal cause chromatin interactions landscape change. Specifically, more active loops gained and repressed CI lost which finally leads to H3K27me3 loss at *IGF2* region and *IGF2* expression upregulation |

**Table S4.** Primers used for ChIP-qPCR, RT-qPCR, Sanger Sequencing, 4C-Seq, 3C-PCR and guide RNAs used for CRISPR.

| Primer name | Sequence (5’ to 3’) |
| --- | --- |
| ChIP-qPCR primers | |
| MRR1-R1-F | TGGTCCATAATGGCTTATCATCCT |
| MRR1-R1-R | GAAACACTTGGCTGTCTGGC |
| MRR1-R2-F | AGCCTGCAGAAATTGACATTGC |
| MRR1-R2-R | CTCTGGCTCCTTTTGGCTCA |
| MRR2-R1-F | AAGAGGGGACCCATCAGGG |
| MRR2-R1-R | TGGGTTTTCTGGAGCCAAGG |
| MRR2-R2-F | TTGTTAGGGACGTCAGTGGC |
| MRR2-R2-R | TGGAGGGAAACGGCAAAAGA |
| HAP1-SE-WT-2ND-F | AGGACAGTTGGGAAGGAGGA |
| HAP1-SE-WT-2ND-R | CAGGGCTGCTGCACCTAC |
| HAP1-SE-WT-3RD-F | CGAGTCCTTCCCGGTGAGAT |
| HAP1-SE-WT-3RD-R | AAGTCCACGGAACAACGACA |
| HAP1-SS-WT-2ND-F | GTATAACACGCCTGCACAGACA |
| HAP1-SS-WT-2ND-R | TCCCAATCCACGTGAGGGA |
| HAP1-SS-WT-4TH-F | CAGAACTTTGCAGCAGGCAG |
| HAP1-SS-WT-4TH-R | GGAGCAGCAGCATCCACTTA |
| IGF2-R1-F | CACCGCGTCAACATACCAGG |
| IGF2-R1-R | CATGTGTGATTCGTGCCTTGC |
| IGF2-R2-F | AAGCAAGGAAGTCACGGGTC |
| IGF2-R2-R | GAGAAATAGGGCTTCGGGCG |
| IGF2-R3-F | GGGCATCTCTGTCATGGTGG |
| IGF2-R3-R | GGCATTTGGGATACACCCGT |
| IGF2-R4-F | GCGAGGTAAACCTCCCAGAG |
| IGF2-R4-R | CGGGTCTGGTGATGCCATAG |
| IGF2-KOsite-F | GCACTCCAAGAAAAGGCCAG |
| IGF2-KOsite-R | GACGTCCCTAACAAAGTGCC |
| IRF2BP2-ChIP-F | GGACAGTGAACAGCGGTCAA |
| IRF2BP2-ChIP-R | CTGAGAGTGCTGCTGGGAAA |
| RT-qPCR primers | |
| GAPDH-F | GCACCGTCAAGGCTGAGAAC |
| GAPDH-R | GGATCTCGCTCCTGGAAGATG |
| FGF18-RT-F | GACGATGTGAGCCGTAAGCA |
| FGF18-RT-R | GAGCTGGGCATACTTGTCCC |
| UBTD2-RT-F | CACCGGAGTTGCTCTAGGTC |
| UBTD2-RT-R | TCCCTCTTGCTGCGTAGTTG |
| FBXW11-RT-F | CCAGTGTCAGATGTCTCCAGATAA |
| FBXW11-RT-R | AGTTTCCTTCTGATGGCCTCTT |
| IGF2-RT-F | TCCTGTGAAAGAGACTTCCAGC |
| IGF2-RT-R | TTGGTGTAGCTCAGCAGAAGG |
| H19-RT-F | CAGGAGTGATGACGGGTGGAG |
| H19-RT-R | CCCTTCTTTCCAGCCCTAGCTC |
| KCNA2-RT-F | ATCCGGTTGGAACGCAGAC |
| KCNA2-RT-R | AGGTGCAGTCATGTGAGGTG |
| BARX2-RT-F | TGCATTCCTGTACGGGCTC |
| BARX2-RT-R | CAGGTGGGAGATGACAGTGG |
| ADAMTS8-RT-F | AGACTGTCTCCTGGATGCCC |
| ADAMTS8-RT-R | AAAGATCTGCCTGCACTGCT |
| SFRP5-RT-F | CTGAAGGGCACTCCTCCTTG |
| SFRP5-RT-R | CCCATCCCTTAGGCCTTGTG |
| HOXB7-RT-F | TCGAGCCGAGTTCCTTCAAC |
| HOXB7-RT-R | TCAGTTCCTGAGCTTCGCAT |
| HS3ST3A1-RT-F | GTCCAAGAGCAGTTTGGAGC |
| HS3ST3A1-RT-R | GCCCGAGTTCAGGTTCTCTC |
| HAS2-RT-F | GGCCGGTCGTCTCAAATTCA |
| HAS2-RT-R | TCACAATGCATCTTGTTCAGCTC |
| NTS-RT-F | GAACAGCCCAGCTGAGGAAA |
| NTS-RT-R | CCTGGATTAACTCCCAGTGTTGA |
| MEIS1-RT-F | ACGGCATCTACTCGTTCAGG |
| MEIS1-RT-R | CCATCACCTTGCTCACTGCT |
| HOXB9-RT-F | TGGGACGCTTAGCAGCTATT |
| HOXB9-RT-R | CGTACTGGCCAGAAGGAAAC |
| EPHB6-RT-F | CGGCCAACGGGAAGAAATAAA |
| EPHB6-RT-R | TGTCACAATGAAGACAAACAGGC |
| GABRQ-RT-F | CCAGGGTGACAATTGGCTTAA |
| GABRQ-RT-R | CCCGCAGATGTGAGTCGAT |
| CDK14-RT-F | CCAAGGAGTTGCTGCTTTTC |
| CDK14-RT-R | GAATGAACTCCAGGCCATGT |
| TMEM108-RT-F | TTTCTCCTGAGCCGTCGGA |
| TMEM108-RT-R | GATTCTGTCCTGGAGTAGAGGG |
| CD276-RT-F | CTCTGACAGCAAAGAAGATGATGG |
| CD276-RT-R | TCCTTTGGAGAAGGAGCCCA |
| SRSF4-RT-F | TCATTCAAGGTCTCGCTCTCG |
| SRSF4-RT-R | ACCTGGACCGAGATCTACTCT |
| RAI1-RT-F | GATGCCTCCACACCTACCAC |
| RAI1-RT-R | GCAGCCTCTTATGTTTGGGAC |
| SRGAP1-RT-F | CCAACATTGATGCCTGTCC |
| SRGAP1-RT-R | TCTCATAAACAGGGCCATCC |
| TRIM13-RT-F | CCTTTCCCACTAGCCGGAGTA |
| TRIM13-RT-R | TCCATCACATCCTCTGTCTCCT |
| LIMK1-RT-F | ACCTCAACTCCCACAACT |
| LIMK1-RT-R | TCTCGCACAGGACGATCC |
| PCBP1-RT-F | AAAGGCGGGTGTAAGATCAAAG |
| PCBP1-RT-R | GGCAAATCTGCTTGACACACTC |
| DUSP12-RT-F | ATCTATGGCGCCTCTTCGTG |
| DUSP12-RT-R | CTGCATGACAGTGCACCAAC |
| INTS7-RT-F | ATCCTGTGGCAAGAGCCATC |
| INTS7-RT-R | TGTGCAGAGAAGTTTGCAGC |
| VPS45-RT-F | GTACTTGTCAATTCGCCGCC |
| VPS45-RT-R | ACTCACTATGCCAGTCGTCTC |
| DLX3-RT-F | CTTACTCGCCCAAGTCGGAA |
| DLX3-RT-R | TCCTTCACCGACACTGGGT |
| ZNF639-RT-F | ACCCTTCTCGTTATTCAGATTCCT |
| ZNF639-RT-R | GGCTGTCTCATAGAACAGACACT |
| HOXD13-RT-F | GGCACGAGGCCTACATCTC |
| HOXD13-RT-R | TTAGAGCCACATCCCCTGGA |
| HBB-RT-F | GAGAACTTCAGGCTCCTGGG |
| HBB-RT-R | GCGAGCTTAGTGATACTTGTGG |
| HBZ-RT-F | CCGGTCAACTTCAAGCTCCT |
| HBZ-RT-R | CTCAGCGGTACTTCTCGGTC |
| HBE1-RT-F | TCACTAGCAAGCTCTCAGGC |
| HBE1-RT-R | AAACAACGAGGAGTCTGCCC |
| IRF2BP2-F | GGCCCTTCGAGAGCAAGTTT |
| IRF2BP2-R | CTTGCAACTGCTTTAGACCCG |
| CRISPR genotyping primers and guide RNAs | |
| CRISPR-step2-F | GCCTTTTGCTGGCCTTTTGCTC |
| CRISPR-step2-R | CGGGCCATTTACCGTAAGTTATGTAACG |
| MRR1-gRNA-F-F | CACCGTTTTTCCTGTCACGCTGCTC |
| MRR1- gRNA-F-R | AAACGAGCAGCGTGACAGGAAAAAC |
| MRR1- gRNA-S-F | CACCGCTCTGCTTTAAGAGCATCAC |
| MRR1- gRNA-S-R | AAACGTGATGCTCTTAAAGCAGAGC |
| MRR2- gRNA-F-F | CACCGCTTGGAAGAGGGGACCCATC |
| MRR2- gRNA-F-R | AAACGATGGGTCCCCTCTTCCAAGC |
| MRR2- gRNA-S-F | CACCGGGGCTCAAGGGCATGCTACG |
| MRR2- gRNA-S-R | AAACCGTAGCATGCCCTTGAGCCCC |
| MRR1-flanking-F | TGGGCTTTTTCTTAAGCTGCC |
| MRR1-flanking-R | TGGCCAAGTTGATTGTGTTAGT |
| MRR1-internal-F | CACCTCAACAAACCGATCACC |
| MRR1-internal-R | GAATCTAATAGCTGAGGACGAGC |
| MRR2-flanking-F | GATTCGGCCCCTACTTGGAT |
| MRR2-flanking-R | GTGGTTTGTAGAGGGGTGGC |
| MRR2-internal-F | CCCCTGACAGAGGGGCA |
| MRR2-internal-R | GGTGGTTTGTAGAGGGGTGG |
| 4C Primers | |
| IGF2-Outer-F | TCTCACGGAGCATCTGTCC |
| IGF2-Outer-R | TTACAGAGCTAGCACCTGGG |
| IGF2-Nest-F | GAAGCCCACCTTCCACTCA |
| IGF2-Nest-R | GGCTGGTCTCGACAACAAAG |
| FGF18-Outer-F | TCAGGCCTCCTTGCAAGCTAT |
| FGF18-Outer-R | GAACGAAGCTGCCTAGTATGC |
| FGF18-Nest-F | ACATCGGACACAAGTGCAGAA |
| FGF18-Nest-R | TCCAAAGCTGGCTCGCCTA |
| MRR1-Outer-F | CTGGCAGAACGGTTTAGACT |
| MRR1-Outer-R | CAGGGAGGTTACTGCACTTCAT |
| MRR1-Nest-F | CCTCCATACTTTCCCAGGGC |
| MRR1-Nest-R | AAATCCTTGTTCAATGCTTCCC |
| MRR2-Outer-F | GACCCTTGACTTGGGAGTGG |
| MRR2-Outer-R | CATGCCAGGCCTTTCAACTG |
| MRR2-Nest-F | GTTTGCATTCACACGCCCTC |
| MRR2-Nest-R | GCCCTGTGGGTTTGAGAAACT |
| HOXD13-Outer-F | TCATCGGCCATTTCCCTGAG |
| HOXD13-Outer-R | TATCTTACTGGCGACCGTGG |
| HOXD13-Nest-F | GTCAACTGCTCTGTGCAGACTG |
| HOXD13-Nest-R | TCTTACTGGCGACCGTGGAC |
| MYC-Outer-F | TGAAAGAATAACAAGGAGGTGGC |
| MYC-Outer-R | AGAAGGTCCGAAGAAAGAGGA |
| MYC-Nest-F | ATGGAGAACCGGTAATGGCA |
| MYC-Nest-R | AAGGAGGTGGCTGGAAACTT |
| PSMD5-Outer-F | AAAATGAGGAAGACTTGGCTTGC |
| PSMD5-Outer-R | GCAGTTACCACATGATTGCAACT |
| PSMD5-Nest-F | TTTGGCCTTGGACAGATAAT |
| PSMD5-Nest-R | GTTGTCACGATGTAACTTGAAC |
| TOR1A-Outer-F | TGAAATGAGTGAGCCCGGAAA |
| TOR1A-Outer-R | AACCCGATGACATCCAGGAAG |
| TOR1A-Nest-F | TGCGTGGAGCAGTTAATACC |
| TOR1A-Nest-R | GTTCAGACCACCCTCGTAAAT |
| LINC00910-Outer-F | GACCAAATGCACCAAGAGGG |
| LINC00910-Outer-R | ACCGTGACCCAAACTCTCAT |
| LINC00910-Nest-F | TCTGAAGGTACACAGTGACCA |
| LINC00910-Nest-R | TCCATTCATGTCACAGGTGGA |
| ZDHHC11-Outer-F | TGGTGTCACATTTAGAGGACCA |
| ZDHHC11-Outer-R | AAACAGGACCATGGTTCTTTGG |
| ZDHHC11-Nest-F | TCCCAAAGAGACAACAAGGACT |
| ZDHHC11-Nest-R | TCGCCAAGTCACATTGGTAAAA |
| CCND3-Outer-F | CACGTATTGTCTCCCCACTTT |
| CCND3-Outer-R | TGGTCGGTGTAGATGCACAG |
| CCND3-Nest-F | TATTGTCTCCCCACTTTCCAGG |
| CCND3-Nest-R | TGGGGACGCAAGACAGGTAG |
| TMCO4-Outer-F | CTACGCCTCAGTTTGCTGC |
| TMCO4-Outer-R | CCGTACCTACCATCACCCTG |
| TMCO4-Nest-F | GCAGTGGCTCACACCTGTAA |
| TMCO4-Nest-R | TGTAAAATGCCCAGCACAAT |
| 3C-PCR primers | |
| IGF2-promoter-3C-F | GTCACACTTGAGCACCTCCTGGTAACTG |
| F1-R | CATGGAGGTTGGCCTCGTTCTTCTTG |
| F2-R | CTAACCTGACCTGCTCCTTCGACATCTA |

**Table S5**. Genomic coordinates of 4C viewpoints.

| Viewpoint region | Genomic coordinates |
| --- | --- |
| *IGF2* promoter | chr11:2175368-2175873 |
| *FGF18* promoter | chr5:170845209-170845956 |
| *HOXD13* promoter | chr2:176961136-176961617 |
| MRR1-A1 | chr5:171338492-171339433 |
| MRR2-A1 | chr11:2042072-2043745 |
| *MYC* promoter | chr8:128,748,315-128,753,680 |
| *PSMD5* promoter | chr9:123,578,332-123,605,299 |
| *TOR1A* promoter | chr9:132,575,221-132,586,441 |
| *LINC00910* promoter | chr17:41,299,393-41,546,115 |
| *ZDHHC11* promoter | chr5:655,809-883,388 |
| *CCND3* promoter | chr6:41,902,671-41,909,552 |
| *TMCO4* promoter | chr1: 20127601-20133430 |

**Supplementary Figures**

**Figure S1A-D**

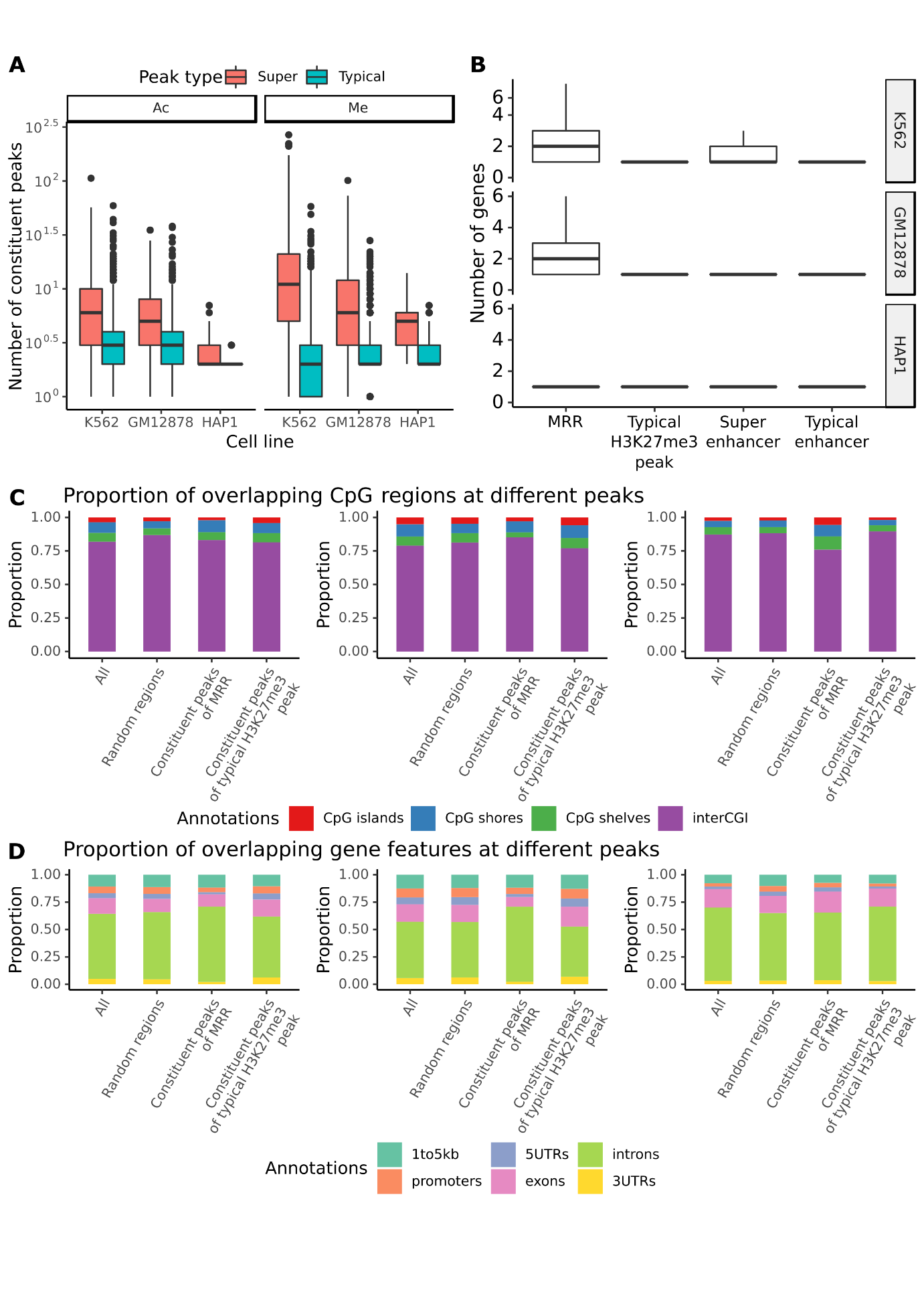

**Figure S1E**

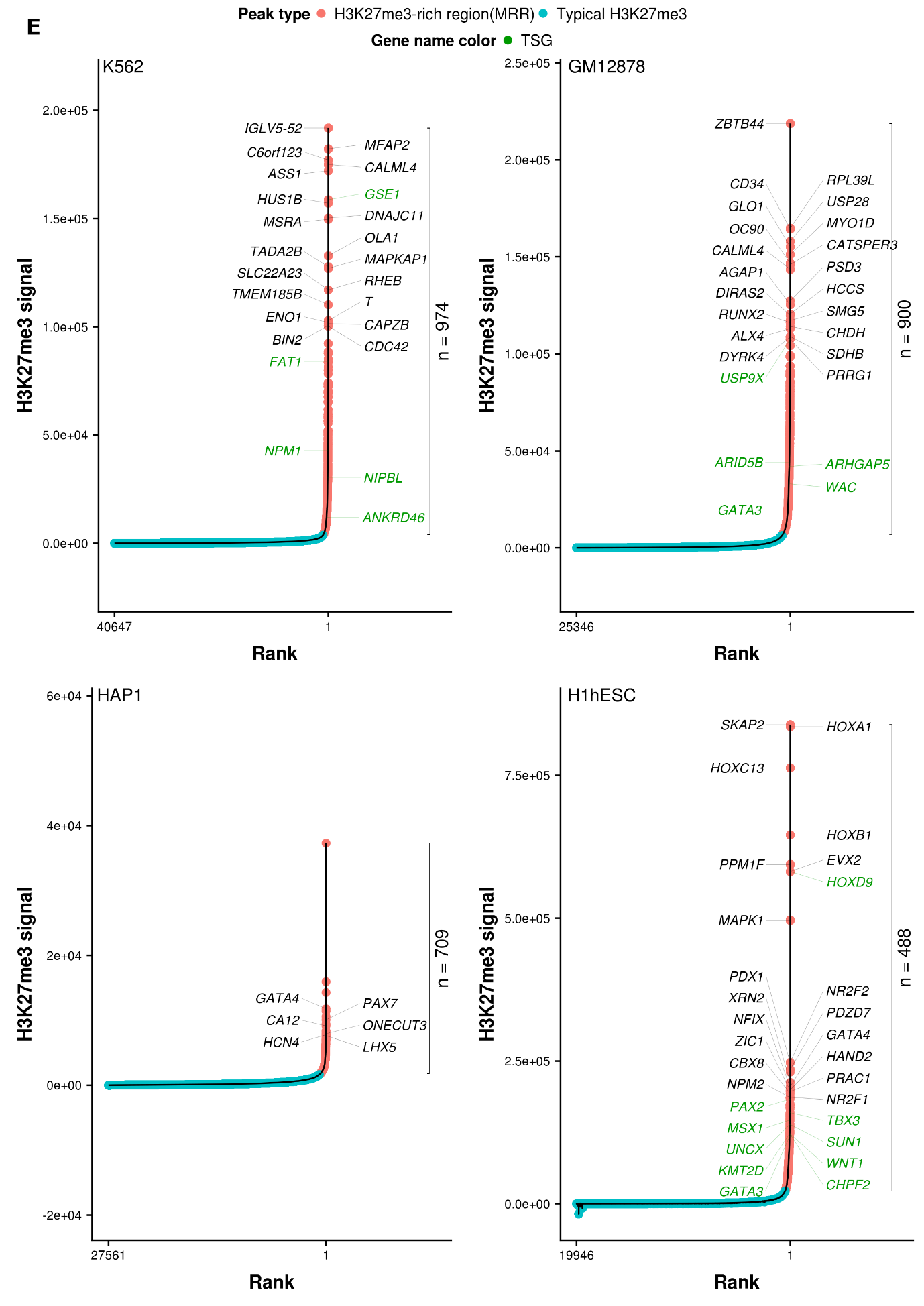

**Figure S1F-G**

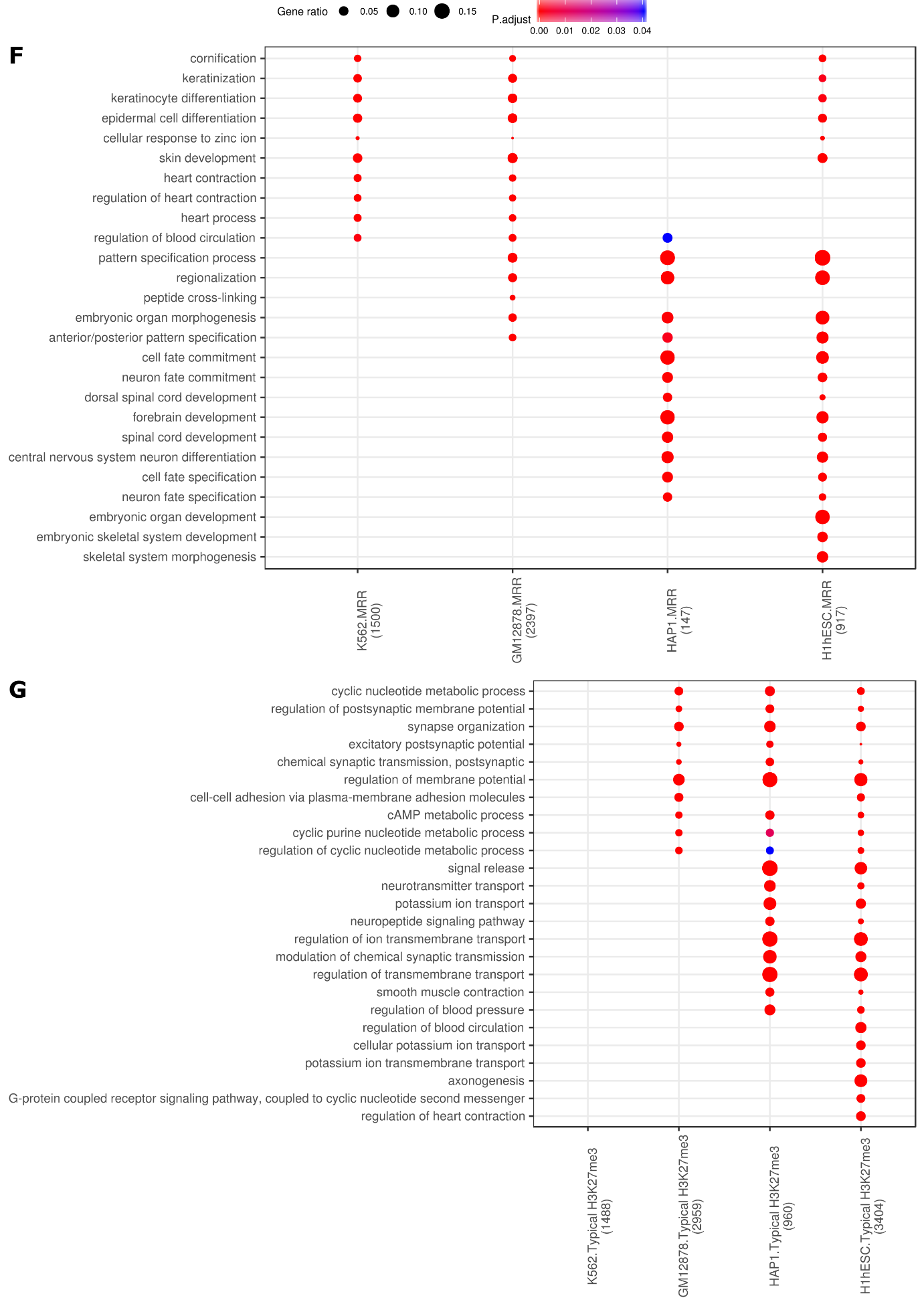

**Figure S1H-K**

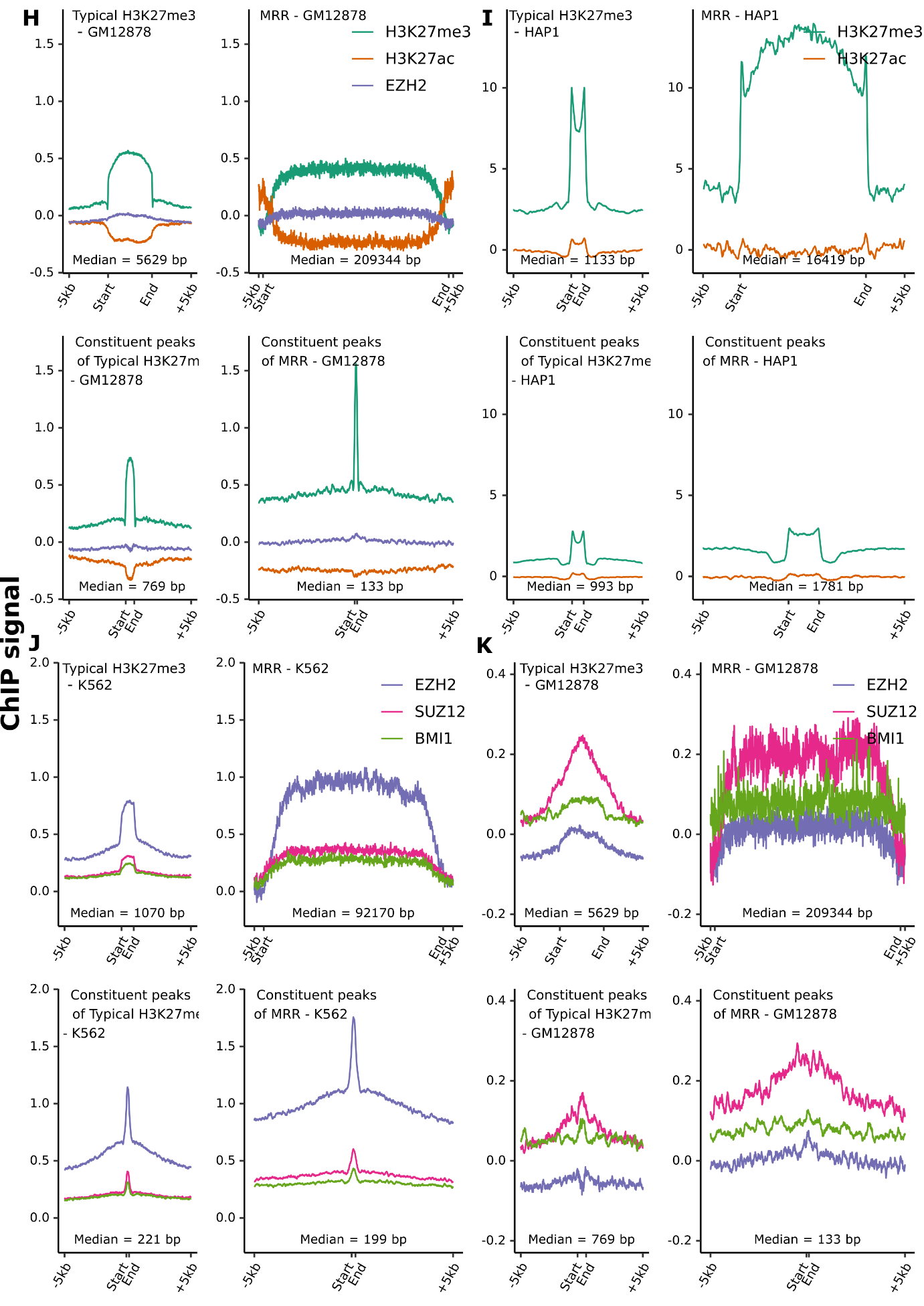

**Figure S1L**

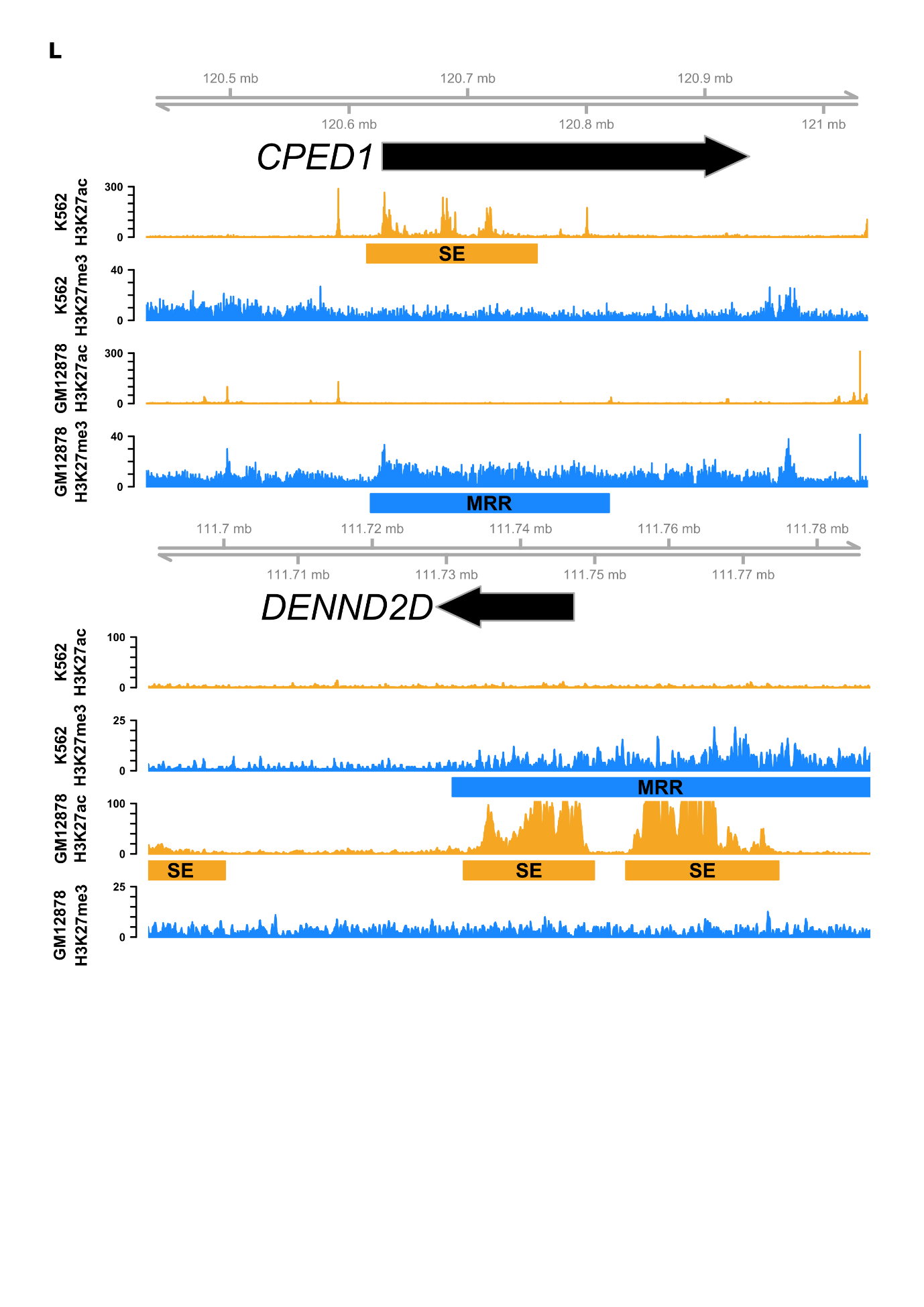

**Figure S1M-N**

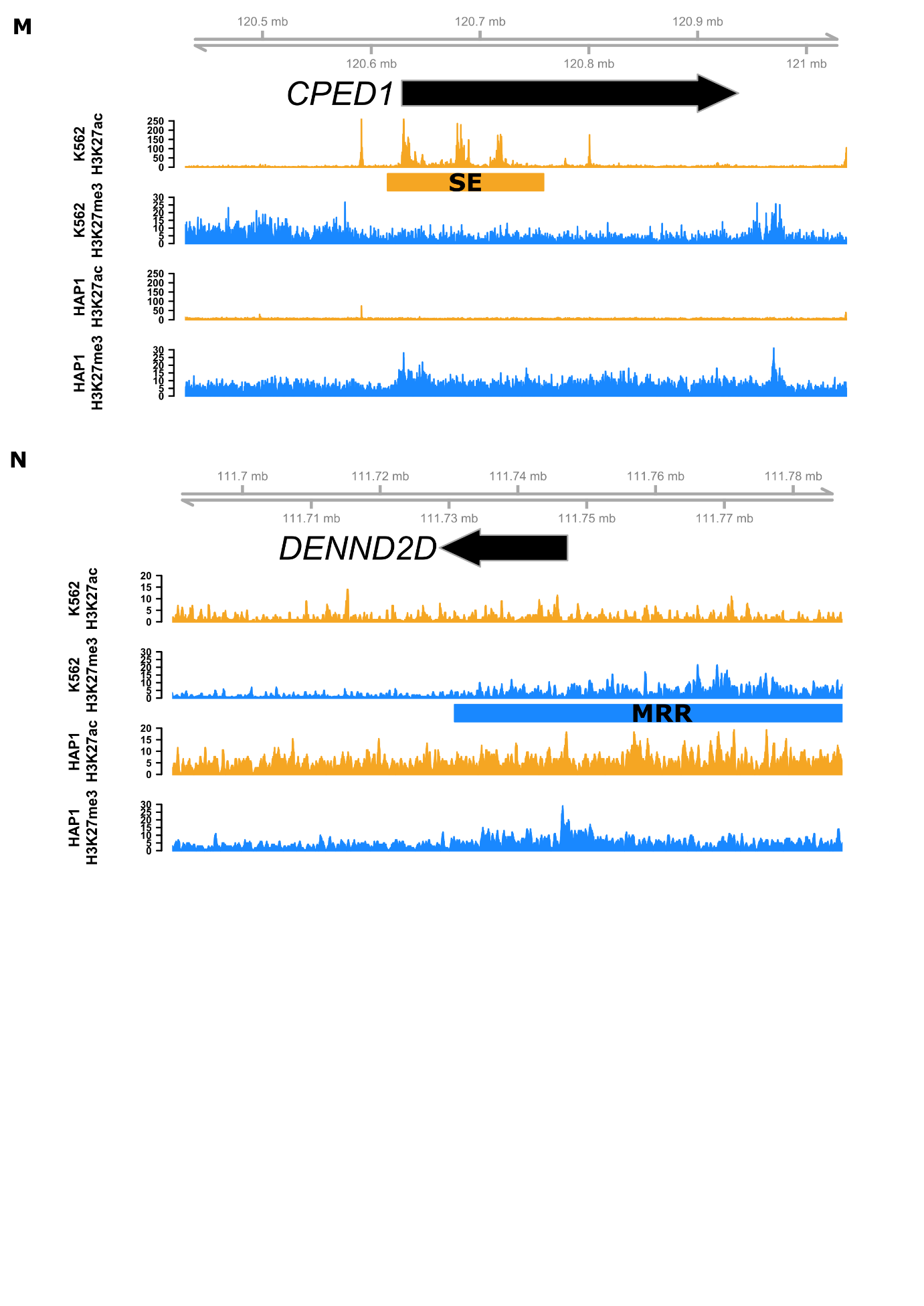

**Figure S1O**

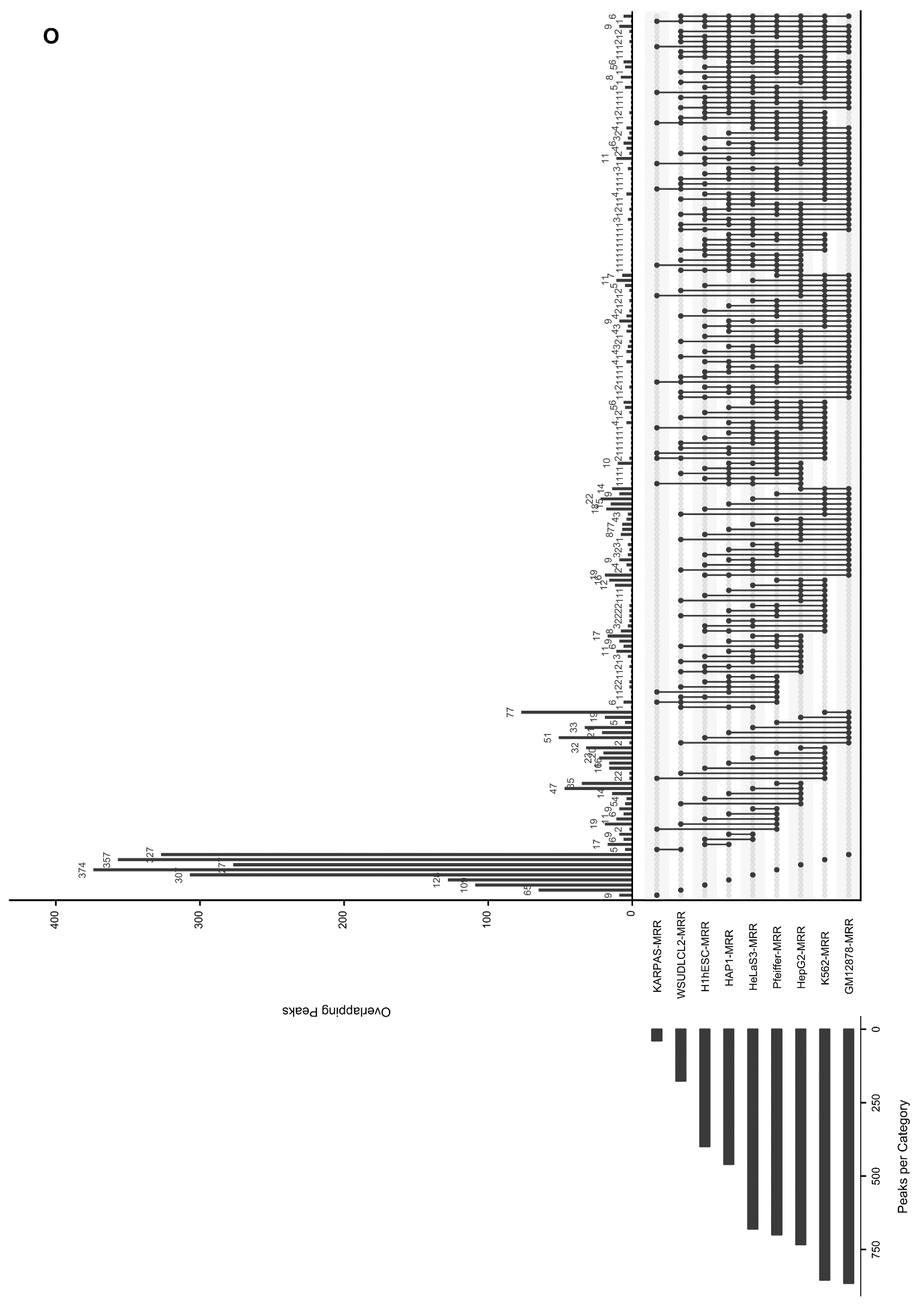

**Figure S1P-R**

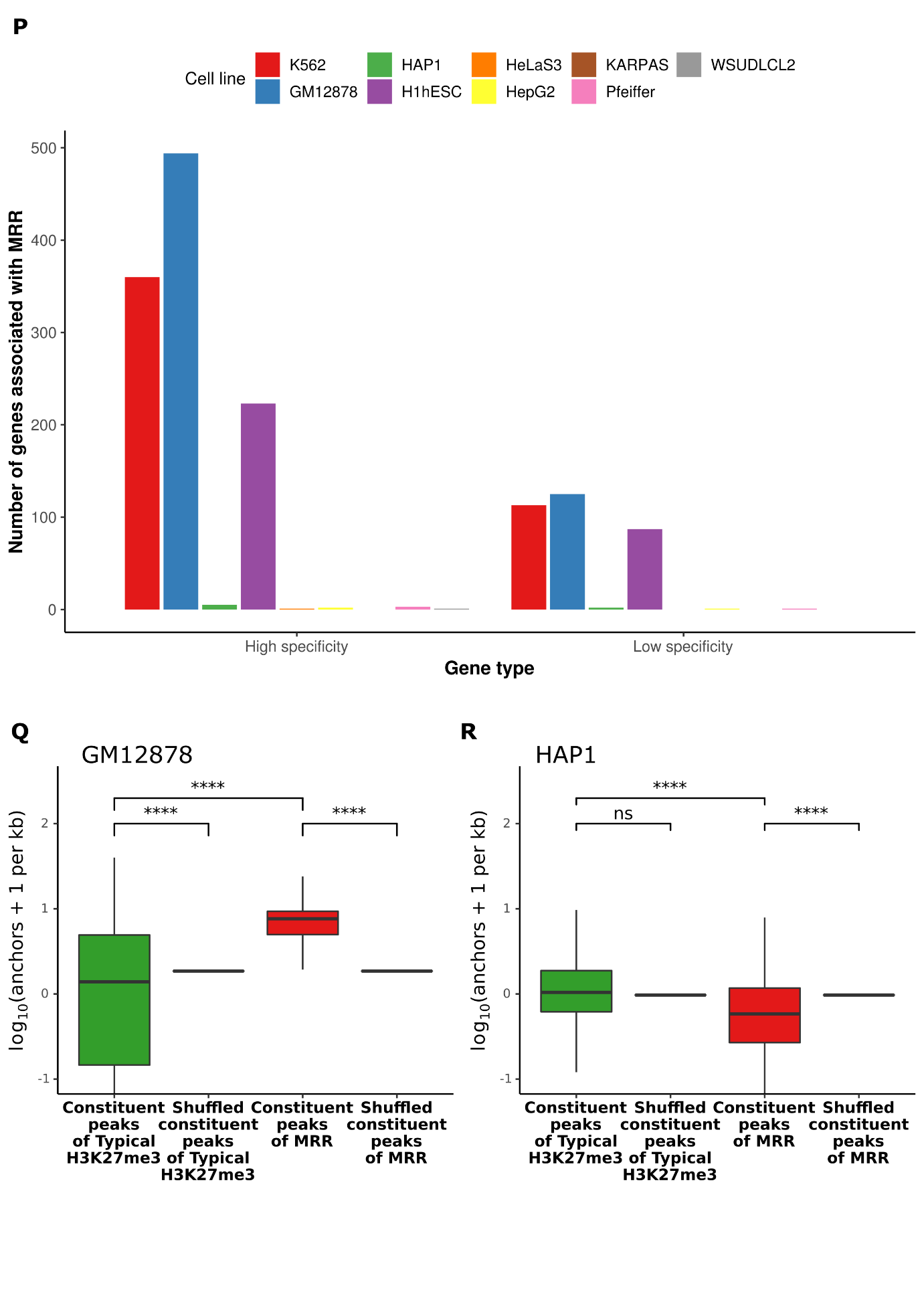

**Figure S2A-C**

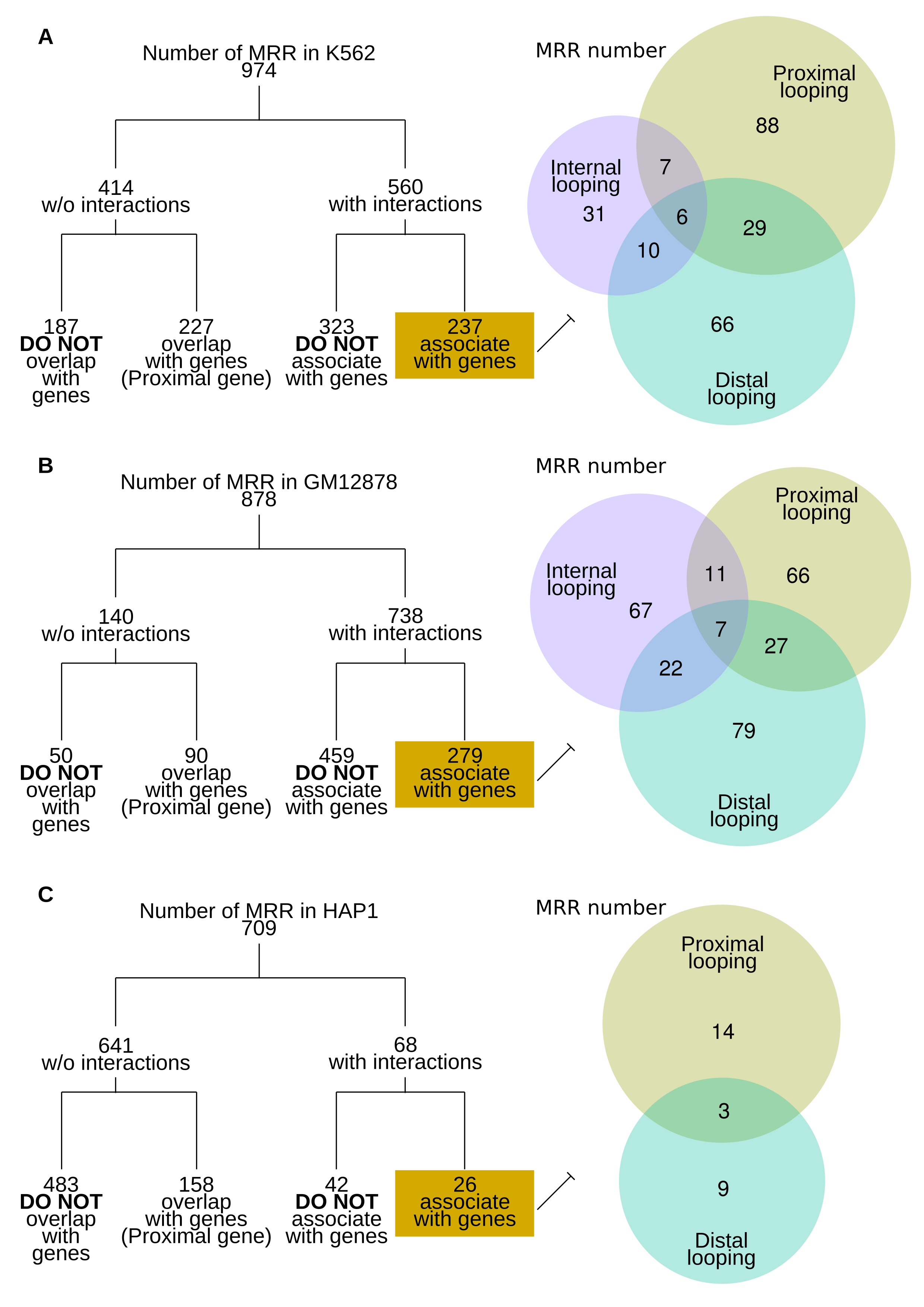

**Figure S2D-F**

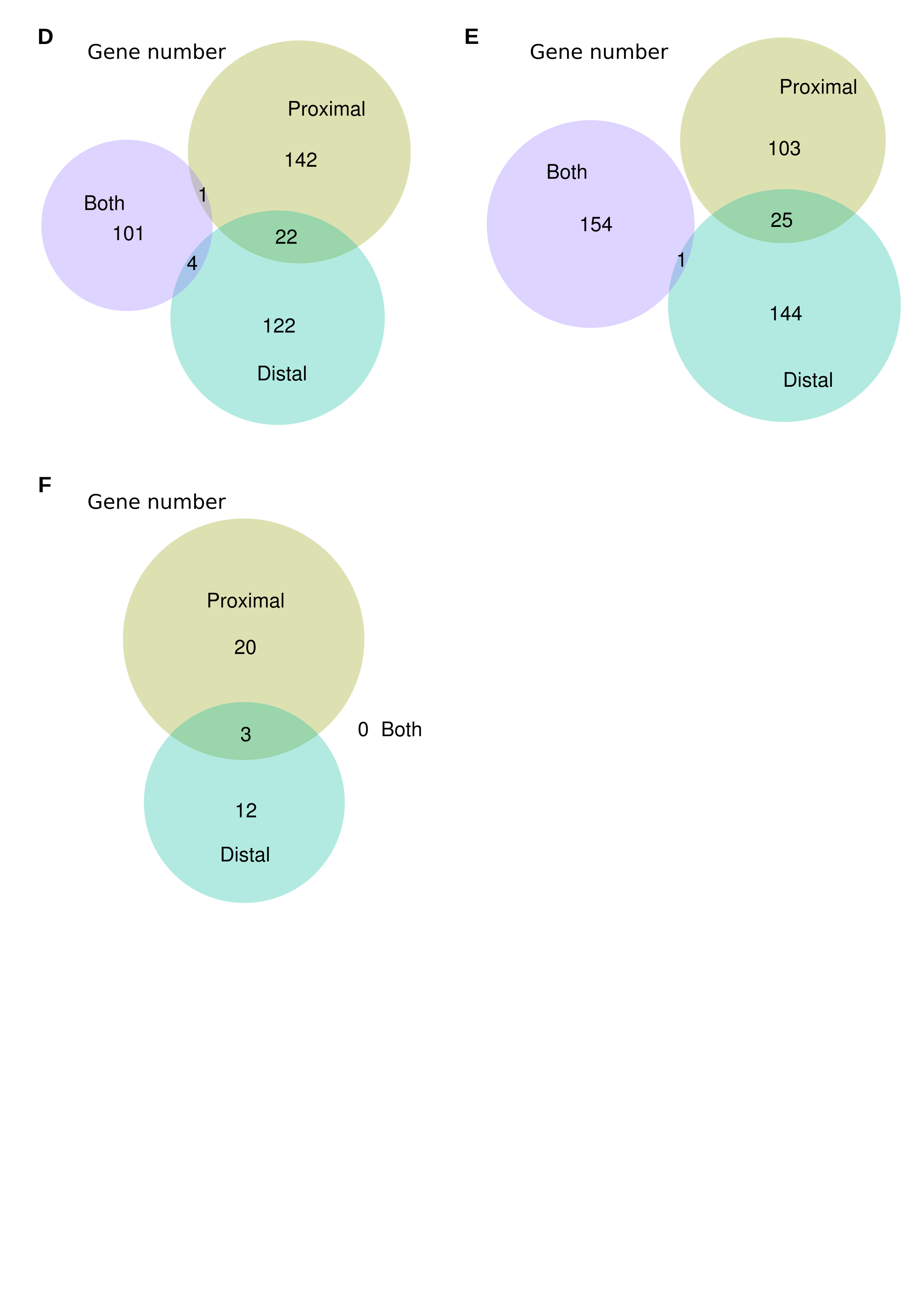

**Figure S2G-H**

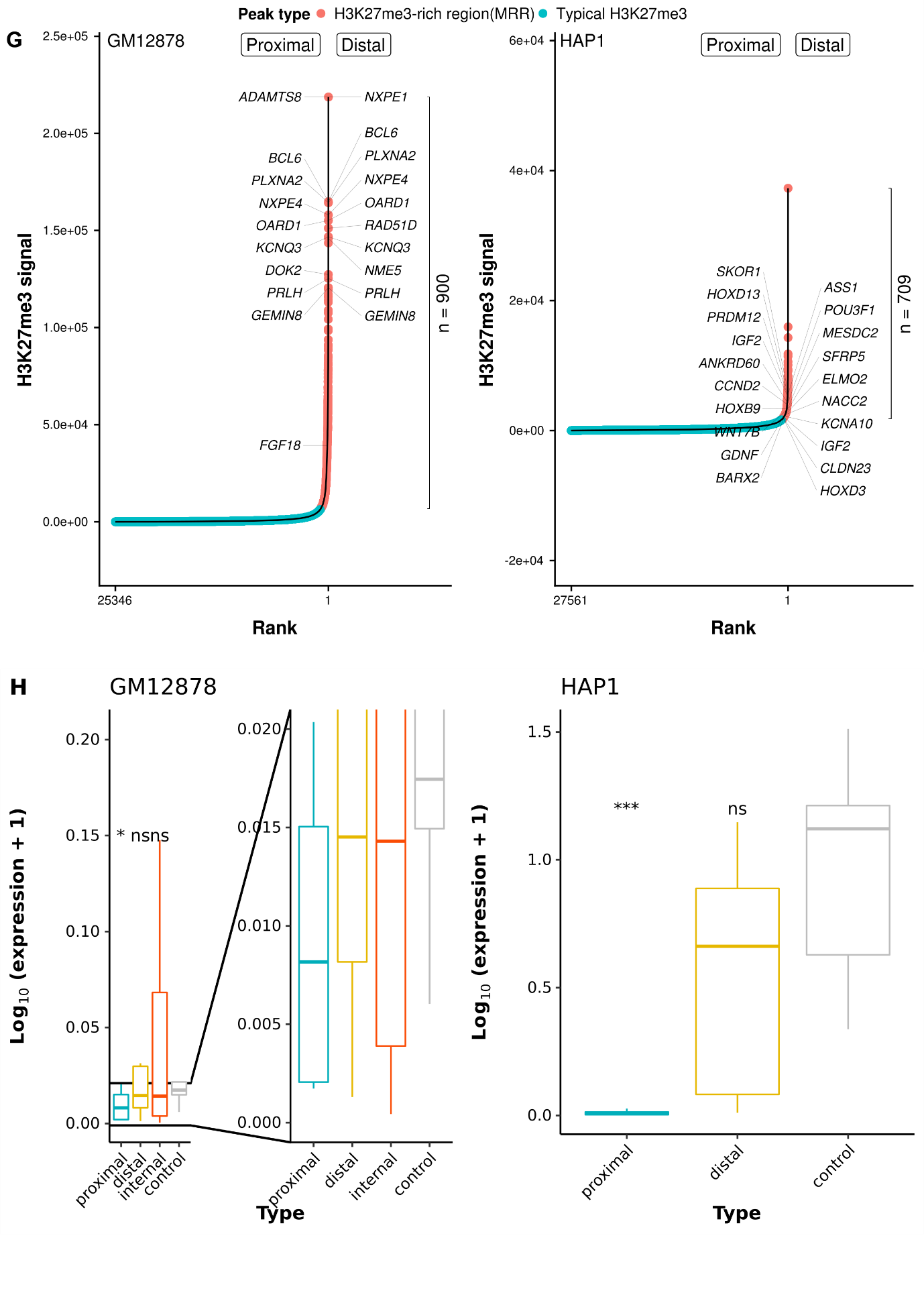

**Figure S2I-M**

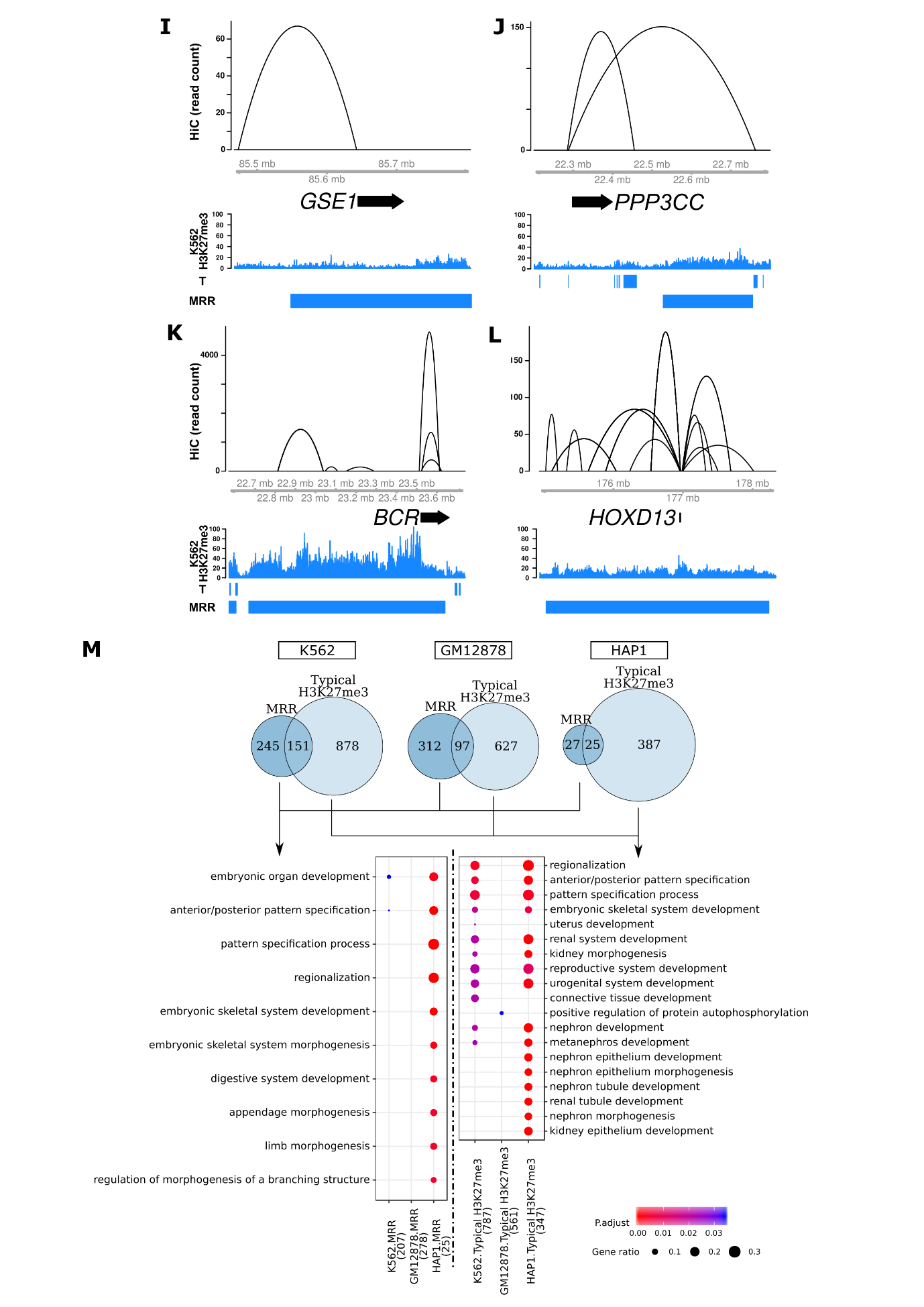

**Figure S2N-P**

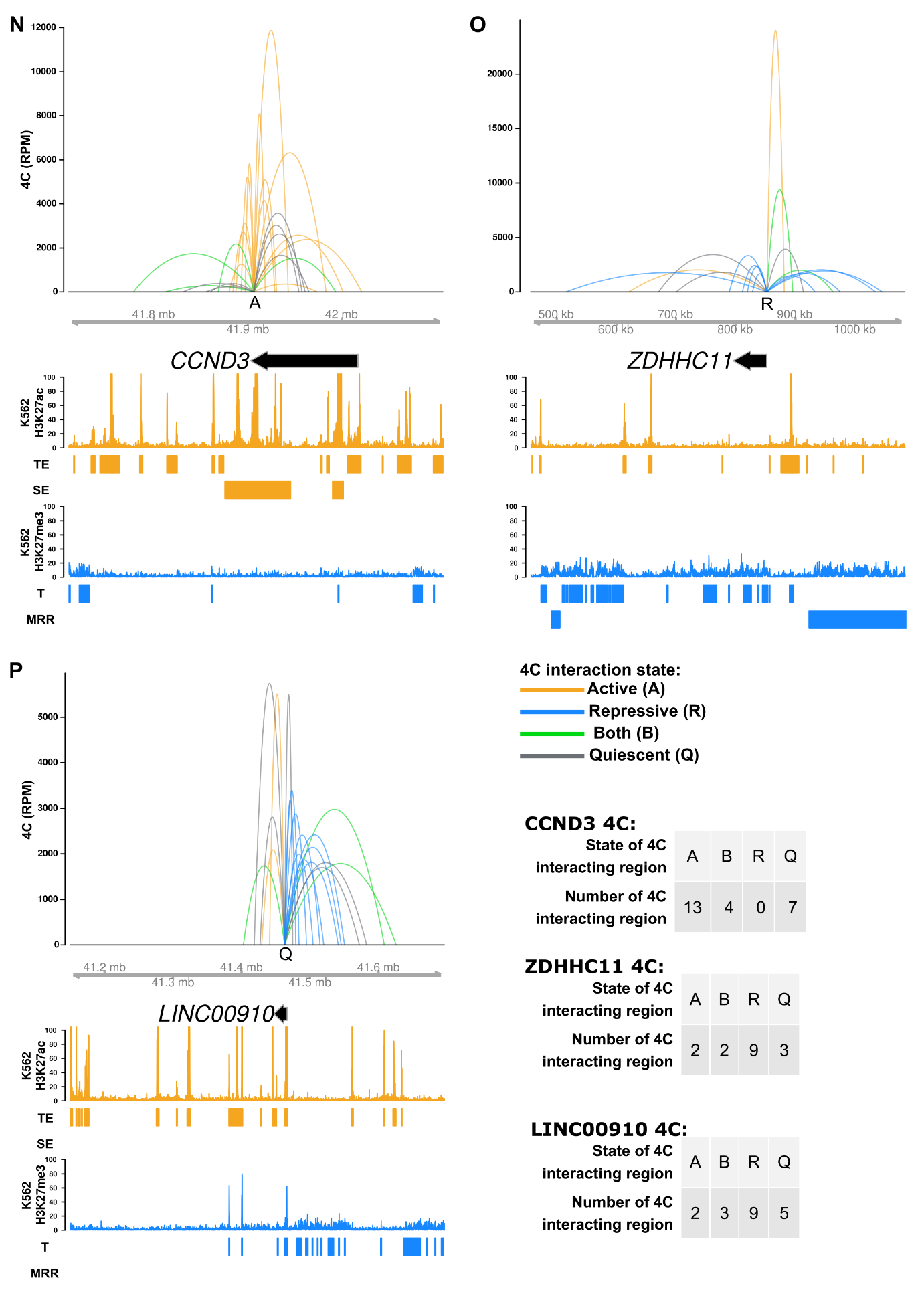

**Figure S2Q**

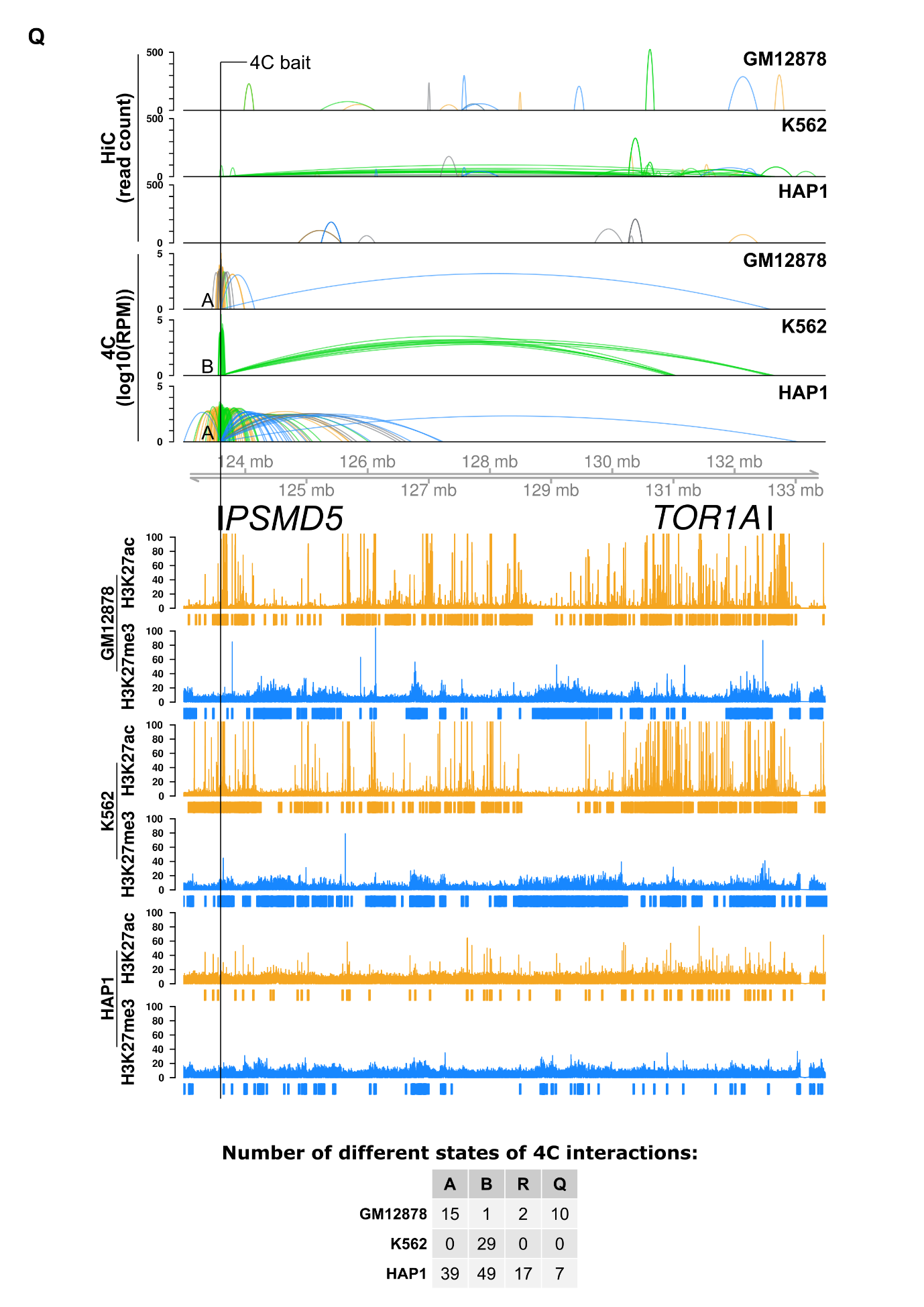

**Figure S2R**

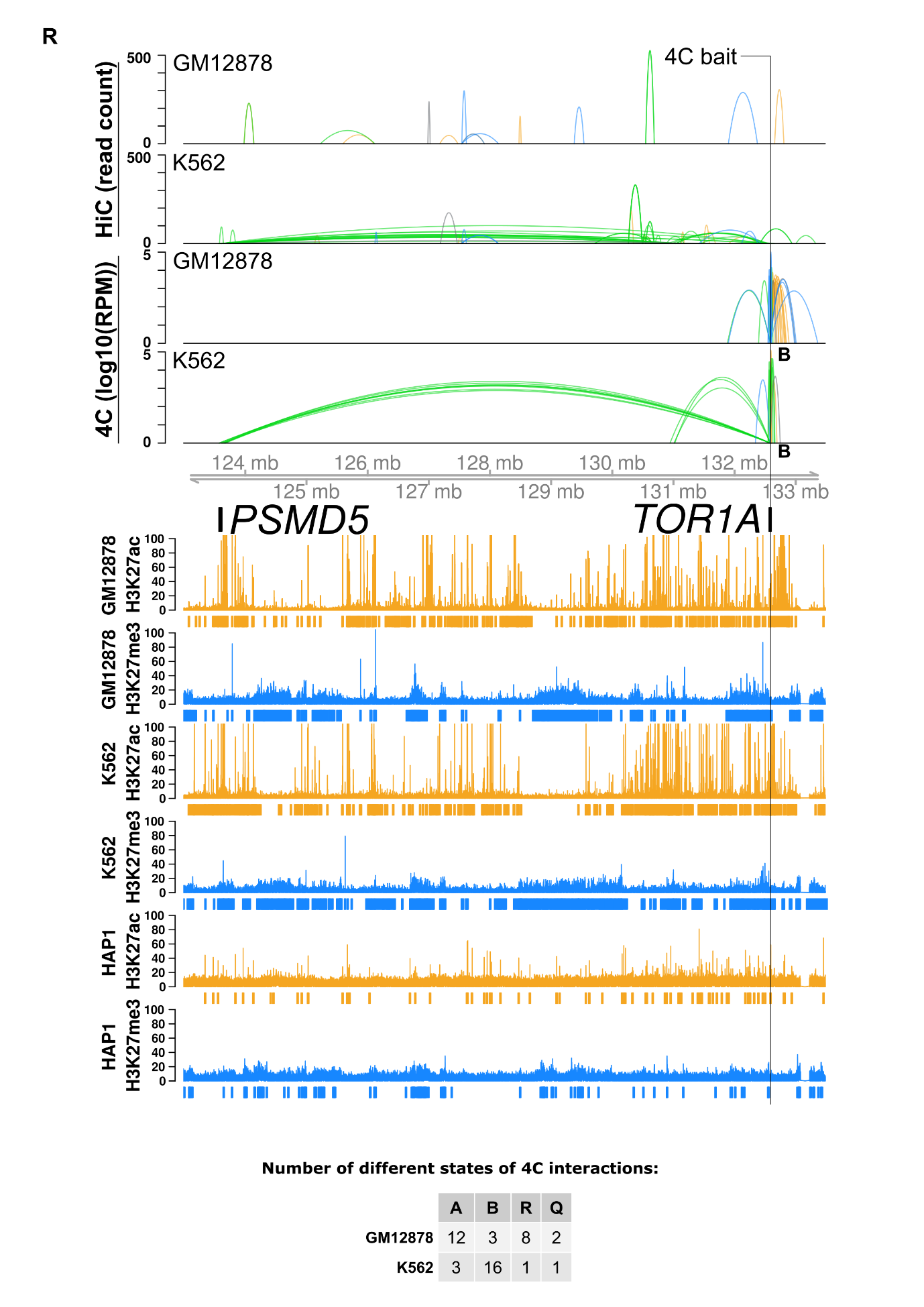

**Figure S3A-C**

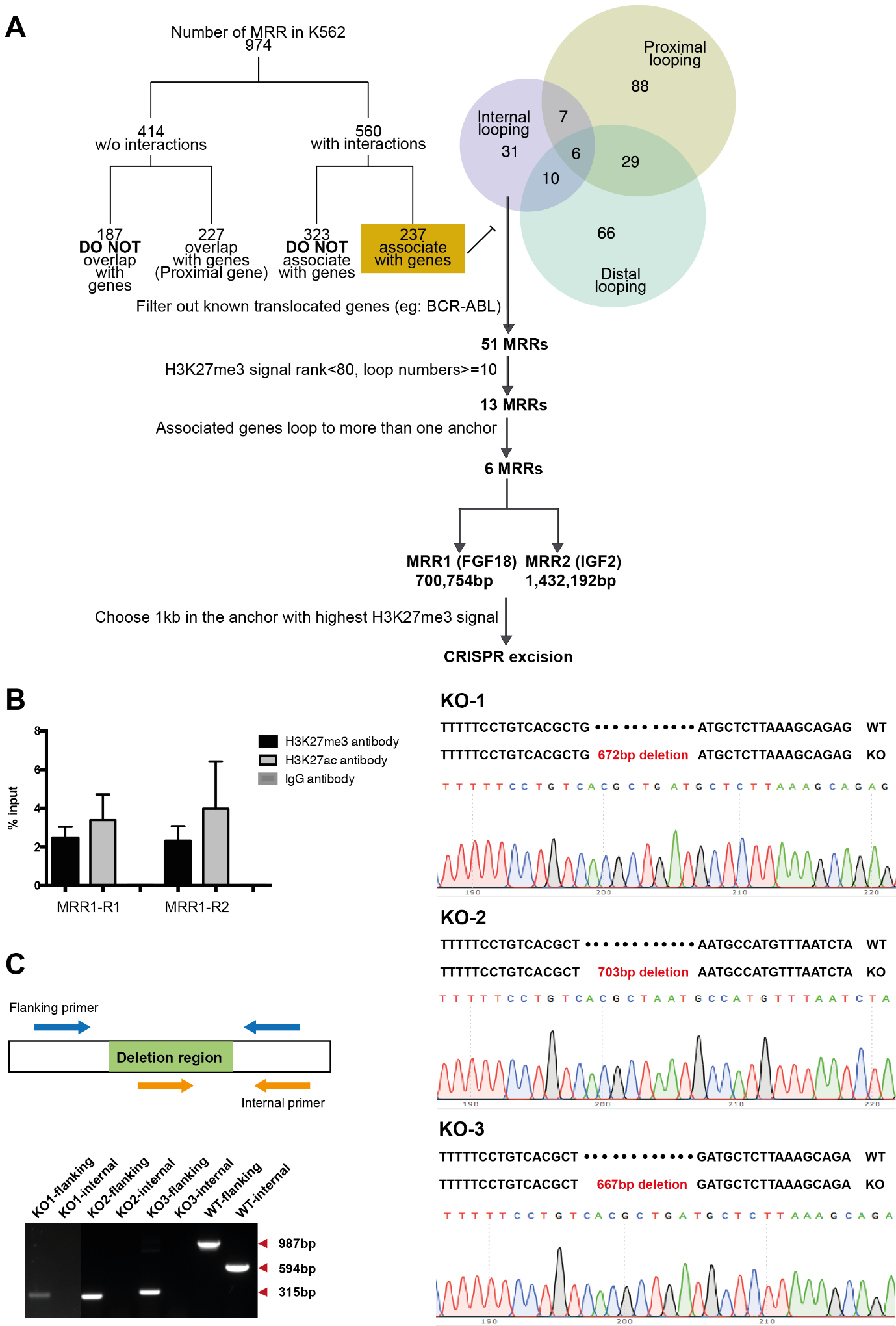

**Figure S3 D-E**

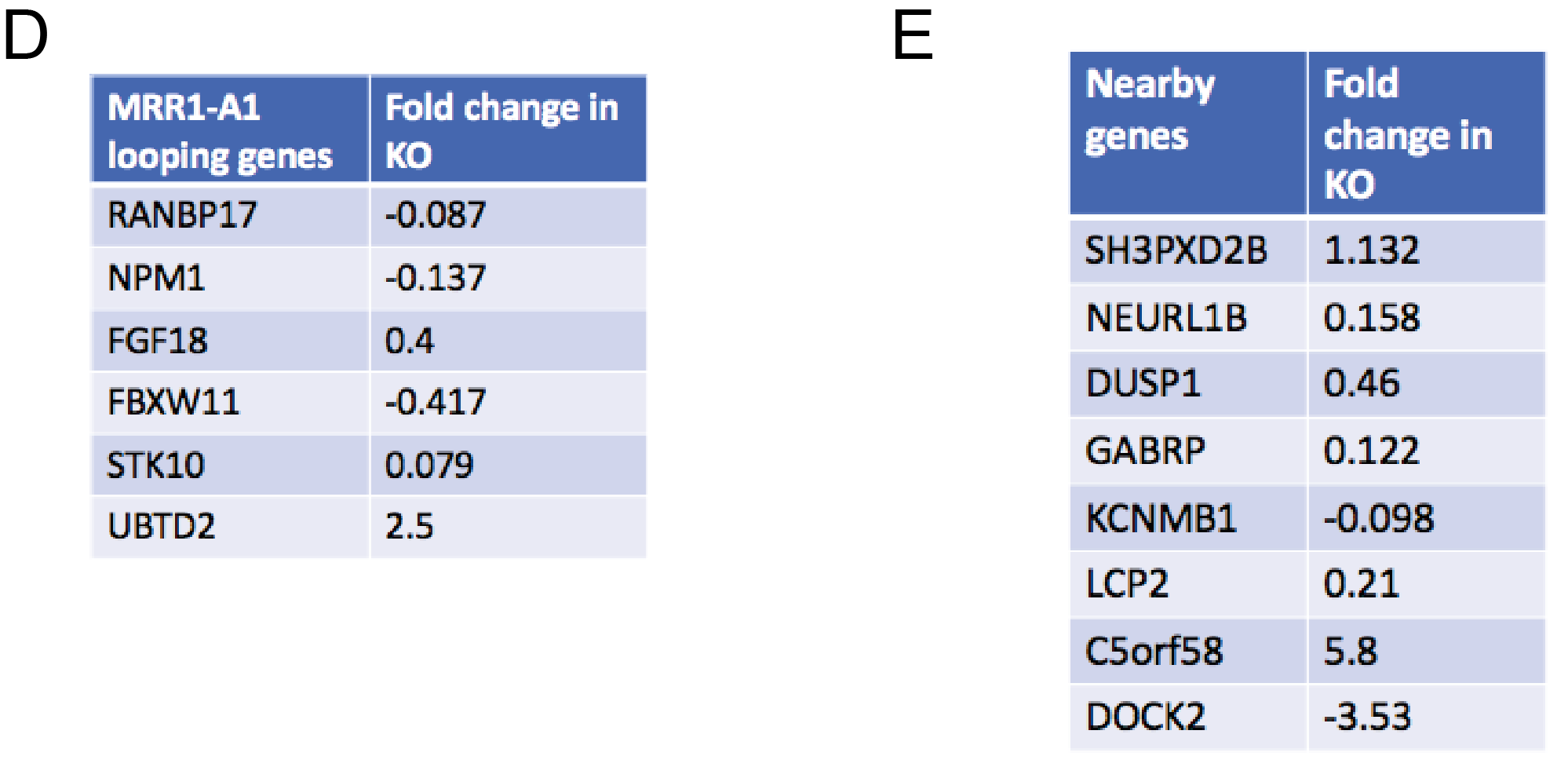

**Figure S4 A-B**
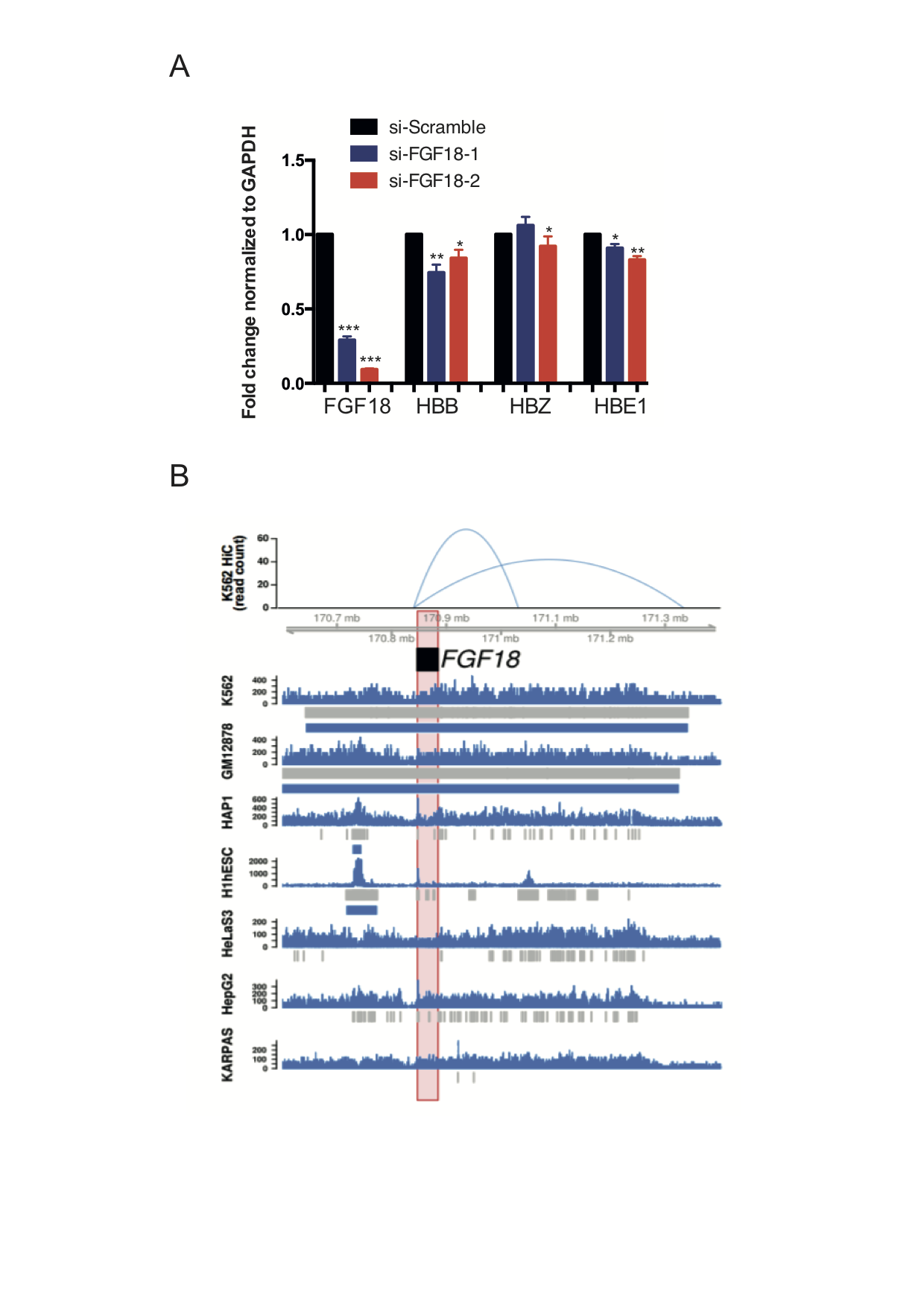

**Figure S5 A-E**

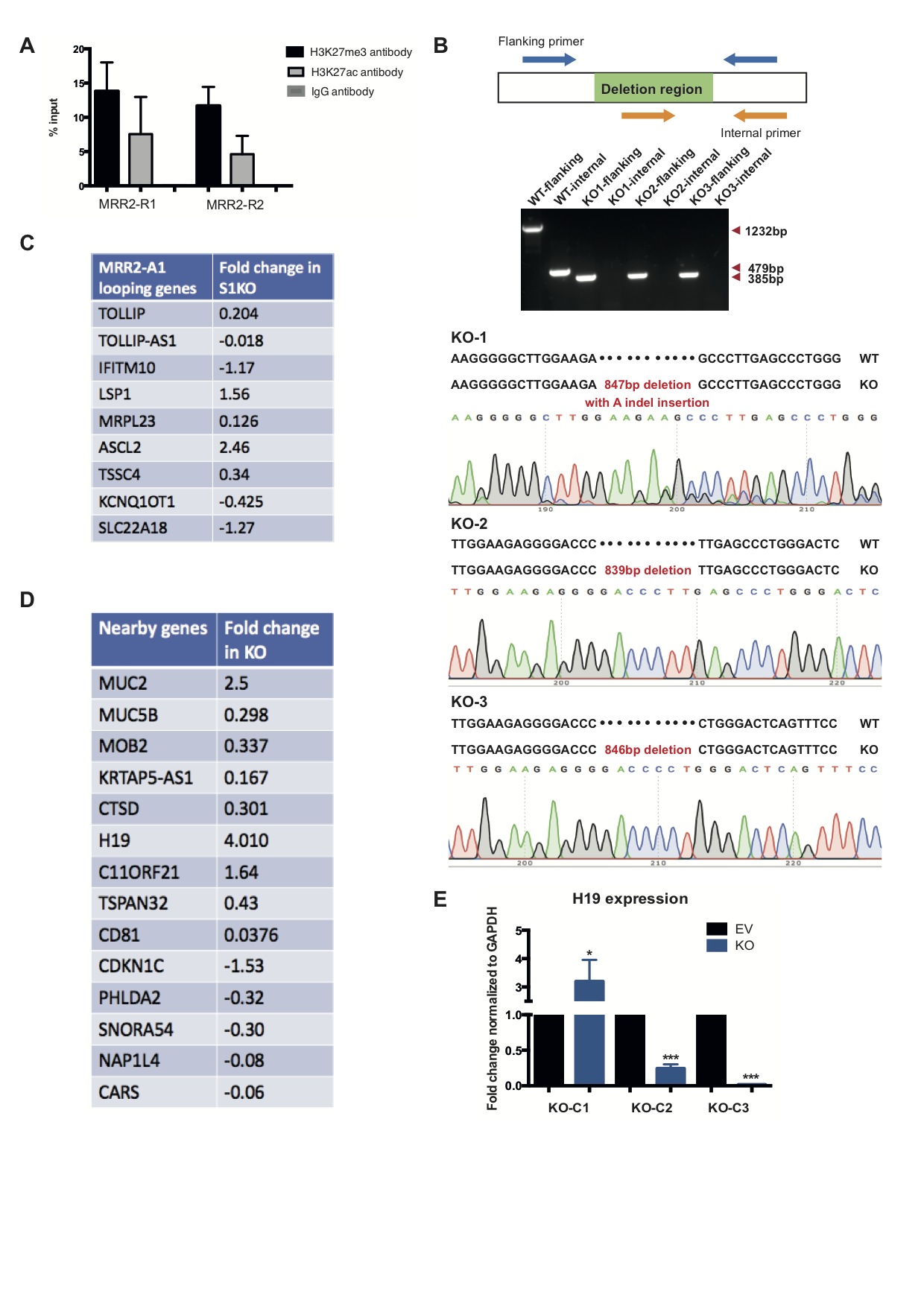

**Figure S5F-I**

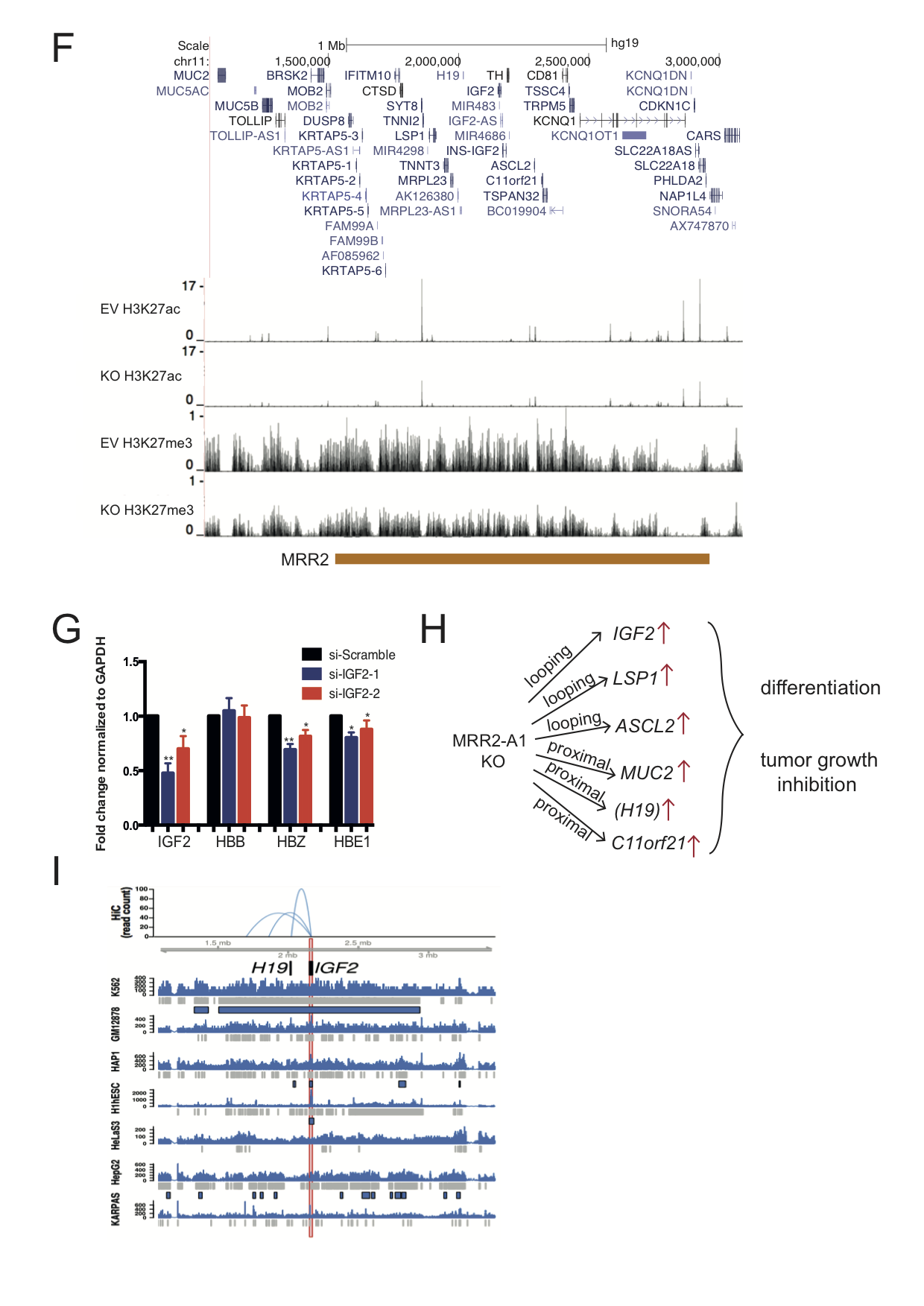

**Figure S6A-D**

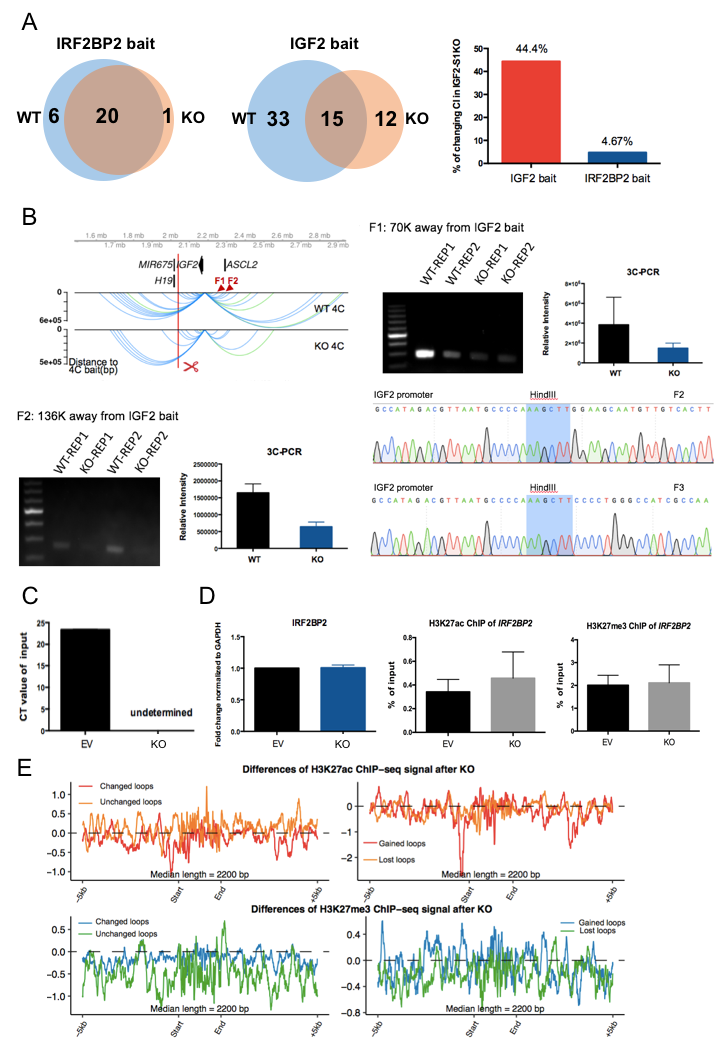

**Figure S7A-D**

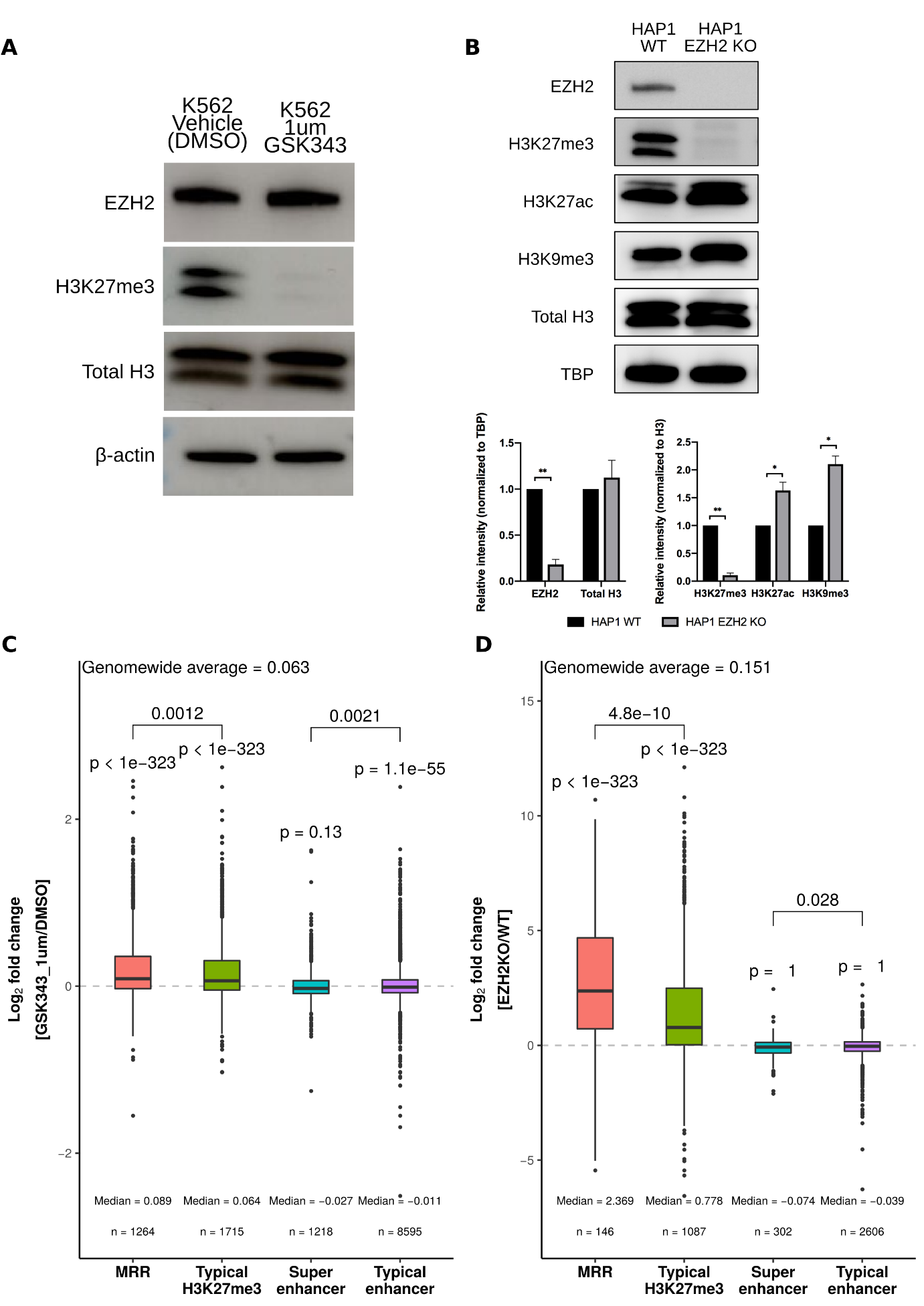

**Figure S7E**

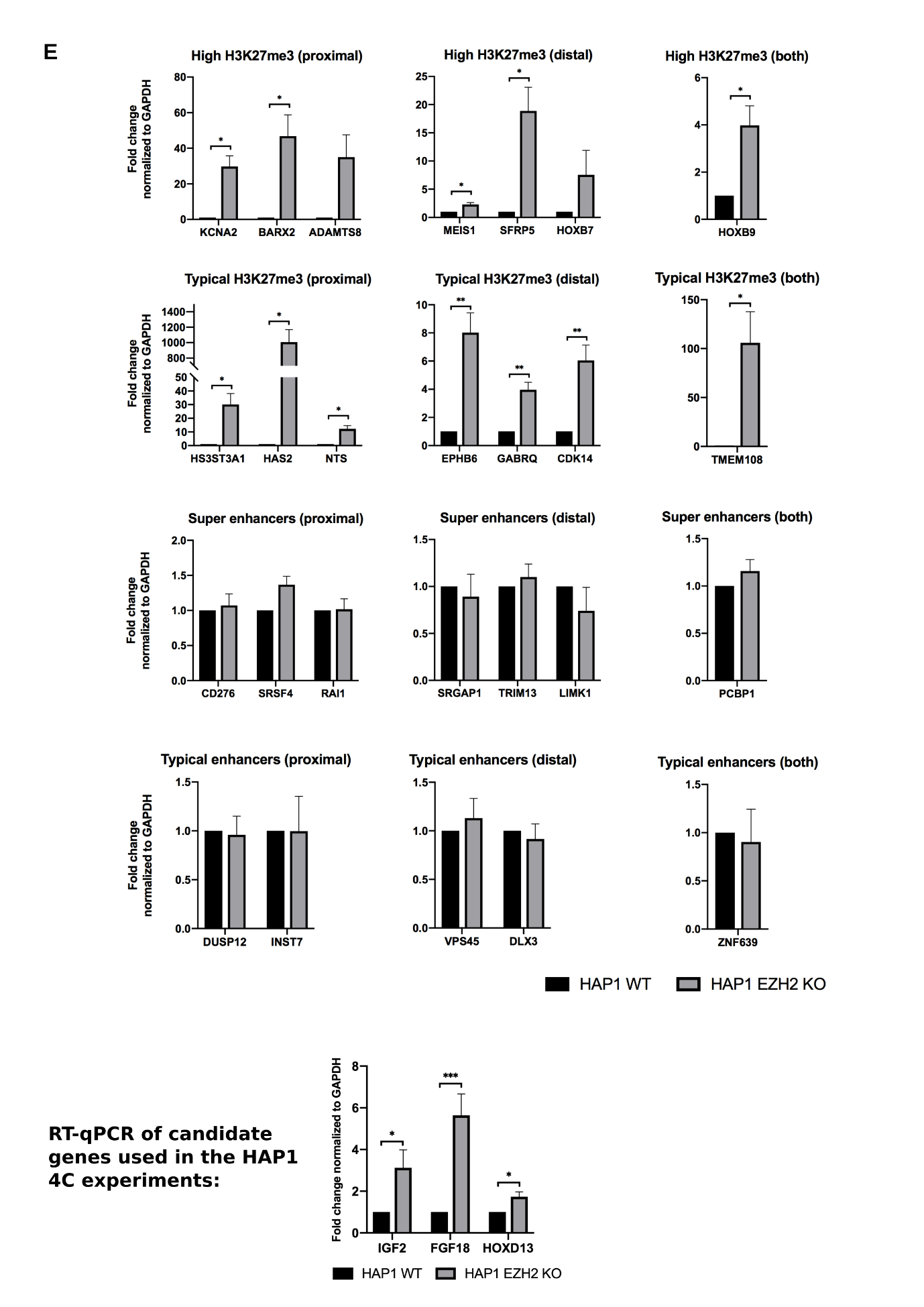

**Figure S7F**

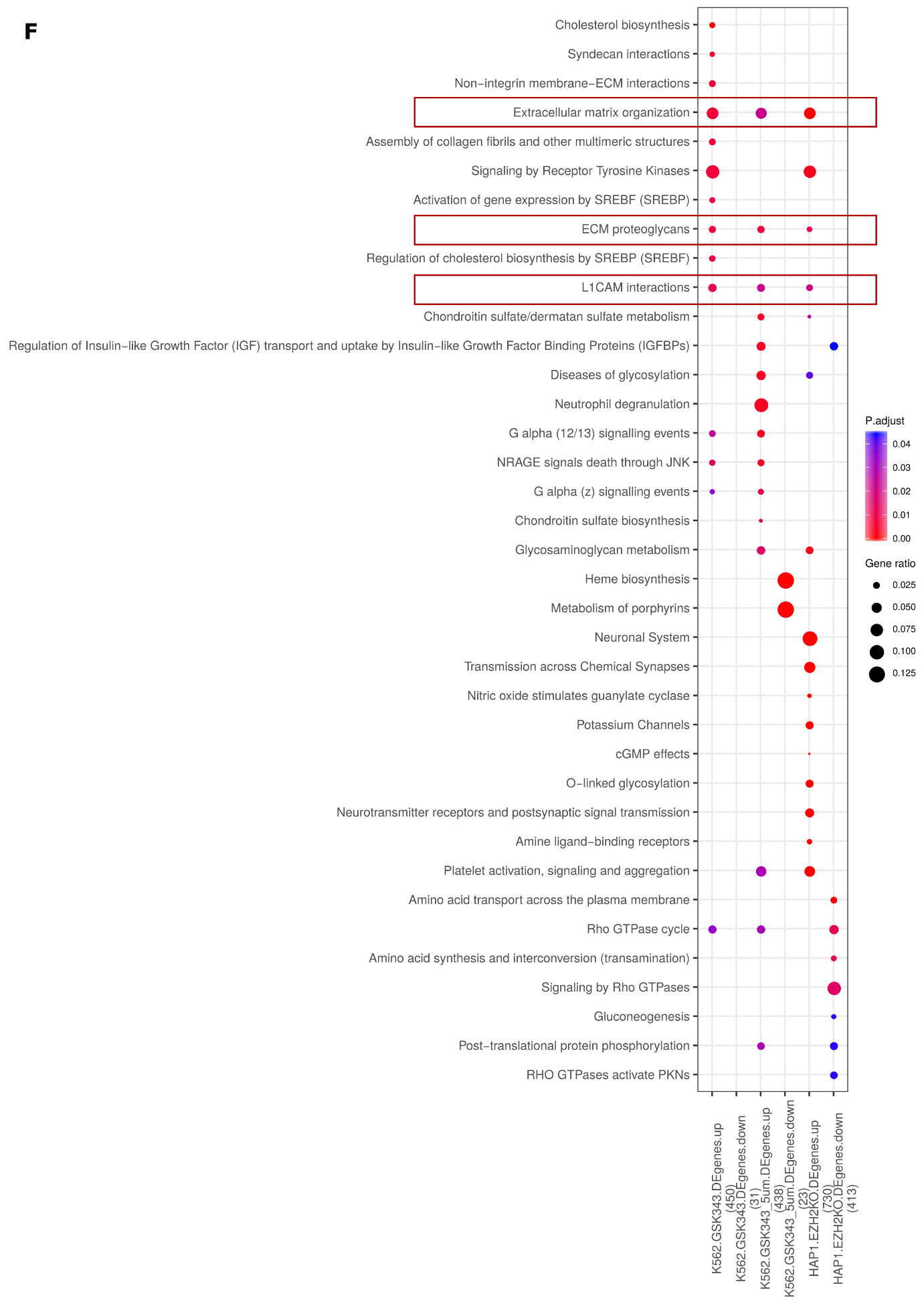

**Figure S7G-I**

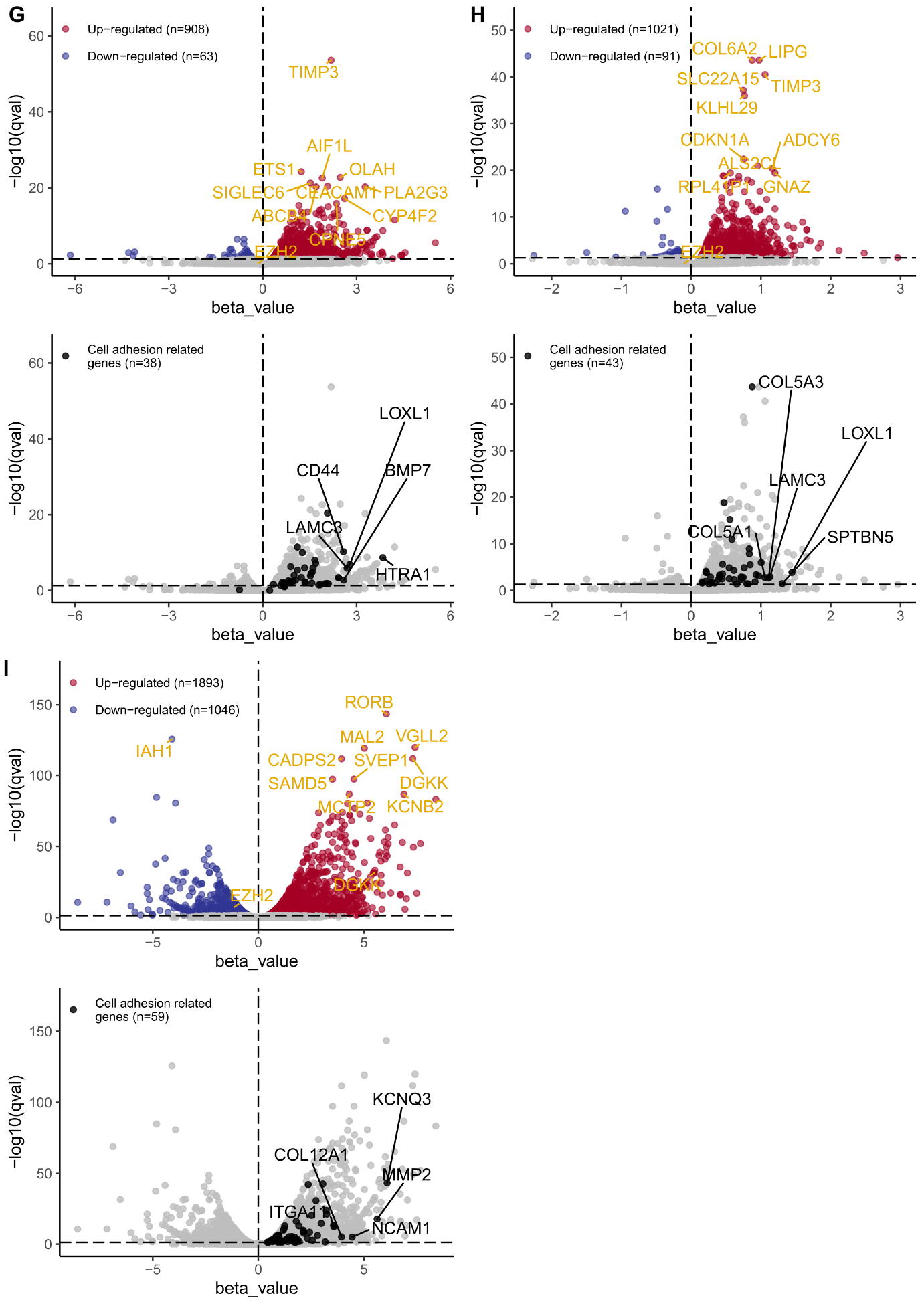

**Figure S7J-K**

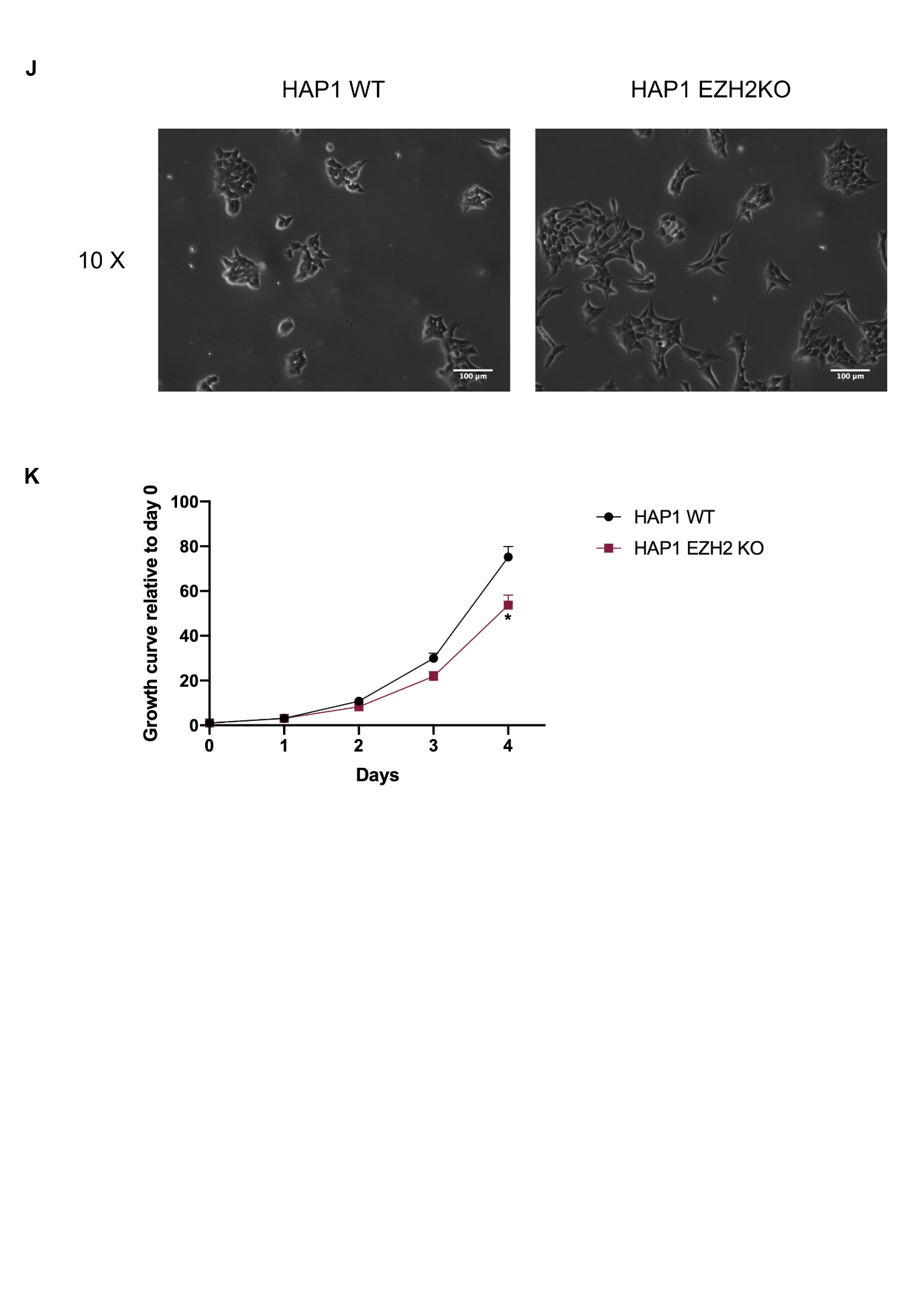

**Figure S7L-M**

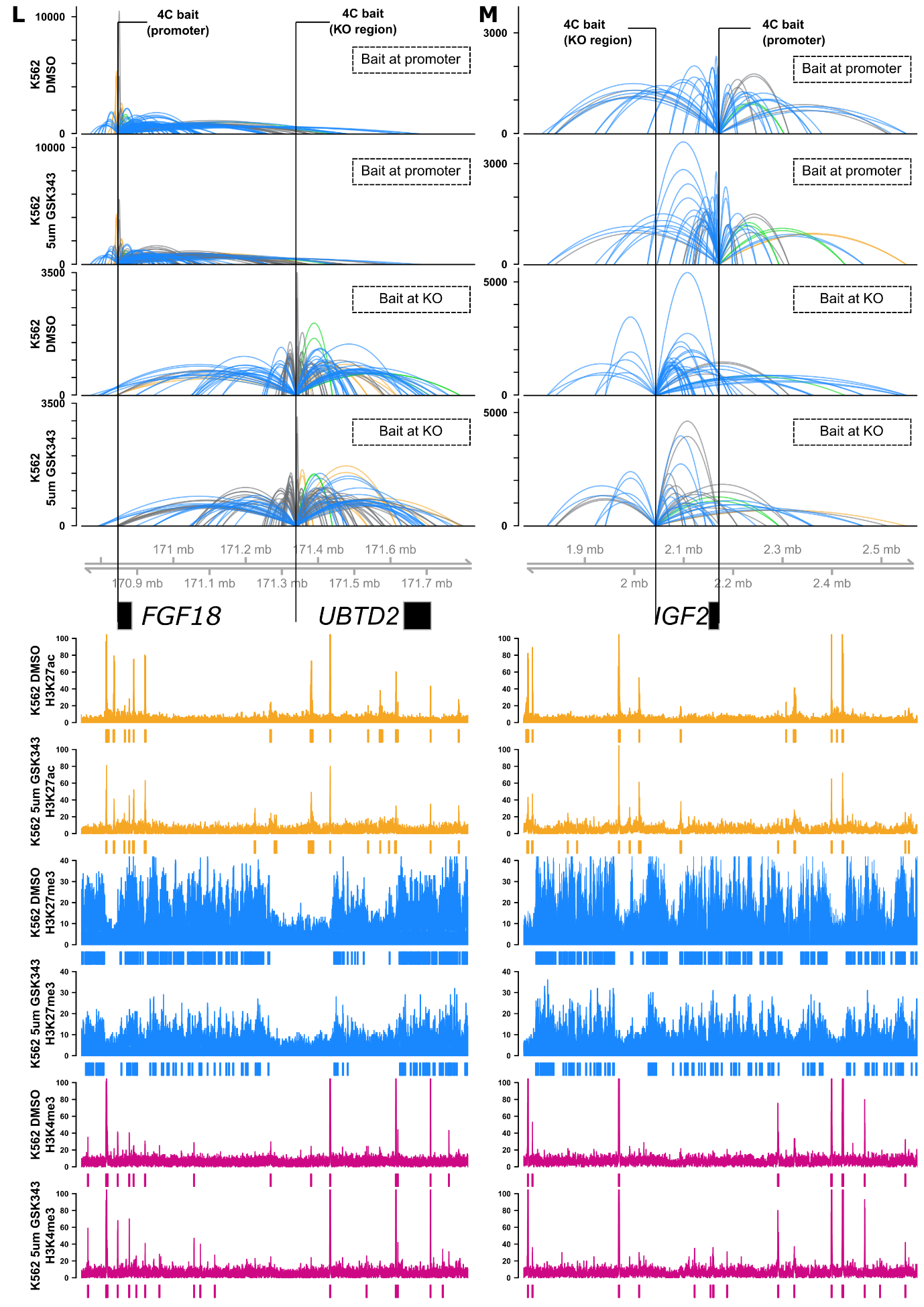

**Figure S7N**

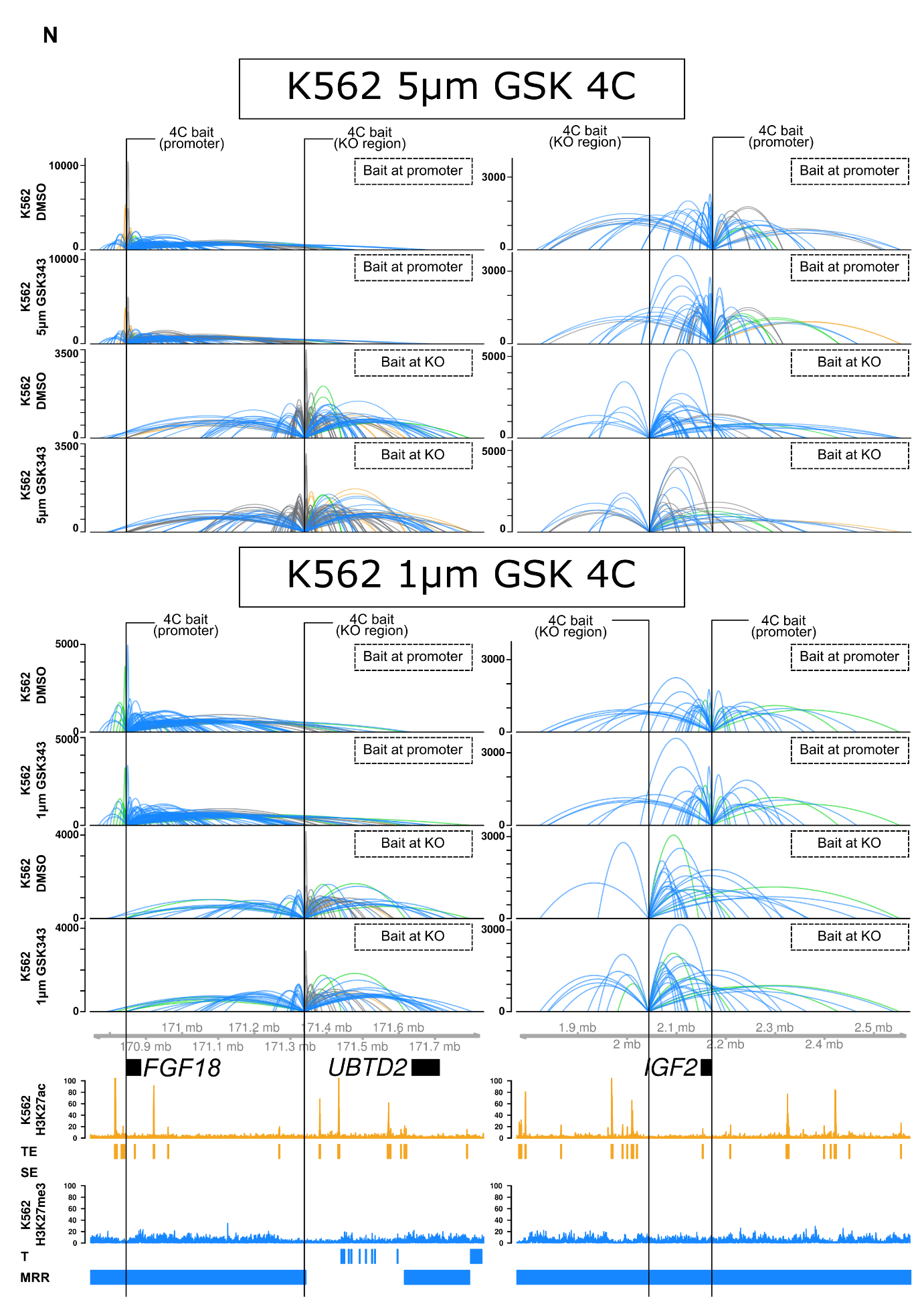

**Figure S7O**

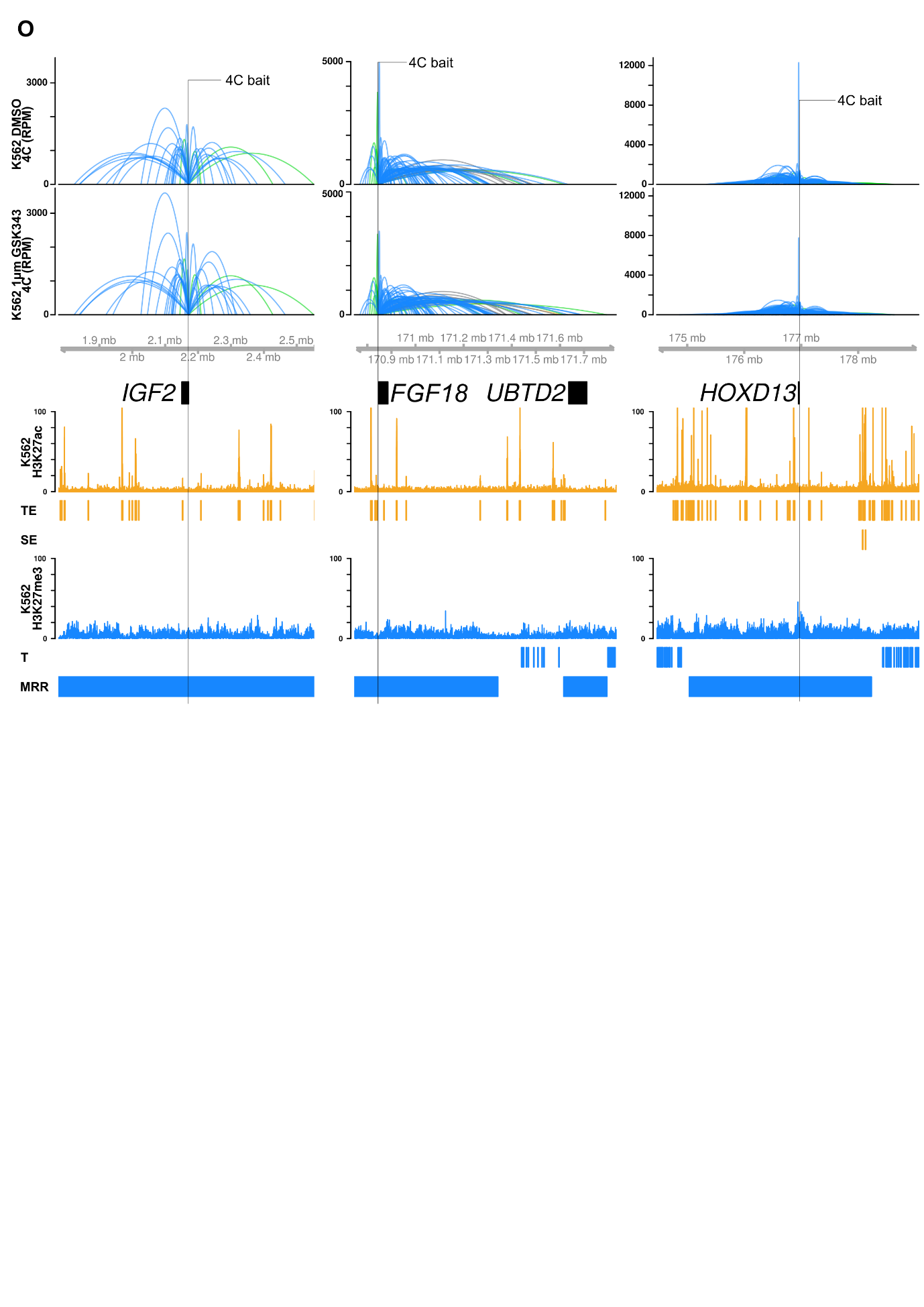

**Figure S7P**

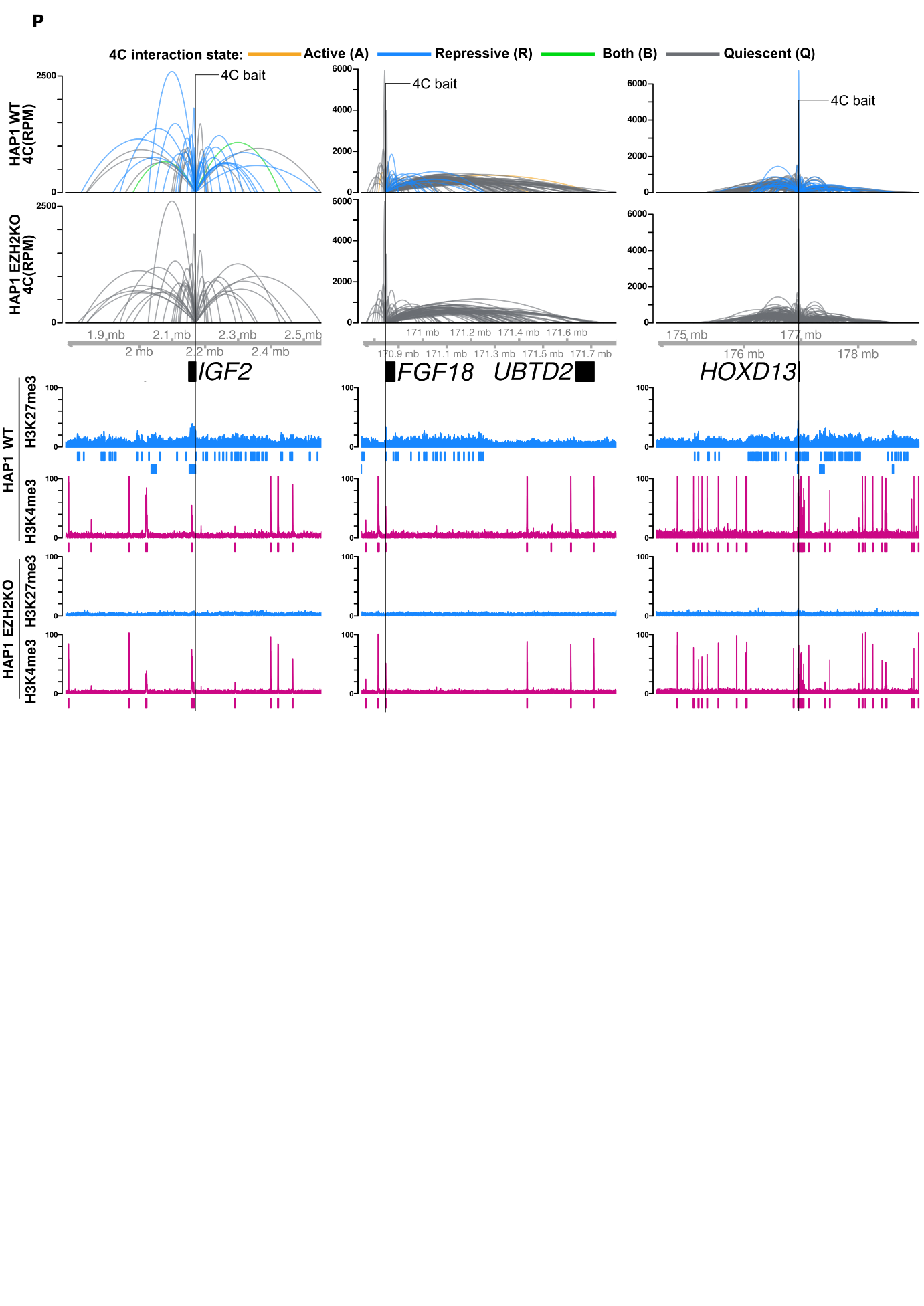

**Figure S7Q**

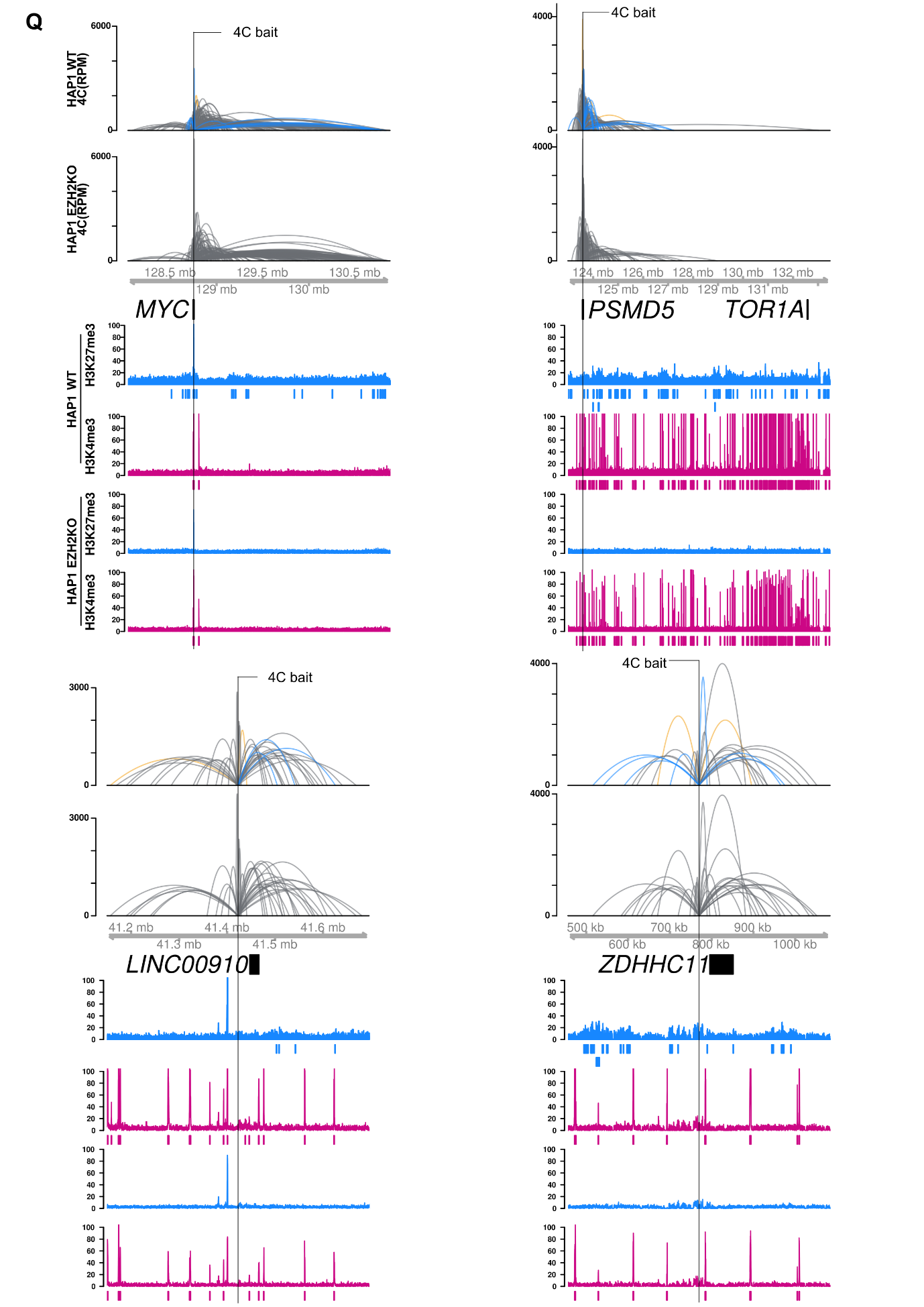

**Figure S7R**

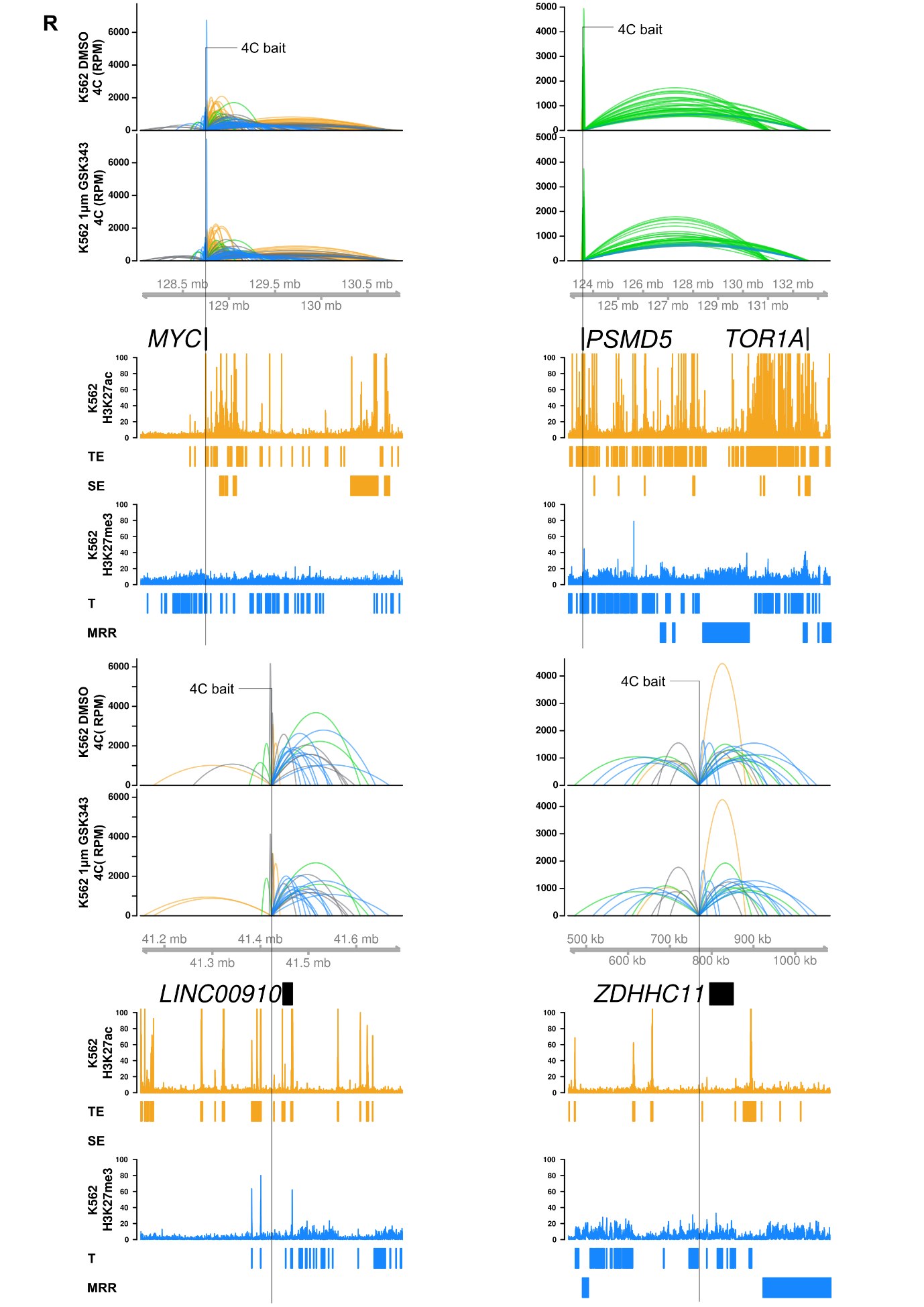

**Figure S8A-D**

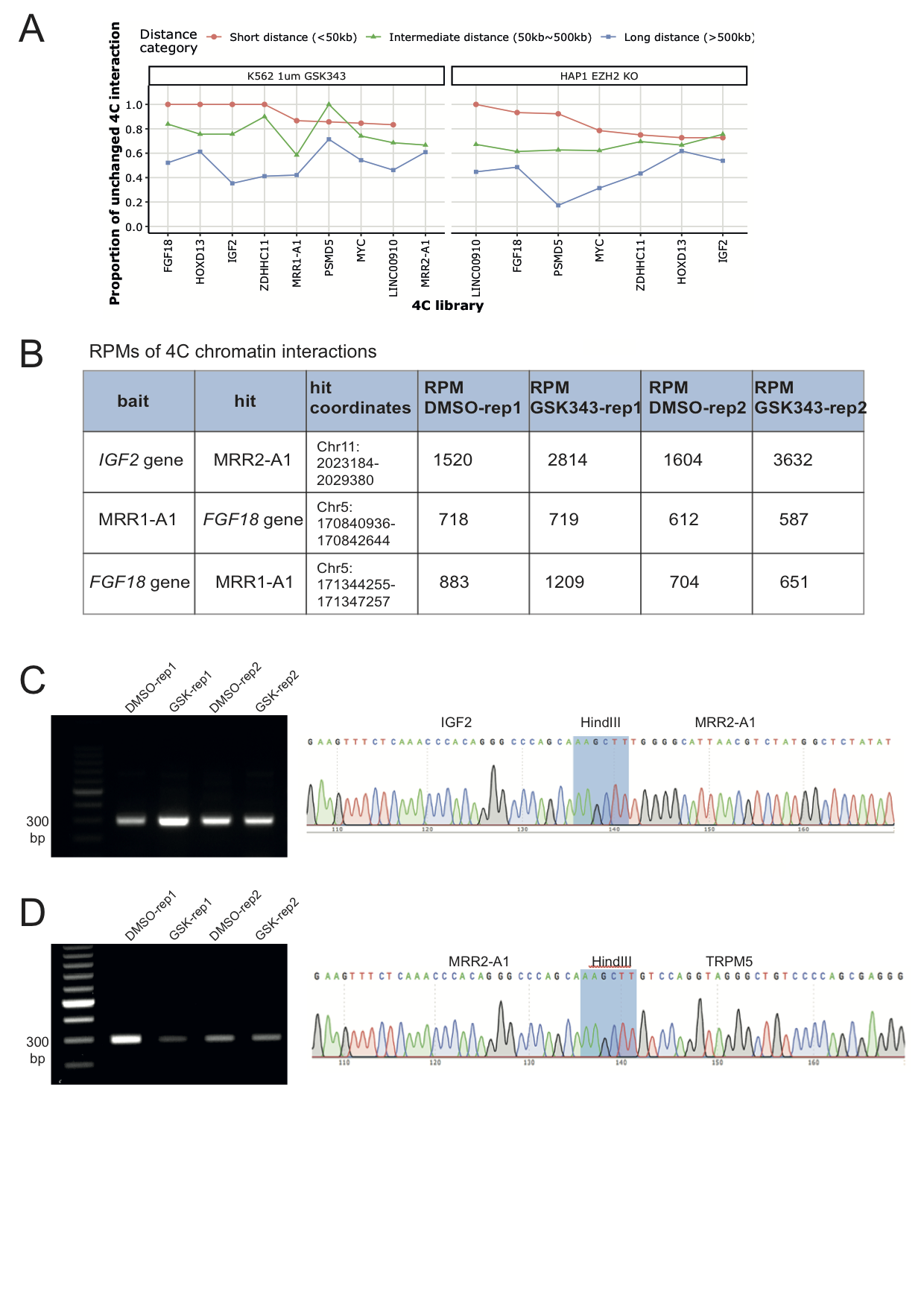

**Figure S9A**

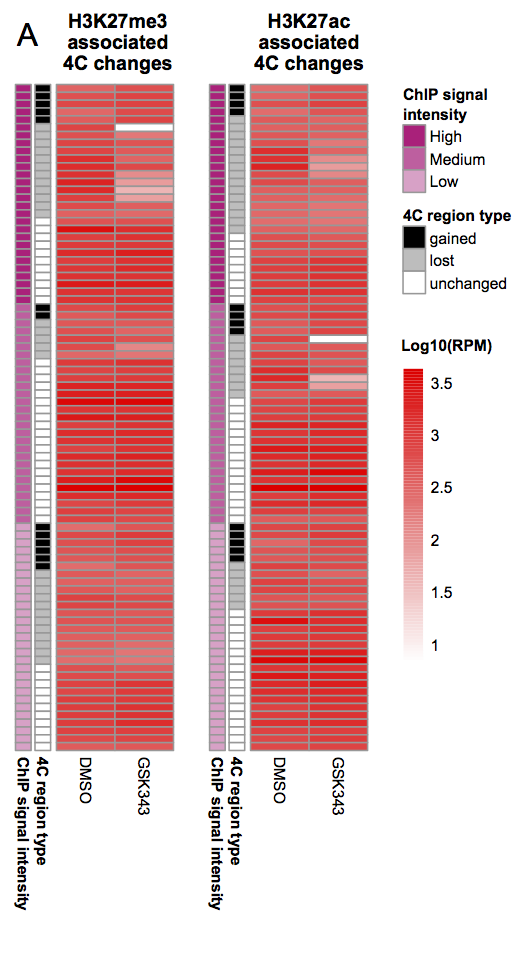

**Figure S9B-D**

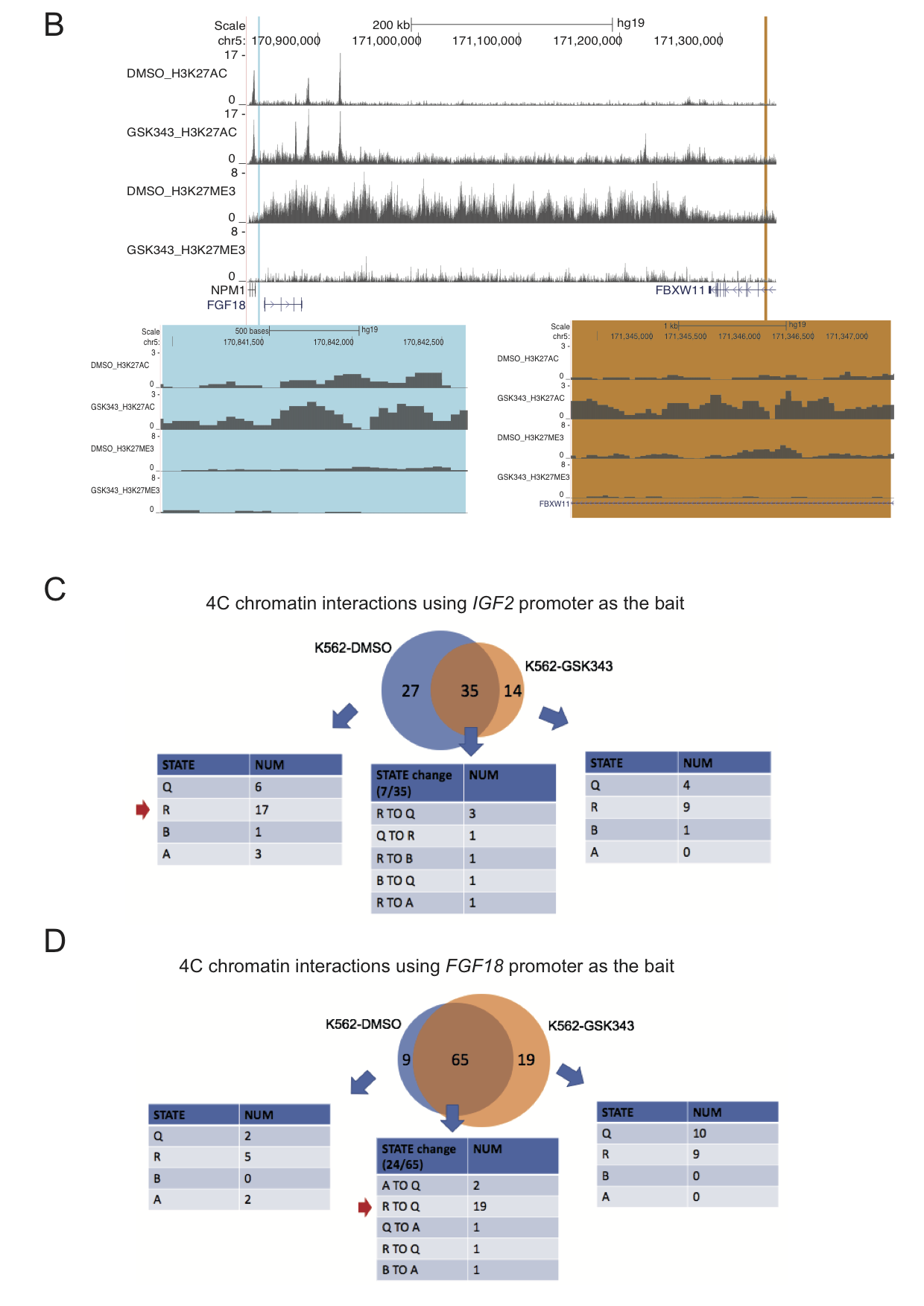
